# sRNA-controlled iron sparing response in Staphylococci

**DOI:** 10.1101/2022.06.26.497478

**Authors:** Rodrigo H. Coronel-Tellez, Mateusz Pospiech, Maxime Barrault, Wenfeng Liu, Valérie Bordeau, Christelle Vasnier, Brice Felden, Bruno Sargueil, Philippe Bouloc

**Author notes:** This work is dedicated to the memory of our colleague Brice Felden who died.

## Abstract

*Staphylococcus aureus*, a human opportunist pathogen, adjusts its metabolism to cope with iron deprivation within the host. We investigated the potential role of small non-coding RNAs (sRNAs) in dictating this process. A single sRNA, named here IsrR, emerged from a competition assay with tagged-mutant libraries as being required during iron starvation. IsrR is iron-repressed and predicted to target mRNAs expressing iron-containing enzymes. Among them, we demonstrated that IsrR down-regulates the translation of mRNAs of enzymes that catalyze anaerobic nitrate respiration. The IsrR sequence reveals three single-stranded C-rich regions (CRRs). Mutational and structural analysis indicated a differential contribution of these CRRs according to targets. We also report that IsrR is required for full lethality of *S. aureus* in a mouse septicemia model, underscoring its role as a major contributor to the iron-sparing response for bacterial survival during infection. IsrR is conserved among staphylococci, but it is not ortholog to the proteobacterial sRNA RyhB, nor to other characterized sRNAs down-regulating mRNAs of iron-containing enzymes. Remarkably, these distinct sRNAs regulate common targets, illustrating that RNA-based regulation provides optimal evolutionary solutions to improve bacterial fitness when iron is scarce.

## INTRODUCTION

Iron is essential for numerous enzymatic processes that use its redox properties, notably in respiration. Despite being one of the most abundant elements on earth, iron bioavailability can be limiting. The mammalian host may prevent pathogen proliferation by restricting access to free iron, creating a nutritional immunity. In parallel, pathogens have developed complex systems for iron assimilation leading to a “battle for iron” (1-4). While iron is required for growth, its excess is toxic. Indeed, ferrous iron decomposes hydrogen peroxide via the Fenton reaction into oxygen radical species that are highly reactive and oxidize a wide range of substrates, including bacterial macromolecules (DNAs, RNAs, proteins, and lipids). Since most living organisms need iron but are sensitive to its excess, strict regulation of iron uptake and iron-containing enzymes is necessary (5,6).

For many bacteria, the main contributor to iron homeostasis is the transcriptional regulator Fur (7). In the presence of iron, Fur acts as a repressor targeting genes involved in iron acquisition, thereby ensuring a feedback regulation to maintain homeostasis of intracellular iron. In *Escherichia coli*, the absence of Fur leads to repression of a subset of genes using iron as a cofactor by an indirect effect. The mystery of this apparent paradox was solved by the discovery of a <u>s</u>mall regulatory <u>RNA</u> (sRNA) repressed by Fur, RyhB (8). In low iron conditions, RyhB, assisted by the RNA chaperone Hfq, accumulates and pairs to mRNAs, preventing the expression of iron-containing proteins and inducing an iron-sparing response (9). RyhB orthologs are present in many species from the *Enterobacteriaceae* family where they contribute to adaptation to low iron conditions (5,10). In *Pseudomonas* species, a major contributor to this adaptation is PrrF, a sRNA down-regulated by Fur contributing to iron homeostasis (11). Of note, the corresponding *ryhB* and *prrF* genes are duplicated in some species. Surprisingly, much less is known concerning Gram-positive bacteria where, so far, RyhB and PrrF orthologs were not found. sRNAs can nevertheless be regulators of the iron-sparing response. In *Bacillus subtilis*, FsrA sRNA is down-regulated by Fur, and with the help of three RNA chaperones, FbpABC, it prevents the expression of iron-containing enzymes (12-14). In *Mycobacterium tuberculosis*, the sRNA MrsI is predicted to be down-regulated by IdeR, a functional homolog of Fur, and to exert a similar iron-sparing response. Interestingly, this bacterium does not possess Hfq (15).

*Staphylococcus aureus* is a human and animal opportunistic pathogen and a leading cause of nosocomial and community-acquired infections (16,17). Its success as a pathogen relies on the expression of numerous virulence factors and its adaptability to various environmental conditions including metal starvation. Indeed, iron limitation in the host is counteracted by alle*via*tion of Fur repression, leading to the production of iron-scavenging siderophores (18). In this condition, *S. aureus* should down-regulate non-essential iron-containing enzymes to preserve iron for vital processes. However, despite significant research efforts devoted to the characterization of this major pathogen, no such system has been characterized to date (18).

As sRNAs are major regulators of iron homeostasis in several species, we designed a selection assay to identify staphylococcal sRNAs that could be involved in this process. We established a collection of *S. aureus* sRNA gene mutants that was used in fitness assays during iron starvation conditions. A single sRNA emerged as being essential for optimum growth when iron becomes scarce. We named it IsrR for ‘iron-<u>s</u>paring response regulator’, an acronym reflecting its function uncovered by the present study. In anaerobic conditions, expression of *isrR* prevented nitrate respiration, a non-essential metabolic pathway involving several iron-containing enzymes. IsrR is required for full lethality in an animal septicemia model of infection.

This study uncovers an iron-sparing response in *S. aureus*, dictated by a regulatory RNA, IsrR, which reallocates iron to essential processes when it is scarce.

## MATERIALS AND METHODS

### Bacterial strains, plasmids, and growth conditions

Bacterial strains used for this study are described in Supplementary Table S1. Most of the work was performed with HG003, a *Staphylococcus aureus* model strain widely used for regulation studies (19). Gene annotations refer to NCTC8325 nomenclature (file CP00025.1) retrieved from Genbank and Aureowiki (20). Plasmids were engineered by Gibson assembly (21) in *Escherichia coli* IM08B (22) as described (Supplementary Table S2), using the indicated appropriate primers (Supplementary Table S3) for PCR amplification. Plasmids expressing sGFP were constructed in MG1655Z1 *pcnB*::Km, a strain that decreases ColE1 plasmid copy number and therefore reduces toxic effects produced by high amounts of certain proteins in *E. coli*. MG1655Z1 *pcnB*::Km was constructed by the introduction of the *pcnB*::Km allele (23) in MG1655Z1 (24) by P1-mediated transduction.

Plasmids were verified by DNA sequencing and transferred into HG003, 8325-4, RN4220, or their derivatives. Chromosomal mutants (deletions and insertions) were either reported (25) or constructed for this study (Supplementary Table S1) using pIMAY derivatives as described (25), except for the *fur*::*tet* allele which was transferred from MJH010 (26) to HG003 and HG003 Δ*isrR*::tag135 by phage-mediated transduction.

Staphylococcal strains were routinely grown in Brain Heart Infusion (BHI) broth at 37°C aerobically or anaerobically under anaerobic conditions (5% H2, 5% CO_2_, and 90% N_2_) in an anaerobic chamber (Jacomex). Δ*fur S. aureus* derivatives in anaerobic conditions were grown in Tryptic Soy Broth (TSB). *E. coli* strains were grown aerobically in Luria-Bertani (LB) broth at 37°C. Antibiotics were added to media as needed: ampicillin 100 μg/ml and chloramphenicol 20 μg/ml for *E. coli*; chloramphenicol 5 μg/ml and kanamycin 60 μg/ml for *S. aureus*. Iron-depleted media was obtained by the addition of either DIP (2,2’-dipyridyl) 1.25 mM, unless stated otherwise; or EDDHA (ethylenediamine-N,N’-bis(2-hydroxyphenylacetic acid)) 0.7 mM and incubated for 30 min prior to the addition of bacteria.

### Fitness assay

Mutants with altered fitness in the presence of iron chelators were identified and analyzed using a reported strategy (25) with three independent libraries containing 48 mutants (Supplementary Table S4). The libraries were grown at 37°C in BHI, BHI DIP 1.25 mM, and BHI EDDHA 0.7 mM for 3 days. Overnight cultures were diluted 1000 times in fresh pre-warmed medium. Samples were withdrawn at OD_600_ 1 and overnight as indicated (Figure 1A).

**Figure 1.**
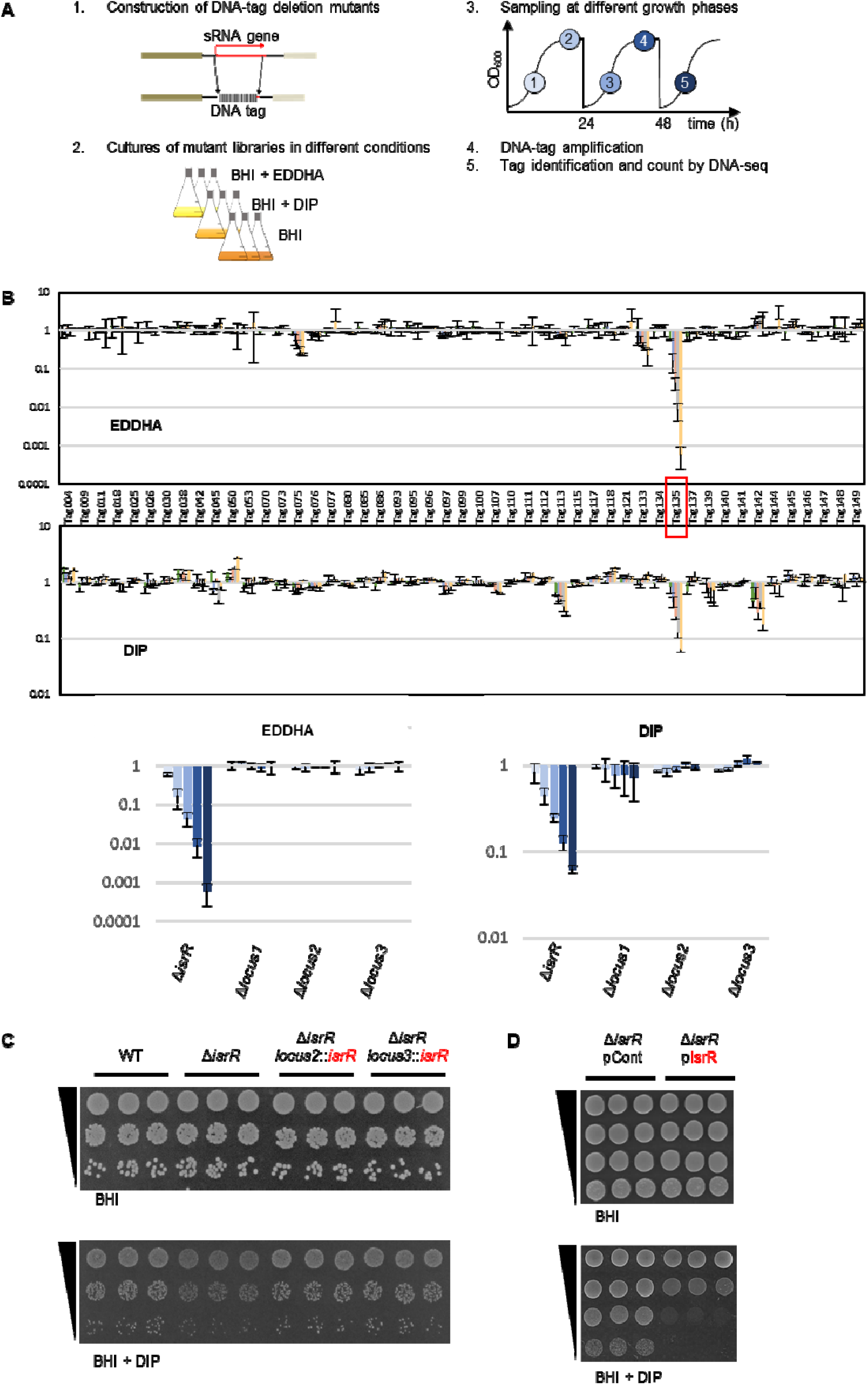
*S. aureus* IsrR sRNA is required for optimal growth when iron is scarce. (A) Experimental protocol scheme to select mutants with altered fitness in media containing iron chelators. (B) Evolution of mutant proportions in libraries (for composition, see Supplementary Table S4) grown in the presence of EDDHA 0.7mM or DIP 1.25mM normalized to the same libraries grown in the absence of iron chelators. Error bars indicate the standard deviation from three independent libraries. Upper parts; data with the complete 48 mutant libraries. Tag135 corresponding to the *isrR* mutant is highlighted by a red box. For Tag/mutant correspondence, see Supplementary Table S1. Note that Tag145 and Tag149 are in a single strain, which corresponds to a double mutant (Δ*sprX2* Δ*sprX1*). Lower parts, selected data (enlargement): Δ*isrR* and three other tagged regions are shown. Loci 1, 2, and 3 correspond to tag insertions in non-transcribed regions expected not to affect bacterial growth. Histogram color code corresponds to sampling time color code from Figure 1A. (C) Δ*isrR* growth defect in iron-depleted media is complemented by an *isrR* ectopic chromosomal copy. For strain constructions see Supplementary Table S1 and Supplementary Figure S2A. Plating efficiency of indicated strains on BHI medium without (upper panels) or with (lower panels) DIP 1.25mM. Three independent biological clones are shown for each strain. For results with EDDHA, see Supplementary Figure S2C. (D) Multicopy *isrR* is toxic in iron-depleted media. Experiments were performed as for Figure 1C. pCont, pCN38; pIsrR, pCN38-IsrR. For strain and plasmid constructions see Supplementary Tables S1 and S2.

### Spot tests

Strain plating efficiency was visualized by spot tests. 5 μL of ten-fold serial dilutions of overnight cultures were spotted on plates containing either BHI, BHI DIP 1.25 mM or BHI EDDHA 0.7mM. Plates were supplement with chloramphenicol 5 μg/ml when strains with plasmids were tested. BHI plates were incubated overnight at 37°C while BHI DIP and BHI EDDHA were incubated for 24h.

### Northern Blots

Total RNA preparations and Northern blots were performed as previously described (27). 10 μg of total RNA samples were separated either by agarose (1.3%) or acrylamide (8%) gel electrophoresis. Membranes were probed with primers ^32^P-labelled with [alpha-P32]Deoxycytidine 5’-triphosphate (dCTP) using Megaprime DNA labelling system (GE Healthcare) (for primers, see Supplementary Table S3) and scanned using the Amersham Typhoon imager.

### Rifampicin assay

Bacteria were cultured overnight in BHI. Then, 200 μl of cultures were transferred into 60 ml of fresh BHI or BHI supplemented with DIP 1.4 mM and grown at 37°C. At OD_600_ 1.5 (t0), rifampicin was added to a final concentration of 200 μg/ml. Six ml of cultures were harvested at t0 and 1, 3, 5, 10, and 20 min after the addition of rifampicin and transferred to tubes in liquid nitrogen to stop bacterial growth. RNA was extracted as described (27) and Northern blots were performed as described above. tmRNA was used as RNA loading control.

### Biocomputing analysis

DNA-seq data from fitness experiments were analyzed using a reported pipeline (25). *isrR* orthologs were identified thanks to the sRNA homolog finder GLASSgo (28) with *isrR* (174nt) as input using the default parameters. Fur boxes were detected using the motif finder FIMO (29) with *isrR* sequences retrieved from GLASSgo (option including 100nt upstream *isrR* sequences) and *S. aureus* NCTC8325 Fur consensus motifs from RegPrecise (30). *IsrR* secondary structure was modelled with LocARNA (31) with 18 IsrR orthologs with non-identical sequences using default parameters. Putative *IsrR* targets were found by CopraRNA (32) set with default parameters and *isrR* orthologs from *S. aureus* NCTC8325 (NC_007795), *S. epidermidis* RP62A (NC_002976), *S. lugdunensis* HKU09-01 (NC_013893), *Staphylococcus pseudintermedius* HKU10-03 (NC_014925), *Staphylococcus pseudintermedius* ED99 (NC_017568), *Staphylococcus xylosus* (NZ_CP008724), *Staphylococcus warneri* SG1 (NC_020164), *Staphylococcus pasteuri* SP1 (NC_022737), *Staphylococcus hyicus* (NZ_CP008747), *Staphylococcus xylosus* (NZ_CP007208), *Staphylococcus argenteus* (NC_016941), *Staphylococcus carnosus* TM300 (NC_012121), *Staphylococcus cohnii* (NZ_LT963440), *Staphylococcus succinus* (NZ_CP018199), *Staphylococcus agnetis* (NZ_CP009623), *Staphylococcus piscifermentans* (NZ_LT906447), *Staphylococcus stepanovicii* (NZ_LT906462), *Staphylococcus equorum* (NZ_CP013980), *Staphylococcus nepalensis* (NZ_CP017460), *Staphylococcus lutrae* (NZ_CP020773), *Staphylococcus muscae* (NZ_LT906464), *Staphylococcus simulans* (NZ_CP023497).

### Probability calculation

For a set of N objects (number of *S. aureus* mRNAs = 2889) with m different objects (number of mRNA expressing Fe-S containing proteins = 32 (estimated number in *S. aureus* [Table S6, (33,34)], the probability of drawing n objects (23 best targets proposed by CopraRNA) and to have among k differentiated objects (mRNA expressing Fe-S containing proteins = 7) is given the following equation 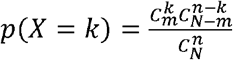 were C represents a combination operator (https://www.dcode.fr/picking-probabilities).

### Electrophoretic mobility shift assay

RNAs were transcribed from PCR products (for primers, see Supplementary Table S3) using T7 RNA polymerase and purified by ethanol precipitation. Wild-type alleles were generated from HG003 genomic DNA and mutated alleles from gBlock DNAs (Integrated DNA Technologies) containing the desired mutations. Gel-shift assays were performed as described (35) with the following modifications: RNAs were denaturated in 50 mM Tris pH 7.5, 50 mM NaCl for 2 min at 80°C, followed by refolding for 20 min at 25°C after adding MgCl2 at final concentration of 5 mM. The binding reactions were performed in 50 mM Tris-HCl (pH 7.5), 50 mM NaCl, 5 mM MgCl2 for 30 min at 37°C. About 0.25 pmoles of labeled IsrR or IsrRmut were incubated with various concentrations of *fdhA* mRNA or *fdhA*mut mRNA. Samples were supplemented with 10% glycerol and were loaded on a native 8% polyacrylamide gel containing 5% glycerol. Gels were dried and visualized by autoradiography.

### Fur boxes reporter assay

Fluorescent tests to evaluate the contribution of Fur boxes were performed with 8325-4, a ‘pigment-less’ strain (36) containing either pP_isrR_::*gfp*, pP_isrR1_::*gfp*, pP_isrR2_::*gfp* or pP_isrR1&2_::*gfp* (Supplementary Table S2). Overnight cultures were diluted 100 times, sonicated for 30 seconds, and fixated with ethanol 70% and PBS 1X. Fluorescence was detected using a Partec Cube 6 flow cytometer with ≈ 500,000 cells for each measure, with 488 and 532 nm wavelength filters for excitation and emission, respectively.

### Quantification of nitrite

*S. aureus* cultures were grown aerobically overnight, diluted 1000 times in fresh media and grown anaerobically overnight. They were then diluted to OD_600_ 0.01, after 2 hours of incubation at 37°C, NaNO_3_ 20 mM was added. 150 or 240 min after, samples were recovered and nitrite in supernatant was determined using the Griess Reagent System (Promega) according to the manufacturer’s instructions. OD_600_ and OD_540_ were determined with a microtiter plate reader (CLARIOstar).

### 5’/3’ RACE mapping

5’/3’ RACE was performed as previously described (37) with some modifications. Total RNA (6 μg) from HG003 Δ*fur* was treated with TAP (Epicentre) for 1 h at 37°C. After phenol/chloroform extraction followed by ethanol precipitation, RNA was circularized using T4 RNA ligase (Thermo Scientific) overnight at 16°C. Circularized RNA was once more extracted with phenol/chloroform, ethanol precipitated, and reverse transcribed using primer 2728 (Supplementary Table S3). A first PCR across the 5’/3’ junction was performed using primers 2729/2730, followed by a nested PCR with primers 2731/2732. The PCR products were cloned using the pJET1.2/blunt system (Thermo Scientific) within *E. coli* and 20 samples were sequenced.

### sRNA activity reporter assay

The principle of a reporter assay for sRNA activity was described for Enterobacteria (38) and Gram-positive bacteria (39). The effect of sRNAs on targets is determined *via* the quantification of leader reporter fusions. The latter results from in-frame cloning of *gfp* downstream of 5’UTRs and first codons of target genes. sRNAs and reporter genes are both on plasmids. We developed a similar system to test *IsrR* activity against its putative substrates.

Plasmids driving constitutive expression of isrR (pRMC2ΔR-IsrR) and its derivatives with CRR deletions (pRMC2ΔR-IsrRΔCRR1, pRMC2ΔR-IsrRΔCRR2, and pRMC2ΔR-IsrRΔCRR3 expressing *isrR*ΔCRR1, *isrR*ΔCRR2, and *isrR*ΔCRR3, respectively) were constructed and introduced in HG003 Δ*isrR* (Supplementary Tables S1 and S2). These strains were subsequently transformed with p5’FdhA-GFP, p5’NarG-GFP, p5’NasD-GFP, and p5’GltB2-GFP reporting the activity of *IsrR* on *fdhA, narG, nasD*, and *gltB2* mRNAs, respectively (Supplementary Tables S1 and S2). The latter plasmids express the complete interaction region with IsrR of each mRNA target predicted by IntaRNA plus ten additional codons in frame with the sGFP coding sequence. Each gene fusion is under the control of the *sarA* promoter. Of note, the four reporter plasmids were constructed in MG1655Z1 *pcnB*::Km. Fluorescence on solid medium was visualized from overnight cultures that were streaked on plates supplemented with chloramphenicol 5 μg/ml and kanamycin 60 μg/ml. Fluorescence was detected using Amersham Typhoon scanner at 488 nm and 525 nm wavelengths for excitation and emission, respectively. Fluorescence in liquid was visualized from overnight cultures diluted 1000-fold in TSB supplemented with chloramphenicol 5 μg/ml and kanamycin 60 μg/ml, and grown in microtiter plates. Fluorescence was measured after six hours of growth using a microtiter plate reader (CLARIOstar), normalized to OD_600_, and set at 1 for the strains with the control plasmid (pRMC2ΔR).

### SHAPE experiments

Synthetic genes placing *isrR* and, portions of *fdhA, narG, nasD*, and *gltB2* under the transcriptional control of the T7 RNA polymerase were constructed by PCR on HG003 genomic DNA using primers indicated in Supplementary Table S3. The resulting synthetic genes were *in vitro* transcribed using T7 RNA polymerase (40). RNA integrity and folding homogeneity were assessed by agarose gel electrophoresis.

RNAs were probed in the presence and absence of 5 mM Mg^2+^ using methyl-7-nitroisatoic anhydride (1M7) and modifications were revealed by selective 2’-hydroxyl acylation and analyzed by primer extension (SHAPE) as described (41,42) with slight modifications. Briefly, 12 pmoles of RNA (*isrR, fdhA, narG, nasD* and *gltB2*) were diluted in 64 μl of water and denatured for 5 min at 85°C. Then, 16 μl of pre-warmed (37°C) folding buffer (final conc. 40 mM HEPES pH 7.5, 150 mM KCl with/without 5 mM MgCl_2_) were added, and the solution was cooled down to room temperature for 7 min. Samples were then incubated for 5 min in a dry bath set at 37°C.

Similarly, when investigating interactions, RNAs were refolded in separate tubes (12 pmoles of IsrR and 60 pmoles of target RNA or the opposite) as described above, then mixed and incubated for 20 min at 37°C.

Refolded RNAs were split in two and either 1M7 (final concentration of 4 mM) or DMSO was added and incubated at 37°C for 5 min. Probed RNAs were precipitated in ethanol in the presence of ammonium acetate 0.5 M, ethanol, and 20 μg of glycogen. The pellets were washed twice with 70% ethanol, air-dried, and resuspended in 10 μl of water.

Modifications were revealed by elongating fluorescent primers (WellRed D2 or D4 fluorophore from Sigma, for sequences, see Supplementary Table S3) using MMLV reverse transcriptase RNase H Minus (Promega). Purified resuspended cDNAs were sequenced on a CEQ 8000 capillary electrophoresis sequencer (9+). The resulting traces were analyzed using QuSHAPE (43). Reactivity was obtained from at least three independent replicates. Reactivity value was set arbitrarily to -10 for the nucleotides for which it could not be determined (intrinsic RT stops). Secondary structures were modeled using IPANEMAP (44) and drawn with VaRNA (45). Probing profiles were compared as described (40).

### Animal studies

Female Swiss mice (Janvier Labs), 6-8 weeks old and weighing ∼30 g were used for the septicemia model. Experiments were monitored in the ARCHE-BIOSIT animal lab in Rennes, and were performed in biological duplicates. We used groups of 5 mice for the mild septicemia model. Mice were infected intravenously by the tail vein with 200 μl of suspensions in 0.9% NaCl containing 3×10^8^ bacteria with either HG003, HG003 Δ*isrR*, or HG003 Δ*isrR locus2*::*isrR*^+^ strains. Mouse survival was monitored for 8 days, and the statistical significance of difference(s) between groups was evaluated using the Mantel-Cox test. A *p* value < 0.05 was considered as significant. All experimental protocols were approved by the Adaptive Therapeutics Animal Care and Use Committee (APAFiS #2123–2015100214568502v4).

## RESULTS

### IsrR is an sRNA required for growth in iron-depleted media

Iron is essential for *S. aureus* pathogenicity (46) and sRNAs are ubiquitous regulators for adaptation (47); we asked whether sRNAs are required in this bacterium to adapt to low iron availability, a condition encountered during infection. To address this question, we took advantage of a strategy that we recently developed, based on a competitive fitness evaluation of sRNA mutant libraries (25,48). A library of DNA-tagged deletion mutants was constructed in the HG003 strain (Supplementary Table S1); three independent mutants were constructed for each locus, to be subsequently used to constitute three independent libraries (Supplementary Table S4). Gene deletions of sRNAs corresponding to UTR regions or antisense from coding sequences more likely lead to phenotypes due to their associated coding genes; such mutations would interfere with the screening procedure designed to uncover the fine-tuning activity of sRNAs. We therefore restricted our collection to 48 mutants corresponding to nearly all known HG003 “*bona fide* sRNAs”, defined as those that are genetically independent with their own promoter and terminator (49). Deletion and substitution of these genes by tag sequences were designed to avoid interference with the expression of adjacent genes. The deleted region of each mutant was replaced by a specific DNA tag, which allowed us to count each mutant and therefore evaluate their proportion within a population of mutant strains. Using an indexed PCR primer pair, up to 40 samples can be tested in one DNA-seq run (25). Mutants that disappeared or accumulated in a given stress condition indicated a functional role of the corresponding sRNAs with respect to the imposed growth conditions.

The triplicate libraries were challenged to iron depletion by the addition of iron chelators, 2,2’-dipyridyl (DIP) and ethylenediamine-N,N’-bis(2-hydroxyphenyl)acetic acid (EDDHA) to growth media. The proportion of each mutant within the population was determined at different growth steps over 3 days (Figure 1A). Results were normalized to the same medium without iron chelator. Among 48 tested mutants, the strain with tag135 bearing a tagged deletion between the *arlR* and *pgpB* operons had a significant fitness disadvantage with each of the two tested iron chelators (Figure 1B); after about 28 generations, its distribution compared to the control condition decreased more than 10- and 1000-fold in DIP and EDDHA, respectively.

The deletion associated with tag135 inactivated an sRNA reported in two global studies under the names S596 (50) and Tsr25 (51). It is renamed here IsrR for <u>i</u>ron-<u>s</u>paring <u>r</u>esponse <u>r</u>egulator. Neither *rnaIII, rsaA, rsaC, rsaD, rsaE, rsaG, rsaOG, ssr42 nor ssrS* deletion mutants (49) present in the tested libraries were impacted by the presence of either DIP or EDDHA, indicating that their corresponding sRNAs do not significantly contribute to adaptation to low iron conditions in the tested conditions.

The HG003 Δ*isrR* strain had no apparent growth defect in rich medium compared to its parental strain (Figure 1C and Supplementary Figure S1). However, by spot tests, a Δ*isrR* mutant displayed reduced colony size in the presence of DIP and EDDHA, supporting the fitness experiment results and evidencing that the individual Δ*isrR* mutant, in the absence of the other 47 mutants, is required for optimal growth when the iron supply is limited (Figure 1C and Supplementary Figure S1). Of note, the slight growth retardation of Δ*isrR* observed on iron-depleted plates or in liquid cultures corresponds to a drastic fitness cost when in competition with other bacteria. To confirm that the phenotype was solely due to the absence of IsrR, a copy of *isrR* with its endogenous promoter was inserted in the chromosome of the Δ*isrR* strain at two different loci, leading to either Δ*isrR locus2*::*isrR*^+^ or Δ*isrR locus3*::*isrR*^+^ strains (Supplementary Figure S2A). Loci 2 and 3 were chosen since these regions were located between two terminators and seemed not to be transcribed. *isrR* expression from *loci 2* and *3* was confirmed by Northern blot (Supplementary Figure S2B). The two Δ*isrR* strains carrying a single copy of the *isrR* gene had plating efficiencies equivalent to that of the isogenic parental strain when grown with DIP or EDDHA, with colony sizes similar to those of the parental strain. These results confirm that the observed Δ*isrR* phenotype was strictly dependent on *isrR* (Figure 1C and Supplementary Figure S2C). Of note, HG003 carrying p*IsrR* (IsrR expressed from its endogenous promoter) leads to a 100-fold reduced ability to form colonies on iron-depleted media compared to the strain carrying a control vector (Figure 1D). Multicopy *isrR* expression is deleterious for *S. aureus*, possibly due to the disruption of important metabolic pathways; thus, we cannot rule out that the effect is due to an IsrR off-target activity. We conclude that the absence of IsrR is detrimental to *S. aureus* growth under iron-depleted conditions.

### IsrR expression is repressed by the ferric uptake regulator Fur

Extremities of *IsrR* were determined by 5’/3’RACE experiments after circularization of total RNAs (Supplementary Figure S3A). *isrR* transcription starts at position 1362894 and terminates at position 1363067 on the NCTC8325 genome map, generating a 174 nucleotide-long sRNA (Figure 2A) in agreement with tilling array data (50). *isrR* is preceded by a σ^A^ -dependent promoter and ends with a rho-independent terminator (50). Its high expression in a chemically-defined medium and the presence of a putative Fur-box within the promoter region suggested iron-dependent expression of *isrR* (50). Fur-dependence of *isrR* regulation was verified in a HG003 Δ*fur* mutant (Supplementary Table S1). In rich medium, IsrR was not detected by Northern blotting in the parental strain but strongly accumulated in the Δ*fur* mutant, demonstrating that Fur negatively controls *isrR* expression (Figure 2B). The *isrR* gene is predicted to be controlled by two Fur-boxes, one within the promoter region and the second immediately downstream of the transcriptional start site (Figure 2A). To test their relative contributions, a plasmid-based transcriptional reporter system placing a *gfp* gene under the control of the *isrR* promoter was constructed (P_isrR_::*gfp*) (Supplementary Table S2). The contribution of each predicted Fur box was tested by targeted mutagenesis altering either the first (P_isrR1_::*gfp*), the second (P_isrR2_::*gfp*), or both (P_isrR1&2_::*gfp*) Fur-binding motifs (Figure 2C). As expected, the transcriptional reporter generated no fluorescence in a *fur*^+^ strain (Figure 2D). Alteration of the upstream box led to strong fluorescence, which was even more intense when both boxes were mutated. In contrast, mutations only in the downstream Fur motif had no effect on the expression of the reporter fusion. We concluded that both sites contribute to efficient *isrR* Fur-dependent repression, the first one being epistatic over the second.

**Figure 2.**
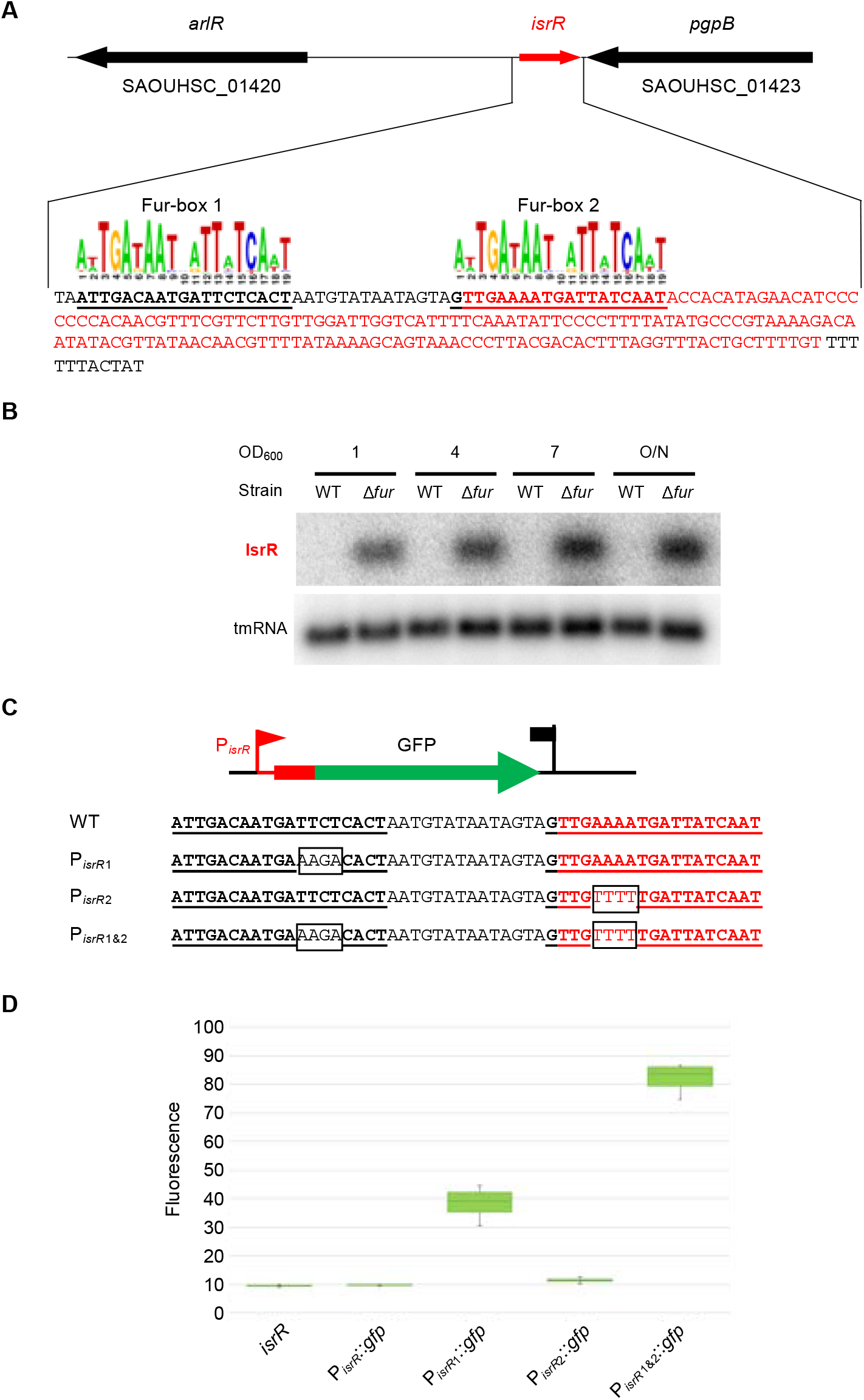
IsrR is regulated by Fur. (A) *isrR* genetic locus. The transcribed region is in red. For *isrR* 3’-5’RACE mapping results, see Supplementary Figure S3A. Predicted Fur boxes are bold and underlined. The staphylococcal Fur box consensus is taken from the RegPrecise database (30). (B) Northern blot experiment with HG003 and its *fur* derivative sampled at the indicated OD_600_ and probed for IsrR and tmRNA (control) (n=2). (C) *isrR* Fur box sequence (WT) and its mutant derivatives (P_isrR1_, P_*i*srR2_ and P_isrR1&2_). Mutations altering the Fur boxes are shown in black boxes. (D) Fluorescence of transcriptional fusions placing *gfp* under the control of the *isrR* promoter and its mutant derivatives (as indicated) measured by flow cytometry, (n=5). The first strain indicated *‘isrR’* is a control strain in which the *isrR* gene is not fused to *gfp*.

### IsrR and its regulation by Fur are conserved in the *Staphylococcus* genus

Most trans-acting bacterial sRNAs are poorly conserved across species (52,53), including among the staphylococci (49,54). However, the *isrR* gene was detected in all screened genomes from the *Staphylococcus* genus (Figure 3A and Supplementary Table S5). Sequence conservation includes the two Fur boxes, which were detected for all *isrR* gene sequences of the different *Staphylococcus* species.

**Figure 3.**
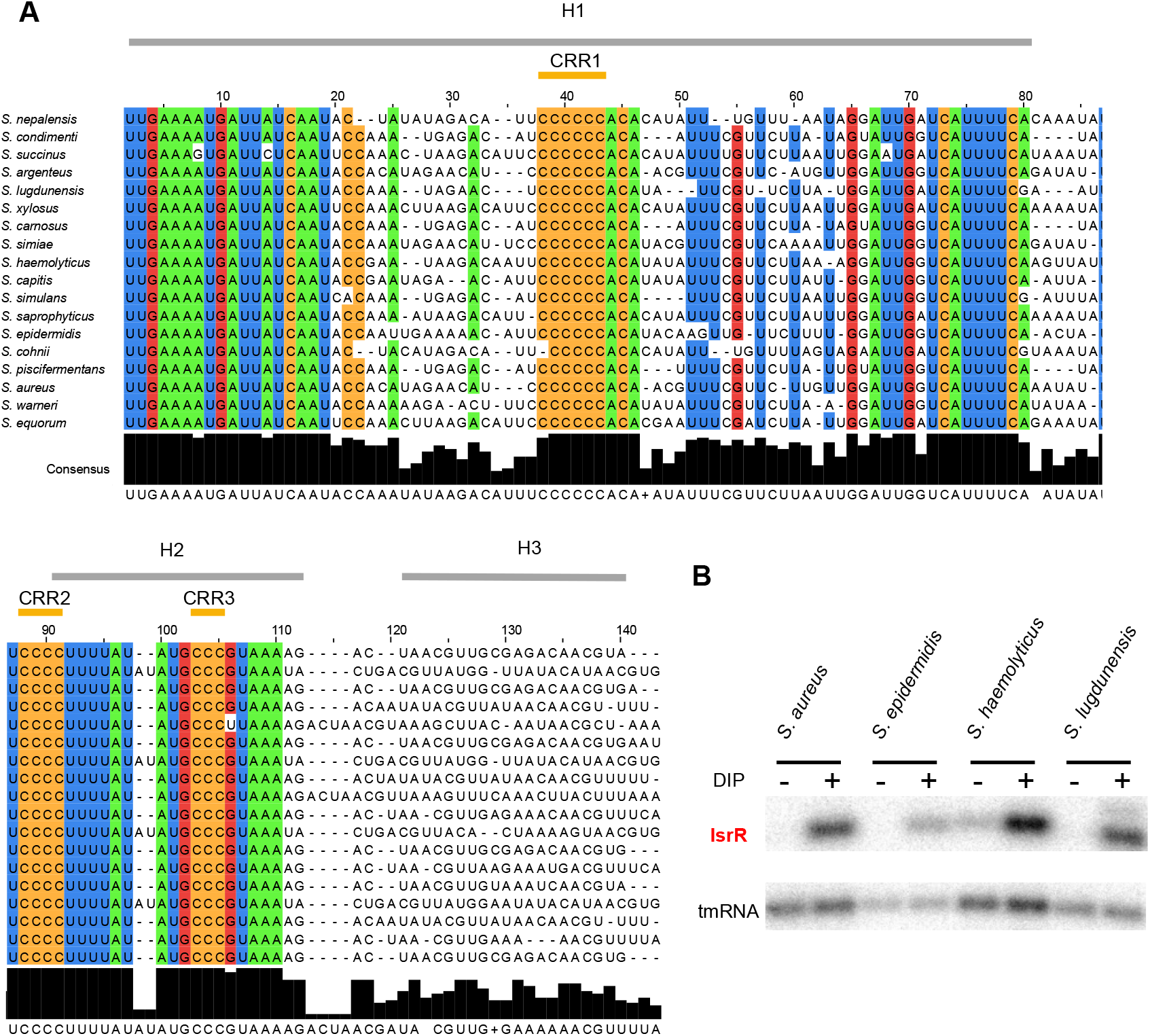
IsrR is conserved and Fur-regulated within the *Staphylococcus* genus. (A) Sequence conservation. The alignment was obtained using LocARNA (31) with *isrR* sequences from indicated strains generated by GLASSgo (28) as inputs. 80% conserved nucleotides within the tested sequences are colored (green, A; blue, U; red, G; ocher, C). Sequences shown are limited to the three first stem-loops (H1 to H3). There is poor nucleotide sequence conservation for H3 and the transcriptional terminator. Three C-rich regions are indicated (CRR1 to CRR3). (B) Northern blot probed for IsrR in *S. aureus* HG003, *S. epidermidis ATCC 12228, S. haemolyticus JCSC1435, and S. ludgunensis N920143*. Bacteria were grown in BHI or BHI supplemented with DIP 1.25 mM and were withdrawn at OD_600_ 1 (n=2).

IsrR orthologs from different species of the Staphylococcus genus were used to feed LocARNA, a software for multiple alignments of RNAs that generates secondary structure predictions and highlights conserved pairing regions (31). The proposed IsrR structure includes three stem-loops (H1 to 3) and a canonical Rho-independent terminator (T) (Supplementary Figure S3B). Three <u>c</u>onserved <u>C</u>-rich regions, named here CRR1 to 3, are predicted to be single-stranded. CRR1 and 3 are within the loops of stem-loop structures H1 and H2, while CRR2 is at the 5’-end abutting stem H2. CRR1 is four to seven C-long, according to the species considered. C-rich motifs are important for sRNA/target recognition in *S. aureus* (55-57). This feature efficiently discriminates targets in staphylococcal strains, since their genomes are ∼70% AT-rich, and G-rich stretches are infrequent except within Shine-Dalgarno sequences. Two first stems, H1 and H2, have conserved primary sequences with little covariation, indicating the importance of each nucleotide, possibly for an interaction with a yet unknown protein. Of note, single-stranded regions adjacent to CRR and within the loops are more variable than H1 and H2. To confirm experimentally its secondary structure, IsrR was subjected to SHAPE chemical probing, a technology revealing single-stranded nucleotides. RNAs were incubated with 1-Methyl-7-nitroisatoic anhydride (1M7), which specifically reacts with flexible nucleotides (58,59). Reactivity data was implemented into the IPANEMAP workflow to predict IsrR secondary structure according to thermodynamics and probabilistic calculations balanced with the experimental data (44). The experimental model obtained (Figure 4) is identical to the one obtained using the phylogenetic approach described above. Most nucleotides with high reactivity were identified in predicted single-stranded regions. However, several nucleotides with medium to high reactivity are in predicted double-stranded regions probably reflecting the poor stability of such A-U rich structures. Surprisingly, CCR1 is weakly reactive, and CRR2 and CRR3 are only mildly reactive, suggesting that although modeled in single-stranded regions, these Cs are involved in some interactions. As IsrR does not feature any stretches of G, Cs from the CRRs might be involved in yet to be identified non-canonical interactions. Of note, such interactions could actually expose the C’s Watson-Crick face to the solvent making them available for an intermolecular interaction. To detect divalent ion-dependent tertiary folding, IsrR was probed in the presence or absence of Mg^2+^ ions (Supplementary Figure S3C). Only a few significant local reactivity changes located after CCR1 stem-loop were observed suggesting that this region of IsrR does not adopt an extensive tertiary structure (60).

**Figure 4.**
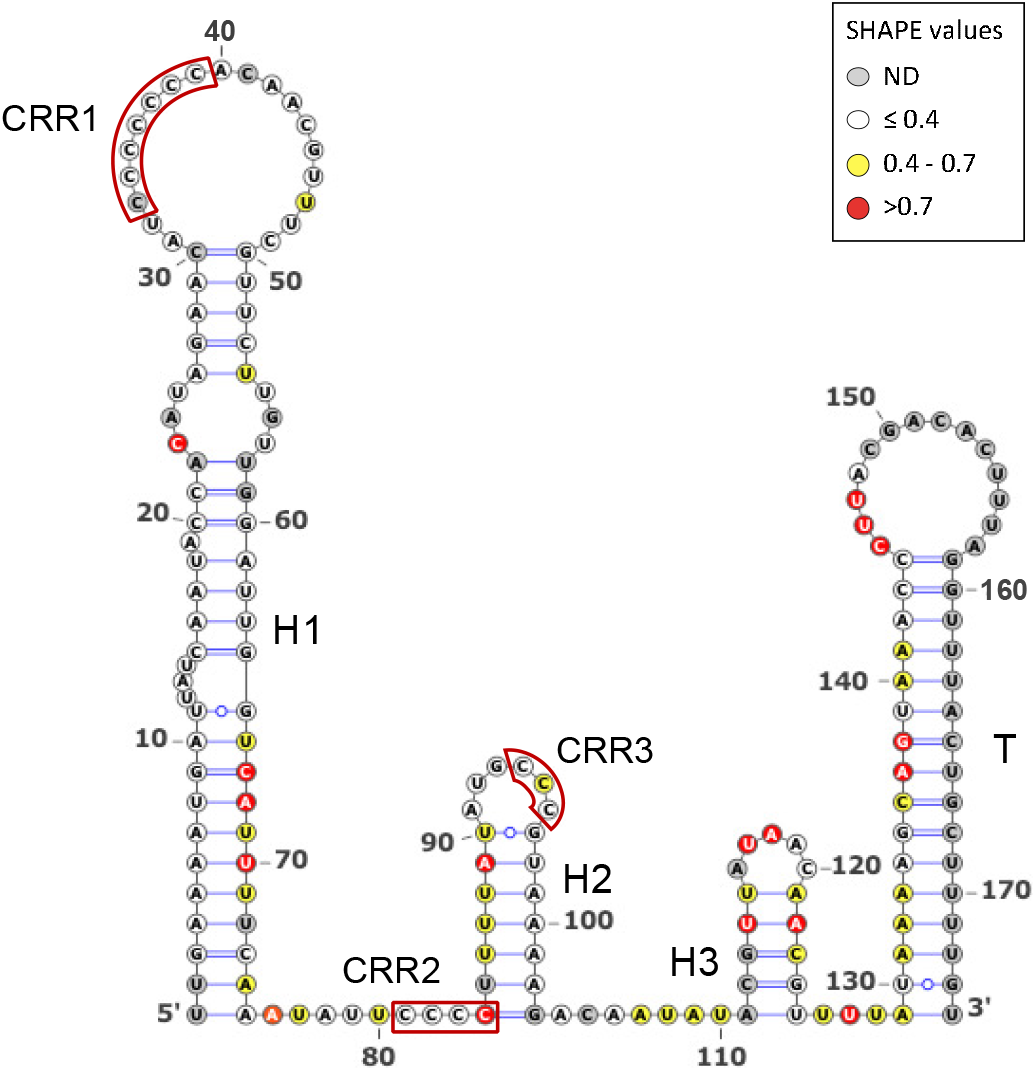
IsrR secondary structure model obtained using IPANEMAP. IsrR was probed with 1M7 at 37°C in 40 mM HEPES pH 7.5, 150 mM KCl with/without 5 mM MgCl_2_. White: low reactivity to 1M7, yellow: moderate reactivity, red: high reactivity. Nucleotides in grey denote undetermined reactivity. Specific regions of the IsrR structure are indicated: Three stem-loop structures (H1 to H3), a rho-independent transcription terminator (T), and three C-rich regions (CRR1 to CRR3).

The expression and iron dependency of *IsrR* were experimentally confirmed for *S. haemolyticus, S. epidermidis*, and *S. lugdunensis* (Figure 3B). Our results suggest that *IsrR* has an important function related to iron metabolism that is conserved throughout the Staphylococcus genus. The presence of CRRs suggests that *IsrR* is likely a conserved transacting small regulatory RNA.

### Bioinformatics predictions indicate that IsrR targets mRNAs encoding enzymes with Fe-S clusters

As *trans*-acting bacterial sRNAs base-pair to RNAs, it is possible to predict their putative targets by bioinformatics; however, some predictive programs generate false positive candidates (61). We used CopraRNA, a comparative prediction algorithm for small RNA targets, which takes into account RNA sequence accessibility within sRNA structures and the evolutionary conservation of the pairings (32). It is reportedly an efficient sRNA-target predictor when ortholog sRNA sequences are detected among different species, especially when they are phylogenetically distant (61). As IsrR is a conserved sRNA in the *Staphylococcus* genus, IsrR ortholog sequences were used as input for CopraRNA. Results indicated a functional convergence of the proposed targets (Supplementary Figure S4). Among the 23 top candidates with the best e-values on pairings, seven are mRNAs that encode iron-sulfur (Fe-S)-containing proteins (NasD, MiaB, GltB2, FdhA, CitB, NarG, and SAOUHSC_01062). Several of these *IsrR* putative targets were previously proposed by Mäder et al. using the same approach (50). Considering 32 different Fe-S containing proteins in *S. aureus* identified by manually curated *in silico* analyses (detailed in Supplementary Table S6), the probability of identifying seven mRNAs encoding Fe-S containing proteins by chance is 2×10^−9^ (see Methods). This remarkably low probability allows one to state with high confidence that Copra RNA identified relevant IsrR targets. Other putative targets, such as *moaD* and *nreC* mRNAs, are also associated with Fe-S containing complexes (62,63). It is therefore likely that most, if not all, predicted targets associated with Fe-S clusters are direct IsrR targets. Remarkably, nitrate reductase, nitrite reductase, glutamate synthase, formate dehydrogenase, aconitase, and hydratase methionine sulfoxide reductase, which are all putatively affected by IsrR (Supplementary Figure S4), also correspond to either putative or demonstrated enzymes affected by RyhB (5,64) suggesting a striking functional convergence between these two sRNAs.

### IsrR targets mRNA encoding nitrate-related enzymes

In the absence of oxygen, *S. aureus* uses nitrate, if available, as an electron acceptor of the respiratory chain (65). Nitrate is converted to nitrite by nitrate reductase encoded by the *nar* operon (Supplementary Figure S5). In a second step, nitrite is converted to ammonium by nitrite reductase encoded by the *nas* operon. Ammonium is then used by glutamate synthase encoded by the *gltB2* gene. Transcription of *nar* and *nas* operons depends on the NreABC regulatory system, whose activity requires the presence of nitrate and the absence of oxygen (63). Strikingly, *narG* (nitrate reductase subunit α), *nasD* (nitrite reductase large subunit), and *gltB2* mRNAs are IsrR putative targets, suggesting that IsrR could affect all steps of nitrate to glutamate conversion.

In *E. coli*, the *fdhF* gene encoding a molybdenum-dependent formate dehydrogenase is regulated by nitrate (66). A strain with a mutated *fdhF* allele lacks nitrate reductase activity (67), suggesting that FdhF is associated with the nitrate dissimilatory pathway. Interestingly, the fdhF ortholog in *S. aureus, fdhA*, is also predicted to be an *IsrR* target.

To assess IsrR targets associated with the nitrate reduction pathway, we selected *fdhA, narG, nasD*, and *gltB2* mRNAs. Detection of long transcripts by Northern blot is often not possible in *S. aureus*. For this reason, the effect of *IsrR* on the amount of its targets was tested by Northern blots solely for *fdhA* and *gltB2* mRNAs, which are expressed from short operons (bi- and mono-cistronic, respectively). In many cases, the interaction of an sRNA with its mRNA target results in the destabilization of transcripts. Consequently, we questioned whether IsrR would affect *fdhA* and *gltB2* mRNA stability. RNA stability was evaluated upon transcription inhibition by rifampicin, where the variations in RNA quantities are assumed to reflect their degradation. This classical approach should nevertheless be interpreted cautiously since i) stability of sRNA and mRNA targets depends on their base pairing and that ii) rifampicin prevents the synthesis of both partners, one of them being possibly limiting (68). In HG003 and its Δ*isrR* derivative, *fdhA* and *gltB2* mRNAs were unstable in rich medium. Addition of DIP to the growth media resulted in a minor increase of both mRNAs in HG003 and Δ*isrR* strain (Figure 5). However, *IsrR* does not have a significant effect on *fdhA* and *gltB2* mRNA stability, with the caveat associated with the possible limitation of using rifampicin to assess regulatory RNA activities.

**Figure 5.**
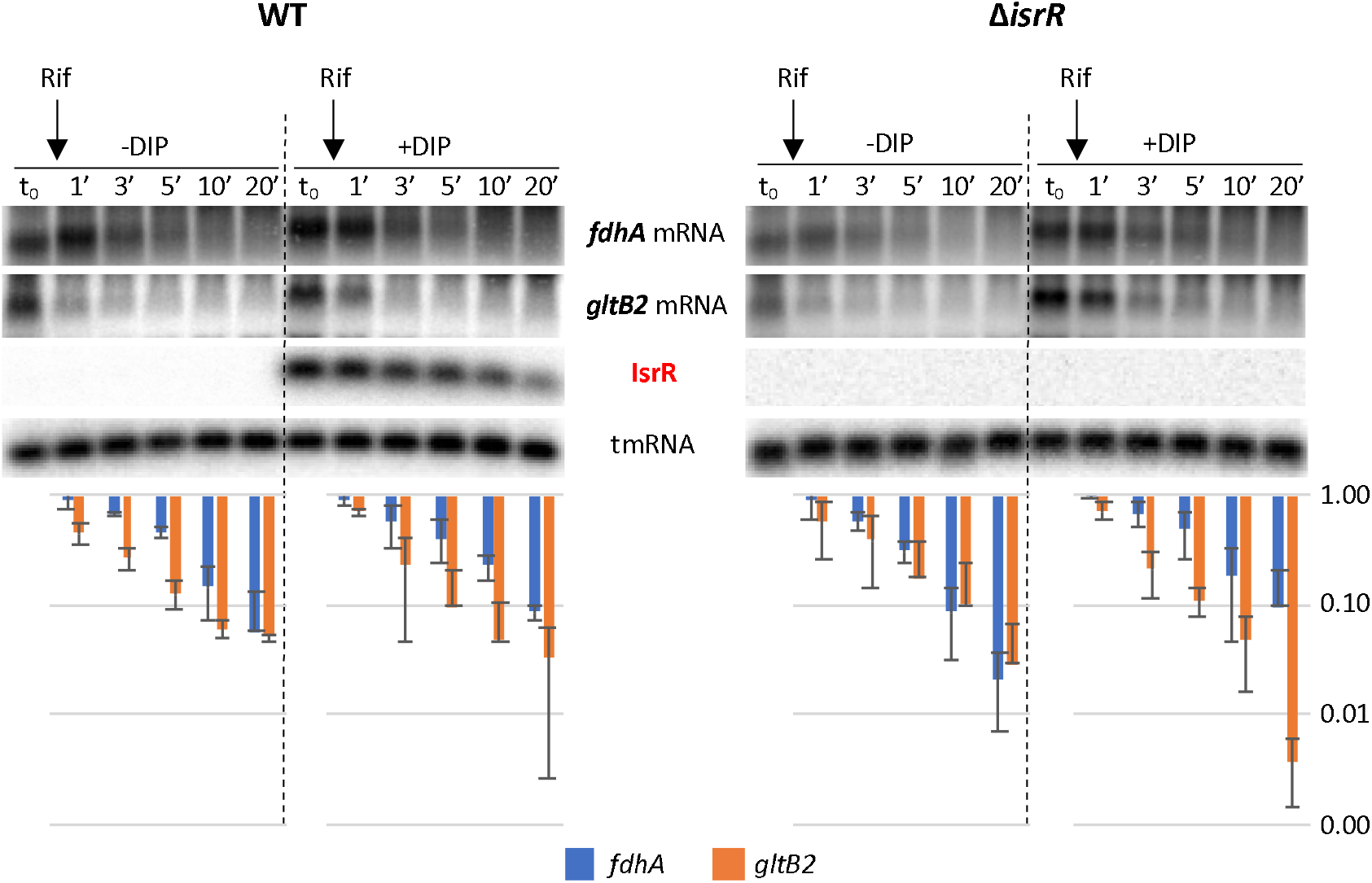
Stability of *fdhA* and *gltB2* mRNAs is not significantly affected by IsrR. HG003 (left panel) and its isogenic Δ*isrR* derivative (right panel) were grown in rich medium with or without the addition of DIP (as indicated). At t_0_, rifampicin (Rif) was added to the growth medium. Cultures were sampled at t_0_, 1, 3, 5, 10, and 20 min after addition of rifampicin, total RNA was extracted, and the amounts of *fdhA* mRNA, *gltB2* mRNA, tmRNA (loading control), and IsrR were assessed by Northern blot. Histograms show the quantification of *fdhA* and *gltB2* mRNAs normalized to t_0_ from two rifampicin assays as shown in the upper panel. Vertical axis, arbitrary units. Error bars indicate the standard deviation from two independent experiments (n=2).

We used the SHAPE technology to probe and model the secondary structure of *fdhA, gltB2, narG*, and *nasD* 5’UTR as well as the heteroduplex they form with IsrR. To this end, RNA either corresponding to the four 5’UTR alone or in complex with *IsrR* were submitted to a SHAPE probing experiment with 1M7 (*IsrR* structure probing and modelling is reported above). These experiments were carried out in triplicate and the results were used as constraints to model RNA secondary structure according to the IPANEMAP workflow (43). IPANEMAP yielded secondary structure models very consistent with the SHAPE probing data for *fdhA, gltB2* (Figure 6) and *nasD* (Supplementary Figure S7) mRNAs, whereas the *narG* mRNA reactivity map did not allow modelling of a stable structure, suggesting it does not adopt one. Interestingly, *fdhA* Shine-Dalgarno sequence is embedded in a stable structure whereas the initiation codon is highly reactive while the opposite is observed for *gltB2* mRNA. The limited availability of either of these sequence elements suggests that the expression of these two genes is regulated by their own RNA structure.

**Figure 6.**
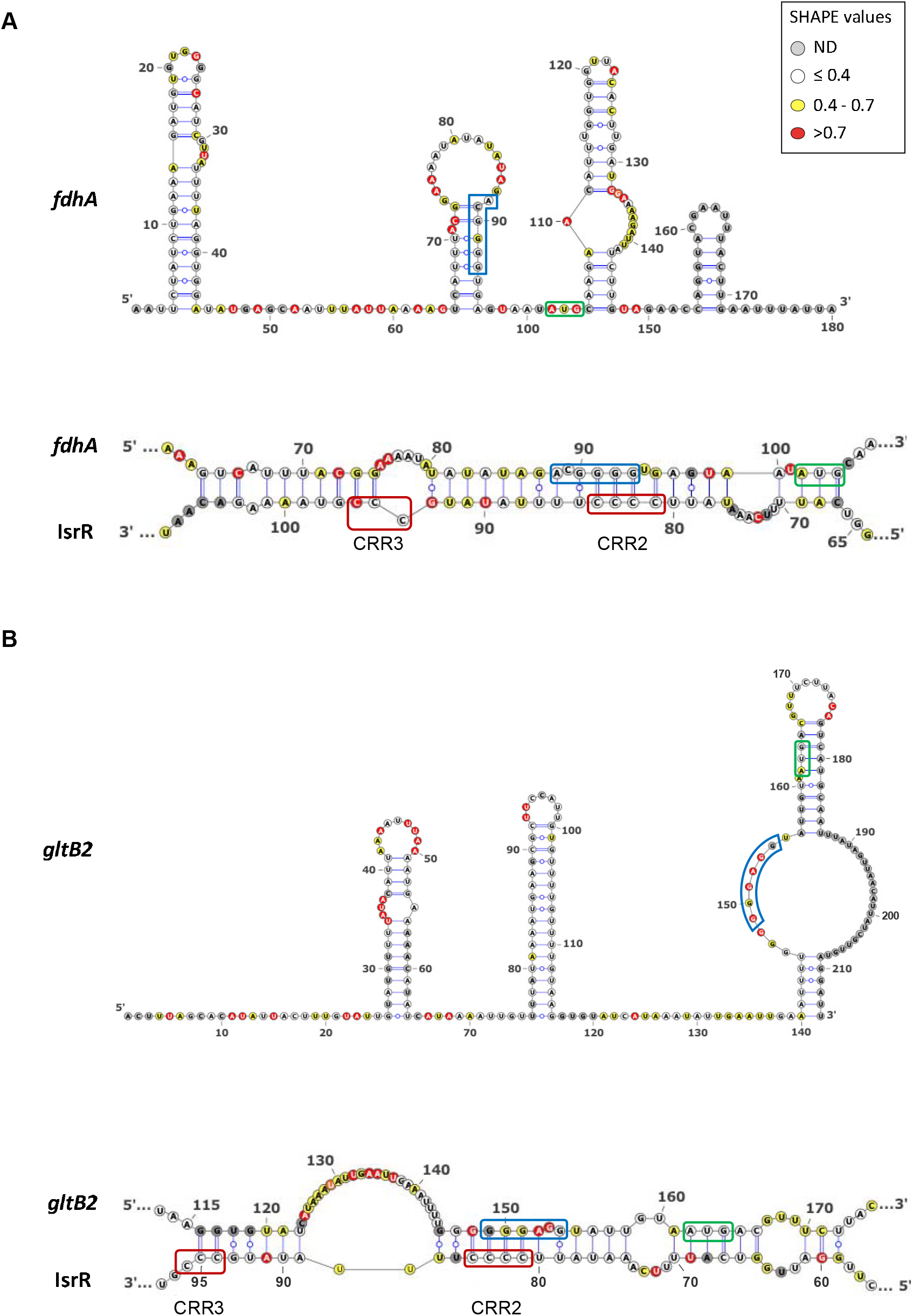
Secondary structure models for *fdhA* and *gltB2* 5’UTR alone or in interaction with IsrR. Top panels: Secondary structure model obtained with IPANEMAP for *fdhA* (A) and *gltB2* (B) 5’UTRs using 1M7 reactivity as constraints. Nucleotides are coloured according to their reactivity in the absence of IsrR with the indicated colour code. ND, not determined. Bottom panels: model for interaction between IsrR and *fdhA* (A) and *gltB2* (B) mRNAs based on changes in reactivity in the presence of IsrR. mRNA Shine-Dalgarno sequence is shown in a blue rectangle and the start codon in a green rectangle; IsrR C-rich regions are shown in red rectangles.

Statistically significant reactivity variations for specific nucleotides were observed for IsrR and each of its mRNA targets when incubated alone or as an IsrR/5’ UTR target pair (Supplementary Figures S6 and S7). Such data strongly support an interaction of IsrR with the four putative mRNA targets tested. Nucleotides for which reactivity decreases are likely to be involved in base pairing between IsrR and a target mRNA, whereas an increase of reactivity probably reflects the destabilization of a structure upon formation of the sRNA/mRNA heterodimer (*e*.*g*., the initiation codon of *fdhA* and the region including the Shine-Dalgarno of *gltB2*, which were highly reactive in the absence of IsrR, present a significantly decreased reactivity in the presence of *IsrR*). Interestingly, no significant differences are observed in IsrR 50 first nucleotides, suggesting that CCR1 is not involved in the interaction with the mRNA targets. These data were used as constraints for IPANEMAP to model the interaction between *IsrR* and *fdhA, gltB2*, and *nasD* mRNAs. The interactions proposed include IsrR CRR2 and CCR3 as well as the Shine-Dalgarno and the AUG initiator codon on the mRNA target side, suggesting that IsrR impairs translation of its targets.

The binding of IsrR to one putative nitrate-related target, *fdhA* mRNA, was tested by electrophoretic mobility shift assay (EMSA). An IsrR band shift was observed in the presence of *fdhA* mRNA. The specificity of binding was challenged with a specific competitor (unlabelled *IsrR*) and a nonspecific competitor (polyU) (Supplementary Figure S8A). As expected, unlabelled IsrR, but not polyU, displaced the interaction, confirming the specificity of the IsrR/*fdhA* RNA duplex. To identify nucleotides contributing to the complex formation, EMSA was performed with IsrR and *fdhA* mRNA harbouring point mutations in the predicted interaction zone (Supplementary Figure S8B). Because the interaction between IsrR and *fdhA* mRNA involves likely 31 base-pairs, the replacement of several nucleotides was necessary. The presence of seven point mutations altering either IsrR or *fdhA* mRNA prevented band shifts when associated with their corresponding wild-type partner. However, when the mutated IsrR was incubated with *fdhA* mRNA harbouring compensatory mutations restoring the pairing, a band shift was observed (Supplementary Figure S8C). The decreased amount of RNA duplex with both mutated RNAs can be explained by a less stable complex as predicted by IntaRNA software (69) (ΔG=-16.98 kcal/mol for WT IsrR/*fdhA* duplex *vs*. ΔG=-12.8 kcal/mol for mutated IsrR/*fdhA* duplex). Our data support a direct interaction of IsrR and mRNAs associated with nitrate metabolism.

### Translational control by IsrR

Our *in vitro* experiments indicate a direct interaction of IsrR with its targets. However, at least in the case of *fdhA* and *gltB2* mRNAs, this pairing does not trigger mRNA degradation, therefore, suggesting that IsrR can act on translation without affecting the stability of targeted mRNAs. To support this hypothesis, translation reporter systems were constructed. 5’UTR sequences of predicted mRNA targets of IsrR were fused to a reporter gene. For these experiments, sequences corresponding to the 5’UTRs and the first codons of mRNA targets were cloned under the control of the P1 *sarA* promoter in frame with the GFP coding sequence (p5’FdhA-GFP, p5’NarG-GFP, p5’NasD-GFP, p5’GltB2-GFP; Supplementary Figure S9A). The cloned regions comprise the sequences predicted to pair with IsrR. Note that transcription of the 5’UTRs from P1 *sarA* promoter is likely constitutive and IsrR-independent. The *isrR* gene was placed under the control of the P_tet_ promoter on a multicopy plasmid (pRMC2ΔR-IsrR), and mutants lacking either the first, second or third C-rich motif were constructed, leading to pRMC2ΔR-IsrRΔCRR1, pRMC2ΔR-IsrRΔCRR2, and pRMC2ΔR-IsrRΔCRR3, respectively (Supplementary Table S1). The Δ*isrR* strains containing the different reporters were transformed with pRMC2ΔR (control plasmid), pRMC2ΔR-IsrR, and its derivatives. We confirmed that *isrR* and its ΔCRR derivatives were constitutively expressed (Supplementary Figure S9B). For each strain containing one of the four reporter genes, expression of IsrR led to reduced fluorescence (Figure 7 and Supplementary Figure S10).

**Figure 7.**
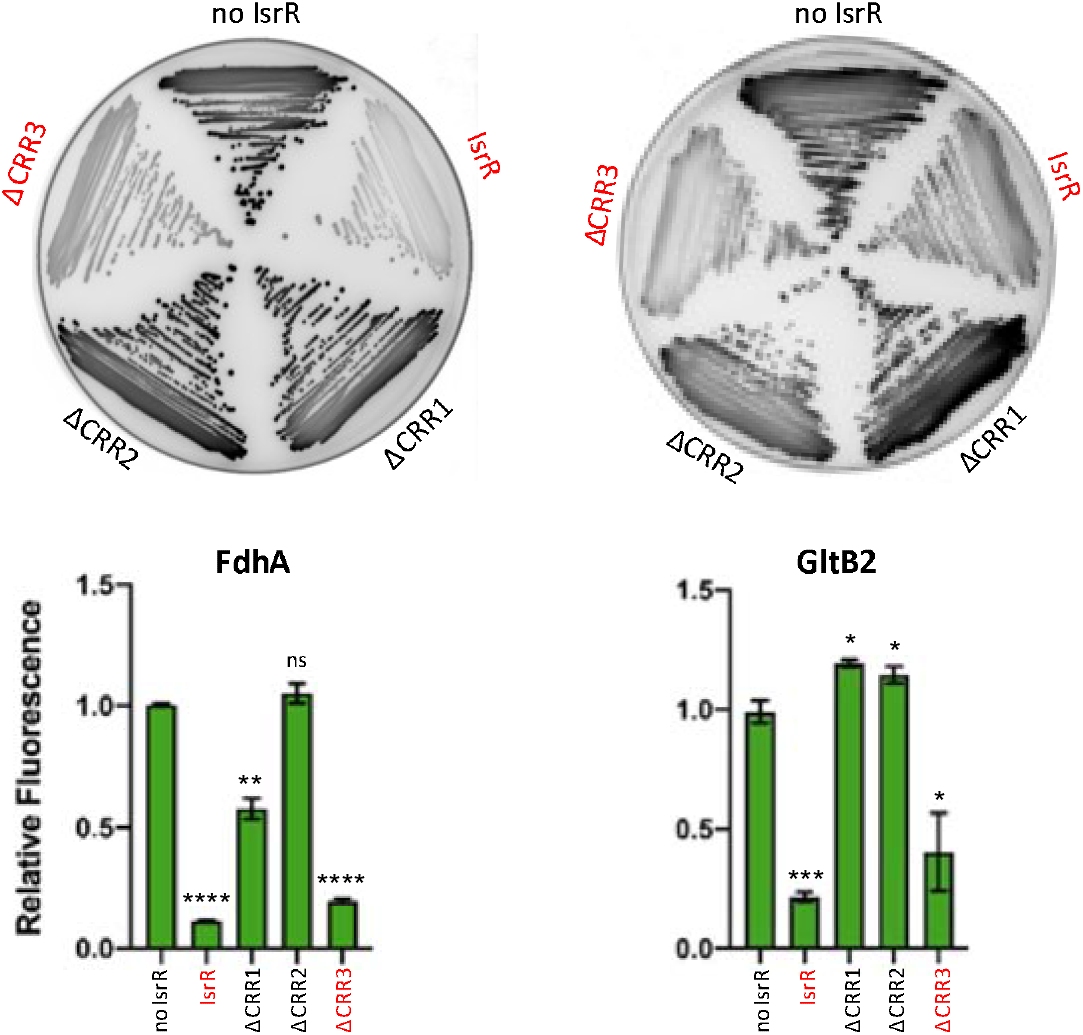
Translational down-regulation by IsrR and the CRR contribution. Leader fusions between the first codons of *fdhA* or *gltB2* and GFP were constructed (Supplementary Figure S9 and Table S2). The cloned fragments include the interaction regions with IsrR as described (Figure 6). HG003 Δ*isrR* derivatives with either a control plasmid (no IsrR; pRMC2ΔR), or plasmids expressing IsrR (pRMC2ΔR-*isrR*), IsrRΔCRR1 (pRMC2ΔR-*isrR*ΔCRR1), IsrRΔCRR2 (pRMC2ΔR-*isrR*ΔCRR2), IsrRΔCRR3 (pRMC2ΔR-*isrR*ΔCRR3) were transformed with each engineered reporter gene fusion. Translational activity from the reporters in the presence of the different *isrR* derivatives were evaluated by fluorescence scanning of streaked clones on plates (n=3). Fully active IsrR derivatives are shown in red. Translational activity of the reporter genes with the different *isrR* derivatives was also determined in liquid culture. Fluorescence of the strains was measured in 6 h cultures using a microtiter plate reader. Results are normalized to 1 for each fusion with the control plasmid. Error bars indicate the standard deviation from three independent experiments (n=3). Statistical analyses were performed using a t-test with Welch’s correction: **** represents p-value <0.0001, *** represents p-value of 0.0003, ** represents p-value of 0.0029, * represents p-value between 0.0111-0.0184.

The integrity of CRR1 was required for the IsrR activity against *fdhA* and *gltB2* reporter fusions. However, CRR1, despite being the largest CRR, was dispensable for IsrR activity against *narG* and *nasD* reporter fusions (Supplementary Figure S10). Interestingly, CRR2 was necessary for IsrR activity against all four reporter fusions. The observations are supported by SHAPE data. The integrity of at least two IsrR CRRs was required for activity against the four targets. These observations revealed that all CRRs are needed to mount the complete IsrR response, but differentially affect activity according to the given mRNA target.

### IsrR down-regulates nitrate metabolism

To determine if IsrR could indeed interfere with nitrate respiration, the amount of nitrite in anaerobic cultures upon addition of nitrate was measured in strains with either no IsrR or overexpressing IsrR. Two systems were used: i) Δ*isrR* carrying a plasmid control (pCont) compared to the same strain with a plasmid expressing *IsrR* (p-*IsrR*) and ii) Δ*fur* compared to Δ*fur* Δ*isrR* strains (Figure 8A), Note that *isrR* is constitutively expressed in the Δ*fur* background. In both systems, IsrR accumulation prevented nitrite production, as expected if nitrate reductase (*narG*) is indeed downregulated by IsrR.

**Figure 8.**
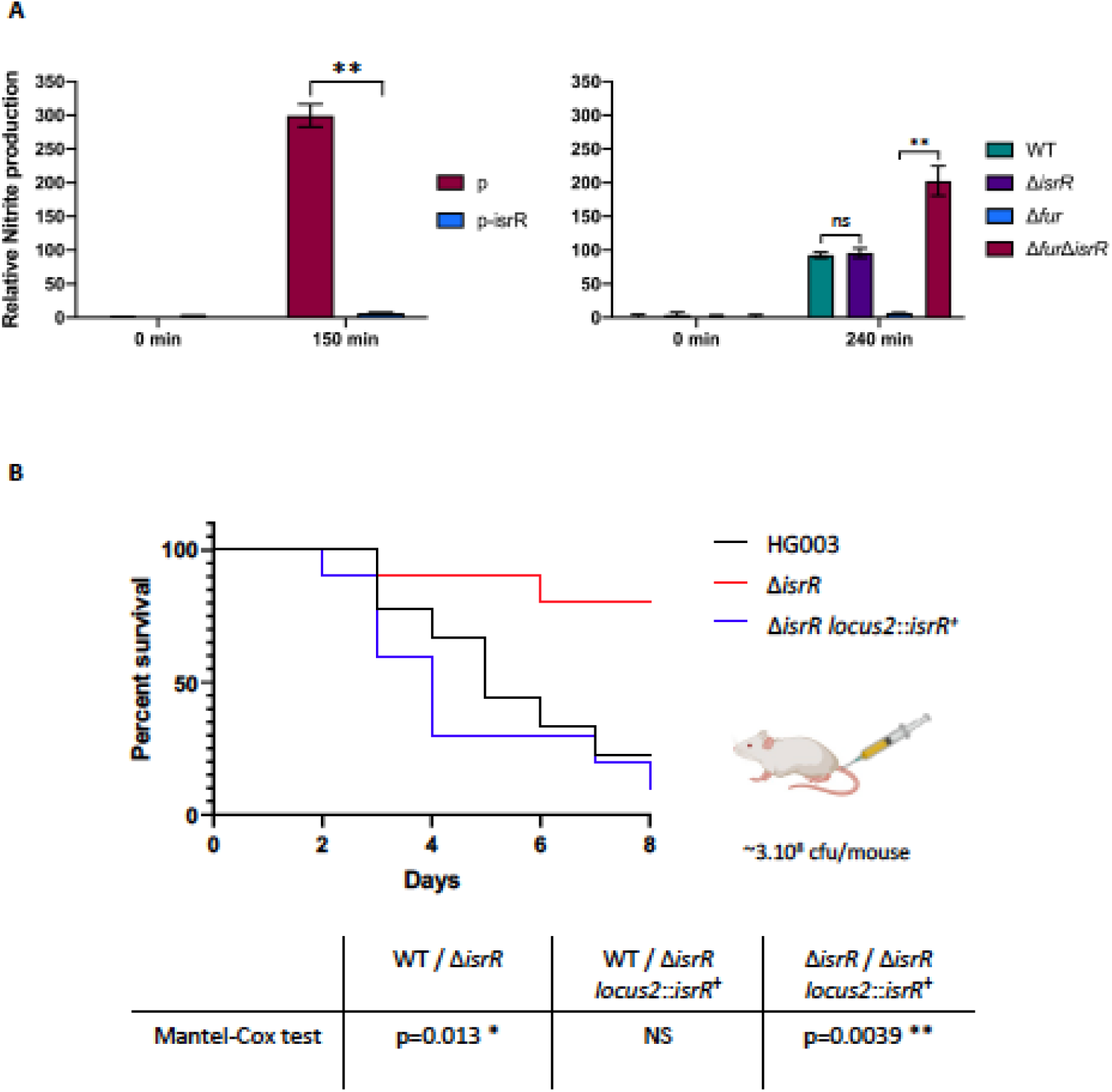
IsrR controls nitrite production and is required for virulence. (A) IsrR prevents nitrate conversion to nitrite. Strains were grown in anaerobic conditions for two hours and nitrate (20 mM) was added to cultures. Growth media: left panel, BHI; right panel, TSB. TSB medium was used here since we observed that Δ*fur* mutants grow poorly in BHI under anaerobic conditions. Samples for nitrite measurement were withdrawn at times 0 and 150 (left panel) or 240 min (right panel) upon nitrate addition. p, pRMC2ΔR; p-IsrR, pRMC2ΔR-IsrR. Histograms represent the relative nitrite concentration (Griess assay, OD_540_) normalized to the bacterial mass (OD_600_). Results are normalized to 1 for Δ*isrR* p (left panel) and WT (right panel) samples prior to nitrate addition. Error bars indicate the standard deviation from three independent experiments (n=3). Statistical analyses were performed using a t-test with Welch’s correction: ** represents p-value between 0.001-0.004; ns, non-significant. (B) Kaplan-Meier survival probability plots in a septicemia model of mice infected with either HG003 (WT, black), HG003 Δ*isrR* (red), and HG003 Δ*isrR locus2*::*isrR* (Δ*isrR* complemented, blue). Survival was monitored for 8 days post-infection. Results shown are from 10 mice per group; the experiment was performed twice, and data combined. The Mantel-Cox test was used to determine p values. NS, non-significant.

### IsrR activity is Hfq independent

In Enterobacteriaceae, Hfq is an RNA chaperone required for sRNA-mediated regulations (70). However, it does not seem to be the case in *S. aureus* (71-73), nor for *B. subtilis* (74-76) where its function remains enigmatic. Since RyhB activity is Hfq-dependent in *E. coli* (8), we asked whether IsrR would require a functional Hfq in *S. aureus*. The *fdhA, nasD, narG*, and *gltB2* translation reporter genes described above were introduced into Δhfq strains containing either a control plasmid or a plasmid constitutively expressing *IsrR*. As with the parental strain (Figure 7 and Supplementary Figure S10), the reporter fusions were downregulated by *IsrR* despite the absence of Hfq (Supplementary Figure S11A). It was conceivable that *isrR* cloned on a multi-copy plasmid produced high levels of sRNA that would bypass the need for Hfq. We therefore introduced the *fdhA* translation reporter in HG003, HG003 Δ*hfq*, and HG003 Δ*isrR*. When present, *isrR* expression was induced from its endogenous locus by the addition of DIP to the medium. On DIP-containing plates, the reporter fusion showed reduced activity in the parental and Δhfq strain compared to the strain lacking IsrR (Supplementary Figure S11B). Additionally, in vivo nitrite production in anaerobiosis was compared qualitatively between a HG003 WT strain and its Δ*hfq* derivative containing either a control plasmid or a plasmid constitutively expressing IsrR. 150 minutes after the addition of nitrate (NaNO_3_), the Δhfq strain expressing IsrR efficiently inhibited nitrite production as observed with the parental strain (Supplementary Figure S11C). Of note, IsrR induction through the addition of DIP was not used in this case since under anaerobic conditions, iron starvation results in severe growth arrest of the strains. We conclude that IsrR activity does not require Hfq for the tested phenotypes.

### IsrR RNA is required for *S. aureus* virulence

Host iron scavenging plays a crucial role in *S. aureus* infection (4,77). As the Δ*isrR* mutant has altered fitness in iron-restricted environments, we postulated that it could also impact virulence. We compared the virulence of Δ*isrR*, Δ*isrR locus2*::*isrR*^+^ and parental (HG003) strains injected intravenously in a mouse septicemia model (78). Of note, the absence of *isrR* does not alter *in vitro* growth in rich media (Figure 1 and Supplementary Figure S1) and *IsrR* is expressed in the Δ*isrR locus2*::*isrR* strain (Supplementary Figure S2B). Most mice inoculated with strains expressing a functional *isrR* (HG003 and Δ*isrR locus2*::*isrR*) were dead within 8 days (Figure 8B). In striking contrast, the lack of *IsrR* expression significantly reduced HG003 virulence, demonstrating the important role of this regulatory RNA during *S. aureus* infection in this model.

## DISCUSSION

The link between *S. aureus* pathogenicity and its iron status is established (79). The host exerts nutritional immunity that depletes iron in response to invading bacteria as a means to restrict infection. Here, we identified IsrR as the only sRNA, from 45 tested, required for optimum *S. aureus* growth in iron-restricted conditions and indeed, its absence attenuates *S. aureus* virulence.

IsrR was previously reported in a large-scale transcriptome study performed in numerous growth conditions (50), and was also observed to be the most highly upregulated sRNA when *S. aureus* growth in human serum was compared to that in a rich laboratory medium (51). Environments that failed to provide free iron, such as serum, promoted the induction of iron-repressed genes in infecting bacteria (80,81). Our findings on *isrR* regulation identify iron starvation as the signal for IsrR induction. RsaOG, RsaG, Teg16, SsrS and RsaD sRNAs were reported as upregulated in serum (51), but they do not contribute to *S. aureus* optimized fitness in the iron-starved conditions we tested. Supporting our observation, their corresponding genes are not preceded by Fur boxes (49,50) and their susceptibility to serum is likely not related to iron starvation. RsaC is an *S. aureus* sRNA produced in response to manganese (Mn) starvation that inhibits the synthesis of the superoxide dismutase SodA, a Mn-containing enzyme (82). RsaC is possibly the Mn counterpart of IsrR for iron and may contribute to adaptation to metal-dependent nutritional immunity. However, while putative RsaC targets are also associated with iron homeostasis, the *rsaC* mutant was not significantly affected by the chelating conditions tested here. *IsrR* is the only sRNA tested affected by both DIP and EDDHA and it is Fur regulated.

Inactivation of the Fur repressor leads to down-regulation of numerous genes (83,84). This paradoxical regulation could be achieved by a Fur-dependent expression of negative regulators, which act as ‘inverters’ of the Fur response, with IsrR being an *S. aureus* Fur inverter. Of note, several genes reported down-regulated upon iron starvation in *S. aureus* (85) are *IsrR* targets predicted by Copra RNA, including *narG* and *nasD*.

In the absence of oxygen, Staphylococci use nitrate and nitrite as electron acceptors of the anaerobic respiratory chain (86). These products are provided to humans by nutrients and are also generated by the oxidation of endogenous NO synthase products (87,88). Nitrate and nitrite reductases are expressed when needed, *i*.*e*., in anaerobic conditions when nitrate is present. Their expression is controlled by the staphylococcal transcriptional regulator NreABC (89-91). IsrR provides an additional checkpoint to nitrate respiration by linking transcription of *narG*HJ and *nirR nasDEF* operons to the presence of iron, an essential element of these encoded respiratory chain components (Figure 9).

**Figure 9.**
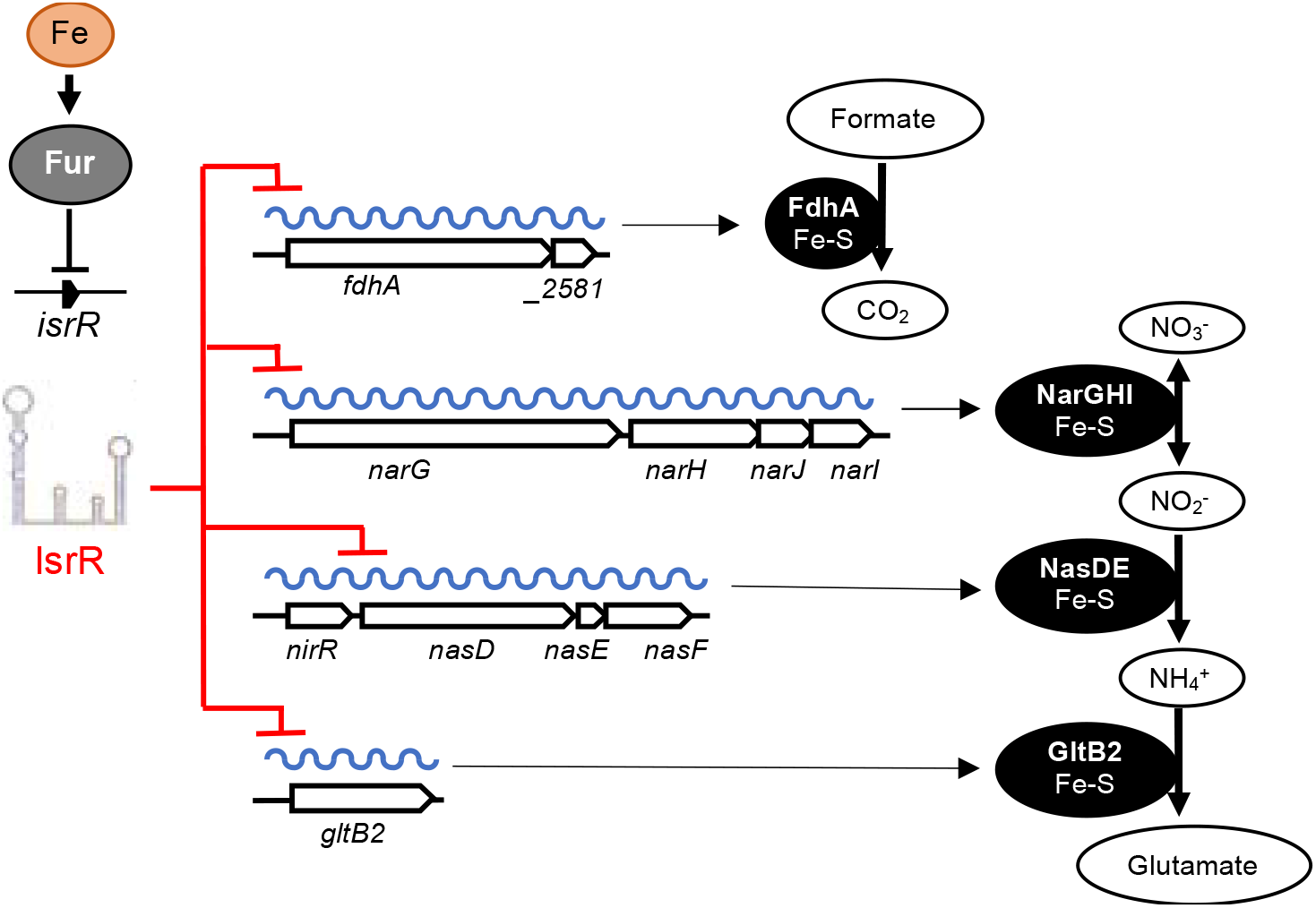
IsrR control of dissimilatory nitrate reduction. *isrR* is Fur-regulated. Its expression is induced in iron-free growth conditions. *IsrR* base pairs to SDs of mRNAs encoding components of formate dehydrogenase, nitrate reductase, nitrite reductase and glutamate synthase, thus preventing translation of the encoded iron-sulfur containing enzymes and nitrate dissimilatory reduction.

When iron is limiting, IsrR targets various mRNAs expressing iron-containing proteins that are not essential for growth. The dissimilatory nitrate reduction pathway is inactivated, while *S. aureus* can still grow by fermentation in anaerobic conditions. Biocomputing analysis (Supplementary Figure S4) suggests that IsrR targets *nreC* mRNA; If so, the expression of the nitrogen respiration pathway would be simultaneously downregulated by targeting mRNAs encoding dissimilatory nitrate reduction enzymes and their transcriptional activator. sRNAs that directly affect nitrogen metabolism have been characterized in Alphaproteobacteria, Gammaproteobacteria and Cyanobacteria (92-94); in several cases, the sRNAs directly downregulate regulators of nitrogen respiratory pathways. To our knowledge, IsrR would be the first sRNA example from the Firmicutes phylum.

A characteristic feature of sRNAs in Firmicutes and in particular in *S. aureus* is the presence of one or several exposed CRRs that can act on the G-rich region like those of the Shine Dalgarno sequences. The interaction models proposed from the probing experiments indicate that CRR2 and to a lesser extent CCR3 are involved in the base pairing of *IsrR* with *fdhA, gltB2*, and *nasD* mRNAs. Translational activity assays with a reporter fusion system further confirmed the CRR2 requirement to regulate expression of the mRNA targets. The results of the SHAPE footprinting do not suggest the involvement of CCR1 in the interaction with the mRNAs, although it appears important but not crucial for *fdhA* regulation. It should be noted that this assay only detects state-changing nucleotides and cannot detect those not reactive when the RNA is incubated alone, for example, CCR1 nucleotides. Therefore, it could be possible that the performed footprinting assay does not reveal an existing interaction involving CCR1. However, we did not observe any significant changes in IsrR 50 first nucleotides, suggesting that the long hairpin harboring CCR1 is not destabilized upon the interaction. We therefore suggest that the CCR1 role is not to base pair with the target. The complex models include both the SD sequence and the initiation codon further suggesting that IsrR acts primarily on translation rather than on mRNA stability.

IsrR-spared iron can be reallocated to vital processes. Consequently, IsrR plays a central role in *S. aureus* adaptation, including resistance to host nutritional immunity. Preventing IsrR activity by dedicated RNA antisense molecules or other means would antagonize staphylococcal pathogenicity.

Most staphylococcal sRNAs are poorly conserved across different species (49). One explanation is that the trans-acting sRNAs mostly act by imperfect pairing to untranslated regions (*i*.*e*., sRNAs and targeted UTRs); unlike ORFs, these sequences are prone to numerous inconsequential mutations and thus subject to rapid evolutionary changes. However, *isrR* and its Fur regulation are conserved throughout the *Staphylococcus* genus. This less-common interspecies conservation (another example in *S. aureus* is RsaE (95)) reveals a selective pressure to maintain IsrR sequence, structure, and regulation intact. IsrR pairs with several mRNA targets related to important functions, such as iron homeostasis, which may explain this conservation; the occurrence of random *isrR* mutations would affect iron-related metabolism, and therefore be counter-selected.

sRNAs that down-regulate mRNAs encoding iron-sulfur clusters are found in Gram-positive and Gram-negative bacteria. In addition to IsrR, these include RyhB in enteric bacteria (5), the paralogs PrrF1 and PrrF2 in *P. aeruginosa* (11), MrsI in *M. tuberculosis* (15) and FsrA sRNA in *B. subtilis* (96). Remarkably, none of these sRNAs share the same RNA chaperone dependency (e.g., Hfq for RyhB and PrrF1/PrrF2; FbpA, FbpB and FbpC for FsrA) with IsrR, and their genes share neither synteny nor sequence homology. FsrA, but not the other sRNAs, binds to the Shine-Dalgarno sequence via a C-rich motif (96), but requires three chaperone functions. These sRNAs are therefore not IsrR orthologs. Nevertheless, they do share several target mRNAs encoding the same iron-sulfur-containing enzymes (Supplementary Table S7). Indeed, the iron-sparing response of these sRNAs includes targets involved in nitrate metabolism and the TCA cycle. The TCA cycle mRNA encoding aconitase (acnA/citB) is targeted by all the sRNAs discussed above.

The common regulation of these sRNAs by iron and their shared targets suggest convergent evolution, which can be reasonably explained as follows: The accumulation of non-essential iron-containing enzymes is deleterious in iron-scarce environments; however, an sRNA induced during iron starvation that mutates to pair with these transcripts provides an immediate selective advantage, more energetically efficient than the production of specialized regulatory proteins. Since Fur is a widely conserved iron-dependent repressor in bacteria, any Fur-regulated sRNAs, possibly originating from spurious transcriptions, can be recruited to fulfill this task. <u>R</u>NAs responsive to <u>F</u>e (rrF) are indeed common in bacteria (5,93). The long-term evolution of some rrFs is expected to lead *in fine* to the birth of IsrR/RyhB functional analogs.

## DATA AVAILABILITY

Data generated or analyzed during this study are presented. Strains and plasmids are available from the corresponding author on request.

## ACKNOWLEDGEMENT

We are grateful to our colleagues, Sandy Gruss (INRAE, MICALIS) for critical reading of the manuscript, Cintia Gonzalez (UMR_S 1230) and Svetlana Chabelskaia (UMR_S 1230) for assistance with animal model experiments, Elise Borezée-Durant (INRAE, MICALIS) for the gift of staphylococcal strains, Aurélie Jaffrenou (I2BC) and Pierre Boudry (I2BC) for the gift of plasmids, Isabelle Hatin (I2BC) for the use of a flow cytometer, Olga Soutourina (I2BC) for access to an anaerobic chamber, Kam Pou Ha (I2BC) for the help with statistical analysis and Béatrice Py (LCB, UMR 7283) for her expertise to identify proteins containing iron-sulfur clusters in *S. aureus* presented in Supplementary Table S6. We thank our lab colleagues and Paul Aner for helpful discussions, technical help, and warm support. We acknowledge the high-throughput sequencing facility of I2BC for its sequencing and bioinformatics expertise and for its contribution to this study (http://www.i2bc.paris-saclay.fr), and the animal core facility ARCHE (ARCHE/SFR BIOSIT, Université Rennes 1, France) for animal experiments.

## AUTHOR CONTRIBUTIONS

P.B. conceived the project. R.H.C.T., W. L., M. B. and P.B. designed experiments and interpreted the data. V.B. and B.F. designed, performed the animal experiments, EMSAs and interpreted the data. M.P., C.V. and B.S. designed, performed experiments, and interpreted the data to determine RNA structures and interactions. R.H.C.T. and P.B. wrote the manuscript. R.H.C.T., B.F., M.P., B.S., and P.B. edited the paper.

