## Supplementary data for "sRNA-controlled iron sparing response in Staphylococci"

**Table S1. *Staphylococcus aureus* strains ... 2**

**Table S2. Plasmids ... 9**

**Table S3. Primers ... 13**

**Table S4. Fitness library composition ... 22**

**Table S5. IsrR sequences in *Staphylococcus* genus ... 23**

**Table S6. Proteins containing an Fe-S cluster in *S. aureus* ... 25**

**Table S7. IsrR functional analogs and their targets ... 26**

**Figure S1. Absence of IsrR is detrimental when iron is scarce ... 27**

**Figure S2. *isrR* complementation restores optimal growth in low-iron conditions ... 28**

**Figure S3. IsrR 5'/3'RACE mapping and secondary structure prediction ... 29**

**Figure S4. IsrR putative targets ... 30**

**Figure S5. IsrR putative targets involved in nitrate respiration pathway ... 31**

**Figure S6. Comparison of *fdhA* and *gltB2* mRNAs reactivity to 1M7 obtained in the presence/absence of IsrR ... 32**

**Figure S7. Comparison of *nasD* mRNA reactivity obtained to 1M7 in the presence/absence of IsrR with proposed interaction model ... 33**

**Figure S8. Electrophoretic mobility shift assay of IsrR in the presence of *fdhA* mRNA ... 34**

**Figure S10. Translational down-regulation of *narG* and *nasD* mRNAs by IsrR and CRR contribution ... 36**

**Figure S11. Hfq is not required for IsrR activity ... 37**

**REFERENCES ... 38**

**Table S1.** ***Staphylococcus aureus*** **strains**

| Name | Relevant genotype | Reference or construction |
|---|---|---|
| RN4220 | NCTC8325 derivative used for transformations with plasmids constructed in *E. coli pcnB*⁻ strain | (1) |
| 8325-4 | NCTC8325 derivative | (2) |
| HG003 | NCTC8325 *rsbU* and *tcaR* repaired | (3) |
| **Δ*sRNA* tagged mutants for libraries (Figure 1B)** | | |
| SAPhB618 | as HG003 Δ*rnaIII*::tag004 | (4) |
| SAPhB347 | as HG003 Δ*rsaOG*::tag009 | (4) |
| SAPhB349 | as HG003 Δ*rsaG*::tag011 | (4) |
| SAPhB368 | as HG003 Δ*teg147*::tag018 | (4) |
| SAPhB380 | as HG003 Δ*rsaB*::tag025 | (4) |
| SAPhB682 | as HG003 Δ*rsaD*::tag026 | (4) |
| SAPhB386 | as HG003 Δ*teg116*::tag030 | (4) |
| SAPhB397 | as HG003 Δ*sau85*::tag038 | (4) |
| SAPhB402 | as HG003 Δ*sau6353*::tag042 | (4) |
| SAPhB404 | as HG003 Δ*rsaE*::tag045 | (4) |
| SAPhB412 | as HG003 Δ*ssr42*::tag050 | (4) |
| SAPhB415 | as HG003 Δ*teg155*::tag053 | (4) |
| SAPhB960 | as HG003 Δ*sprF3*::tag070 | HG003 + pIM-*sprF3*::tag070 |
| SAPhB961 | as HG003 Δ*sprF3*::tag070 | HG003 + pIM-*sprF3*::tag070 |
| SAPhB962 | as HG003 Δ*sprF3*::tag070 | HG003 + pIM-*sprF3*::tag070 |
| SAPhB862 | as HG003 Δ*sRNA334*::tag073 | HG003 + pIM-*sRNA334*::tag073 |
| SAPhB863 | as HG003 Δ*sRNA334*::tag073 | HG003 + pIM-*sRNA334*::tag073 |
| SAPhB864 | as HG003 Δ*sRNA334*::tag073 | HG003 + pIM-*sRNA334*::tag073 |
| SAPhB943 | as HG003 Δ*rsaA*::tag075 | HG003 + pIM-*rsaA*::tag075 |
| SAPhB944 | as HG003 Δ*rsaA*::tag075 | HG003 + pIM-*rsaA*::tag075 |
| SAPhB945 | as HG003 Δ*rsaA*::tag075 | HG003 + pIM-*rsaA*::tag075 |
| SAPhB890 | as HG003 Δ*sau76*::tag076 | HG003 + pIM-*sau76*::tag076 |
| SAPhB891 | as HG003 Δ*sau76*::tag076 | HG003 + pIM-*sau76*::tag076 |
| SAPhB962 | as HG003 Δ*sau76*::tag076 | HG003 + pIM-*sau76*::tag076 |
| SAPhB883 | as HG003 Δ*rsaOI*::tag077 | HG003 + pIM-*rsaOI*::tag077 |

| | | |
|---|---|---|
| SAPhB884 | as HG003 Δ*rsaOI*::tag077 | HG003 + pIM-*rsaOI*::tag077 |
| SAPhB885 | as HG003 Δ*rsaOI*::tag077 | HG003 + pIM-*rsaOI*::tag077 |
| SAPhB865 | as HG003 Δ*teg16*::tag080 | HG003 + pIM-*teg16*::tag080 |
| SAPhB866 | as HG003 Δ*teg16*::tag080 | HG003 + pIM-*teg16*::tag080 |
| SAPhB867 | as HG003 Δ*teg16*::tag080 | HG003 + pIM-*teg16*::tag080 |
| SAPhB871 | as HG003 Δ*sRNA287*::tag085 | HG003 + pIM-*sRNA287*::tag085 |
| SAPhB872 | as HG003 Δ*sRNA287*::tag085 | HG003 + pIM-*sRNA287*::tag085 |
| SAPhB873 | as HG003 Δ*sRNA287*::tag085 | HG003 + pIM-*sRNA287*::tag085 |
| SAPhB874 | as HG003 Δ*sRNA71*::tag086 | HG003 + pIM-*sRNA71*::tag086 |
| SAPhB875 | as HG003 Δ*sRNA71*::tag086 | HG003 + pIM-*sRNA71*::tag086 |
| SAPhB876 | as HG003 Δ*sRNA71*::tag086 | HG003 + pIM-*sRNA71*::tag086 |
| SAPhB907 | as HG003 Δ*sRNA209*::tag093 | HG003 + pIM-*sRNA209*::tag093 |
| SAPhB908 | as HG003 Δ*sRNA209*::tag093 | HG003 + pIM-*sRNA209*::tag093 |
| SAPhB909 | as HG003 Δ*sRNA209*::tag093 | HG003 + pIM-*sRNA209*::tag093 |
| SAPhB899 | as HG003 Δ*teg106*::tag095 | HG003 + pIM-*teg106*::tag095 |
| SAPhB900 | as HG003 Δ*teg106*::tag095 | HG003 + pIM-*teg106*::tag095 |
| SAPhB946 | as HG003 Δ*teg106*::tag095 | HG003 + pIM-*teg106*::tag095 |
| SAPhB921 | as HG003 Δ*sRNA260*::tag096 | HG003 + pIM-*sRNA260*::tag096 |
| SAPhB922 | as HG003 Δ*sRNA260*::tag096 | HG003 + pIM-*sRNA260*::tag096 |
| SAPhB947 | as HG003 Δ*sRNA260*::tag096 | HG003 + pIM-*sRNA260*::tag096 |
| SAPhB910 | as HG003 Δ*sRNA345*::tag097 | HG003 + pIM-*sRNA345*::tag097 |
| SAPhB911 | as HG003 Δ*sRNA345*::tag097 | HG003 + pIM-*sRNA345*::tag097 |
| SAPhB912 | as HG003 Δ*sRNA345*::tag097 | HG003 + pIM-*sRNA345*::tag097 |
| SAPhB932 | as HG003 Δ*ncRNA2*::tag099 | HG003 + pIM-*ncRNA2*::tag099 |
| SAPhB933 | as HG003 Δ*ncRNA2*::tag099 | HG003 + pIM-*ncRNA2*::tag099 |
| SAPhB934 | as HG003 Δ*ncRNA2*::tag099 | HG003 + pIM-*ncRNA2*::tag099 |
| SAPhB940 | as HG003 Δ*ncRNA3*::tag100 | HG003 + pIM-*ncRNA3*::tag100 |
| SAPhB941 | as HG003 Δ*ncRNA3*::tag100 | HG003 + pIM-*ncRNA3*::tag100 |
| SAPhB942 | as HG003 Δ*ncRNA3*::tag100 | HG003 + pIM-*ncRNA3*::tag100 |
| SAPhB954 | as HG003 Δ*ssrS*::tag107 | HG003 + pIM-*ssrS*::tag107 |
| SAPhB955 | as HG003 Δ*ssrS*::tag107 | HG003 + pIM-*ssrS*::tag107 |
| SAPhB956 | as HG003 Δ*ssrS*::tag107 | HG003 + pIM-*ssrS*::tag107 |
| SAPhB1006 | as HG003 Δ*sprF1*::tag110 | HG003 + pIM-*sprF1*::tag110 |

| | | |
|---|---|---|
| SAPhB1007 | as HG003 Δ*sprF1*::tag110 | HG003 + pIM-*sprF1*::tag110 |
| SAPhB1008 | as HG003 Δ*sprF1*::tag110 | HG003 + pIM-*sprF1*::tag110 |
| SAPhB974 | as HG003 Δ*sprX2*::tag111 | HG003 + pIM-*sprX2*::tag111 |
| SAPhB975 | as HG003 Δ*sprX2*::tag111 | HG003 + pIM-*sprX2*::tag111 |
| SAPhB997 | as HG003 Δ*sprX2*::tag111 | HG003 + pIM-*sprX2*::tag111 |
| SAPhB978 | as HG003 Δ*sprY2*::tag112 | HG003 + pIM-*sprY2*::tag112 |
| SAPhB979 | as HG003 Δ*sprY2*::tag112 | HG003 + pIM-*sprY2*::tag112 |
| SAPhB980 | as HG003 Δ*sprY2*::tag112 | HG003 + pIM-*sprY2*::tag112 |
| SAPhB957 | as HG003 Δ*sprY3*::tag113 | HG003 + pIM-*sprY3*::tag113 |
| SAPhB958 | as HG003 Δ*sprY3*::tag113 | HG003 + pIM-*sprY3*::tag113 |
| SAPhB959 | as HG003 Δ*sprY3*::tag113 | HG003 + pIM-*sprY3*::tag113 |
| SAPhB901 | as HG003 Δ*sau41*::Tag115 | HG003 + pIM-*sau41*::Tag115 |
| SAPhB902 | as HG003 Δ*sau41*::Tag115 | HG003 + pIM-*sau41*::Tag115 |
| SAPhB903 | as HG003 Δ*sau41*::Tag115 | HG003 + pIM-*sau41*::Tag115 |
| SAPhB948 | as HG003 Δ*sau5949*::tag117 | HG003 + pIM-*sau5949*::tag117 |
| SAPhB949 | as HG003 Δ*sau5949*::tag117 | HG003 + pIM-*sau5949*::tag117 |
| SAPhB950 | as HG003 Δ*sau5949*::tag117 | HG003 + pIM-*sau5949*::tag117 |
| SAPhB966 | as HG003 Δ*sprF2*::tag118 | HG003 + pIM-*sprF2*::tag118 |
| SAPhB967 | as HG003 Δ*sprF2*::tag118 | HG003 + pIM-*sprF2*::tag118 |
| SAPhB998 | as HG003 Δ*sprF2*::tag118 | HG003 + pIM-*sprF2*::tag118 |
| SAPhB1031 | as HG003 Δ*sprB*::tag121 | HG003 + pIM-*sprB*::tag121 |
| SAPhB1032 | as HG003 Δ*sprB*::tag121 | HG003 + pIM-*sprB*::tag121 |
| SAPhB1033 | as HG003 Δ*sprB*::tag121 | HG003 + pIM-*sprB*::tag121 |
| SAPhB1242 | as HG003 Δ*rsaC*::tag133 | HG003 + pIM-*rsaC*::tag133 |
| SAPhB1243 | as HG003 Δ*rsaC*::tag133 | HG003 + pIM-*rsaC*::tag133 |
| SAPhB1244 | as HG003 Δ*rsaC*::tag133 | HG003 + pIM-*rsaC*::tag133 |
| SAPhB1234 | as HG003 Δ*S204*::tag134 | HG003 + pIM-*S204*::tag134 |
| SAPhB1235 | as HG003 Δ*S204*::tag134 | HG003 + pIM-*S204*::tag134 |
| SAPhB1236 | as HG003 Δ*S204*::tag134 | HG003 + pIM-*S204*::tag134 |
| SAPhB1231 | as HG003 Δ*isrR*::tag135 | HG003 + pIM-*S596*::tag135 |
| SAPhB1232 | as HG003 Δ*isrR*::tag135 | HG003 + pIM-*S596*::tag135 |
| SAPhB1233 | as HG003 Δ*isrR*::tag135 | HG003 + pIM-*S596*::tag135 |
| SAPhB1239 | as HG003 Δ*S808*::tag137 | HG003 + pIM-*S808*::tag137 |

| | | |
|---|---|---|
| SAPhB1240 | as HG003 Δ*S808*::tag137 | HG003 + pIM-*S808*::tag137 |
| SAPhB1241 | as HG003 Δ*S808*::tag137 | HG003 + pIM-*S808*::tag137 |
| SAPhB1015 | *locus3*::tag139 | HG003 + pIM-*locus3*::tag139 |
| SAPhB1016 | *locus3*::tag139 | HG003 + pIM-*locus3*::tag139 |
| SAPhB1017 | *locus3*::tag139 | HG003 + pIM-*locus3*::tag139 |
| SAPhB1012 | *locus2*::tag140 | HG003 + pIM-*locus2*::tag140 |
| SAPhB1013 | *locus2*::tag140 | HG003 + pIM-*locus2*::tag140 |
| SAPhB1014 | *locus2*::tag140 | HG003 + pIM-*locus2*::tag140 |
| SAPhB1009 | *locus1*::tag141 | HG003 + pIM-*locus1*::tag141 |
| SAPhB1010 | *locus1*::tag141 | HG003 + pIM-*locus1*::tag141 |
| SAPhB1011 | *locus1*::tag141 | HG003 + pIM-*locus1*::tag141 |
| SAPhB1018 | as HG003 Δ*sau5971*::tag142 | HG003 + pIM-*sau5971*::tag142 |
| SAPhB1019 | as HG003 Δ*sau5971*::tag142 | HG003 + pIM-*sau5971*::tag142 |
| SAPhB1020 | as HG003 Δ*sau5971*::tag142 | HG003 + pIM-*sau5971*::tag142 |
| SAPhB976 | as HG003 Δ*sprA1*::tag144 | HG003 + pIM-*sprA1*::tag144 |
| SAPhB977 | as HG003 Δ*sprA1*::tag144 | HG003 + pIM-*sprA1*::tag144 |
| SAPhB996 | as HG003 Δ*sprA1*::tag144 | HG003 + pIM-*sprA1*::tag144 |
| SAPhB1027 | as HG003 Δ*sprX2*::tag145 Δ*sprX1*::tag149 | SAPhB976 + pIM-*sprX2*::tag145 |
| SAPhB1028 | as HG003 Δ*sprX2*::tag145 Δ*sprX1*::tag149 | SAPhB976 + pIM-*sprX2*::tag145 |
| SAPhB1029 | as HG003 Δ*sprX2*::tag145 Δ*sprX1*::tag149 | SAPhB976 + pIM-*sprX2*::tag145 |
| SAPhB1003 | as HG003 Δ*sprX1*::tag146 | HG003 + pIM-*sprX1*::tag146 |
| SAPhB1004 | as HG003 Δ*sprX1*::tag146 | HG003 + pIM-*sprX1*::tag146 |
| SAPhB1005 | as HG003 Δ*sprX1*::tag146 | HG003 + pIM-*sprX1*::tag146 |
| SAPhB971 | as HG003 Δ*rsaH*::tag147 | HG003 + pIM-*rsaH*::tag147 |
| SAPhB972 | as HG003 Δ*rsaH*::tag147 | HG003 + pIM-*rsaH*::tag147 |
| SAPhB973 | as HG003 Δ*rsaH*::tag147 | HG003 + pIM-*rsaH*::tag147 |
| SAPhB1021 | as HG003 Δ*sprY1*::tag148 | HG003 + pIM-*sprY1*::tag148 |
| SAPhB1022 | as HG003 Δ*sprY1*::tag148 | HG003 + pIM-*sprY1*::tag148 |
| SAPhB1023 | as HG003 Δ*sprY1*::tag148 | HG003 + pIM-*sprY1*::tag148 |
| **IsrR complementation studies (Figure 1C and S2)** | | |
| SAPhB1372 | as HG003 Δ*isrR*::tag135 pCN38 | SAPhB1231 + pCN38 |
| SAPhB1373 | as HG003 Δ*isrR*::tag135 pCN38-IsrR | SAPhB1231 + pCN38-IsrR |
| SAPhB1500 | as HG003 Δ*isrR*::tag135 *locus2*::*isrR*$^{+}$ | SAPhB1231 + pIM-*locus2*::isrR$^{+}$ |

| | | |
|---|---|---|
| SAPhB1502 | as HG003 Δ*isrR*::tag135 *locus3*::*isrR*$^{+}$ | SAPhB1231 + pIM-*locus3*::isrR$^{+}$ |
| **Fur regulation (Figure 2)** | | |
| MJH010 | as NCTC8325-4 Δ*fur*::*tetR* | (5) |
| SAPhB1542 | as HG003 Δ*fur*::*tetR* | HG003 + ϕ80 on MJH010 |
| SAPhB1558 | as NCTC8325-4 $pP_{isrR}$ | NCTC8325-4 + $pP_{isrR}$ |
| SAPhB1550 | as NCTC8325-4 $pP_{isrR}$::*gfp* | NCTC8325-4 + $pP_{isrR}$::*gfp* |
| SAPhB1552 | as NCTC8325-4 $pP_{isrR1}$::*gfp* | NCTC8325-4 + $pP_{isrR1}$::*gfp* |
| SAPhB1554 | as NCTC8325-4 $pP_{isrR2}$::*gfp* | NCTC8325-4 + $pP_{isrR2}$::*gfp* |
| SAPhB1556 | as NCTC8325-4 $pP_{isrR1\&2}$::*gfp* | NCTC8325-4 + $pP_{isrR1\&2}$::*gfp* |
| **pRMC2-IsrR derivatives** | | |
| SAPhB1801 | as HG003 pRMC2ΔR | HG003 + pRMC2ΔR |
| SAPhB1517 | as HG003 pRMC2ΔR-IsrR | HG003 + pRMC2ΔR-IsrR |
| SAPhB1618 | as HG003 Δ*isrR*::tag135 pRMC2ΔR | SAPhB1231 + pRMC2ΔR |
| SAPhB1519 | as HG003 Δ*isrR*::tag135 pRMC2ΔR-IsrR | SAPhB1231 + pRMC2ΔR-IsrR |
| SAPhB1568 | as HG003 Δ*isrR*::tag135 pRMC2ΔR-IsrRΔCRR1 | SAPhB1231 + pRMC2ΔR-IsrRΔCRR1 |
| SAPhB1674 | as HG003 Δ*isrR*::tag135 pRMC2ΔR-IsrRΔCRR2 | SAPhB1231 + pRMC2ΔR-IsrRΔCRR2 |
| SAPhB1703 | as HG003 Δ*isrR*::tag135 pRMC2ΔR-IsrRΔCRR3 | SAPhB1231 + pRMC2ΔR-IsrRΔCRR3 |
| **IsrR/*gltB2* mRNA pairing reporter assay (Figure 7)** | | |
| SAPhB1628 | as HG003 Δ*isrR*::tag135 pRMC2ΔR p5'GltB2-GFP | SAPhB1618 + p5'GltB2-GFP |
| SAPhB1598 | as HG003 Δ*isrR*::tag135 pRMC2ΔR-IsrR p5'GltB2-GFP | SAPhB1519 + p5'GltB2-GFP |
| SAPhB1600 | as HG003 Δ*isrR*::tag135 pRMC2ΔR-IsrRΔCRR1 p5'GltB2-GFP | SAPhB1568 + p5'GltB2-GFP |
| SAPhB1717 | as HG003 Δ*isrR*::tag135 pRMC2ΔR-IsrR ΔCRR2 p5'GltB2-GFP | SAPhB1674 + p5'GltB2-GFP |
| SAPhB1721 | as HG003 Δ*isrR*::tag135 pRMC2ΔR-IsrR ΔCRR3 p5'GltB2-GFP | SAPhB1703 + p5'GltB2-GFP |
| **IsrR/*fdhA* mRNA pairing reporter assay (Figure 7)** | | |
| SAPhB1745 | as HG003 Δ*isrR*::tag135 pRMC2ΔR p5'FdhA-GFP | SAPhB1618 + p5'FdhA-GFP |
| SAPhB1747 | as HG003 Δ*isrR*::tag135 pRMC2ΔR-IsrR p5'FdhA-GFP | SAPhB1519 + p5'FdhA-GFP |

| | | |
|---|---|---|
| SAPhB1749 | as HG003 Δ*isrR*::tag135 pRMC2ΔR-IsrRΔCRR1 p5'FdhA-GFP | SAPhB1568 + p5'FdhA-GFP |
| SAPhB1751 | as HG003 Δ*isrR*::tag135 pRMC2ΔR-IsrRΔCRR2 p5'FdhA-GFP | SAPhB1674 + p5'FdhA-GFP |
| SAPhB1753 | as HG003 Δ*isrR*::tag135 pRMC2ΔR-IsrRΔCRR3 p5'FdhA-GFP | SAPhB1703 + p5'FdhA-GFP |
| **IsrR/*narG* mRNA pairing reporter assay (Figure S10)** | | |
| SAPhB1624 | as HG003 Δ*isrR*::tag135 pRMC2ΔR p5'NarG-GFP | SAPhB1618 + p5'NarG-GFP |
| SAPhB1590 | as HG003 Δ*isrR*::tag135 pRMC2ΔR-IsrR p5'NarG-GFP | SAPhB1519 + p5'NarG-GFP |
| SAPhB1592 | as HG003 Δ*isrR*::tag135 pRMC2ΔR-IsrRΔCRR1 p5'NarG-GFP | SAPhB1568 + p5'NarG-GFP |
| SAPhB1741 | as HG003 Δ*isrR*::tag135 pRMC2ΔR-IsrRΔCRR2 p5'NarG-GFP | SAPhB1674 + p5'NarG-GFP |
| SAPhB1743 | as HG003 Δ*isrR*::tag135 pRMC2ΔR-IsrRΔCRR3 p5'NarG-GFP | SAPhB1703 + p5'NarG-GFP |
| **IsrR/*nasD* mRNA pairing reporter assay (Figure S10)** | | |
| SAPhB1626 | as HG003 Δ*isrR*::tag135 pRMC2ΔR p5'NasD-GFP | SAPhB1618 + p5'NasD-GFP |
| SAPhB1594 | as HG003 Δ*isrR*::tag135 pRMC2ΔR-IsrR p5'NasD-GFP | SAPhB1519 + p5'NasD-GFP |
| SAPhB1596 | as HG003 Δ*isrR*::tag135 pRMC2ΔR-IsrRΔCRR1 p5'NasD-GFP | SAPhB1568 + p5'NasD-GFP |
| SAPhB1781 | as HG003 Δ*isrR*::tag135 pRMC2ΔR-IsrRΔCRR2 p5'NasD-GFP | SAPhB1674 + p5'NasD-GFP |
| SAPhB1783 | as HG003 Δ*isrR*::tag135 pRMC2ΔR-IsrRΔCRR3 p5'NasD-GFP | SAPhB1703 + p5'NasD-GFP |
| ***hfq* derivatives (Figure S11)** | | |
| SAPhB1024 | as HG003 Δ*hfq*::tag143 | (4) |
| SAPhB1709 | as HG003 Δ*hfq*::tag143 pRMC2ΔR | SAPhB1024 + pRMC2ΔR |
| SAPhB1711 | as HG003 Δ*hfq*::tag143 pRMC2ΔR-IsrR | SAPhB1024 + pRMC2ΔR-IsrR |
| SAPhB1807 | as HG003 Δ*hfq*::tag143 pRMC2ΔR p5'FdhA-GFP | SAPhB1709+ p5'FdhA-GFP |
| SAPhB1809 | as HG003 Δ*hfq*::tag143 pRMC2ΔR-IsrR p5'FdhA-GFP | SAPhB1711+ p5'FdhA-GFP |

| | | |
|---|---|---|
| SAPhB1811 | as HG003 Δ*hfq*::tag143 pRMC2ΔR p5'NarG-GFP | SAPhB1709 + p5'NarG-GFP |
| SAPhB1813 | as HG003 Δ*hfq*::tag143 pRMC2ΔR-IsrR p5'NarG-GFP | SAPhB1711 + p5'NarG-GFP |
| SAPhB1815 | as HG003 Δ*hfq*::tag143 pRMC2ΔR p5'NasD-GFP | SAPhB1709 + p5'NasD-GFP |
| SAPhB1817 | as HG003 Δ*hfq*::tag143 pRMC2ΔR-IsrR p5'NasD-GFP | SAPhB1711 + p5'NasD-GFP |
| SAPhB1819 | as HG003 Δ*hfq*::tag143 pRMC2ΔR p5'GltB2-GFP | SAPhB1709 + p5'GltB2-GFP |
| SAPhB1821 | as HG003 Δ*hfq*::tag143 pRMC2ΔR-IsrR p5'GltB2-GFP | SAPhB1711 + p5'GltB2-GFP |
| SAPhB2056 | as HG003 p5'FdhA-GFP | HG003 + p5'FdhA-GFP |
| SAPhB2057 | as HG003 Δ*isrR*::tag135 p5'FdhA-GFP | SAPhB1231 + p5'FdhA-GFP |
| SAPhB2058 | as HG003 Δ*hfq*::tag143 p5'FdhA-GFP | SAPhB1024 + p5'FdhA-GFP |

**Table S2. Plasmids**

| Name | Relevant properties | Reference /Construction* |
|---|---|---|
| **Mutant library constructions** | | |
| pIMAY | Shuttle *rep*(Ts) vector in *S. aureus* | (6) |
| pIM-*sprF3*::tag070 | *sprF3* replacement by tag070 | up: 2147/2148; dw: 2149/2150 |
| pIM-*sRNA334*::tag07 | *sRNA334* replacement by tag073 | up:1959/1960; dw; 1961/1962 |
| pIM-*rsaA*::tag075 | *rsaA* replacement by tag075 | up: 1928/1929; dw: 1930/1931 |
| pIM-*sau76*::tag076 | *sau76* replacement by tag076 | up: 1836/1837; dw: 1838/1839 |
| pIM-*rsaOI*::tag077 | *rsaOI* replacement by tag077 | up:1840/1841; dw: 1842/1843 |
| pIM-*teg16*::tag080 | *teg16* replacement by tag080 | up: 1931/1932; dw: 1933/1934 |
| pIM-*sRNA287*::tag085 | *sRNA287* replacement by tag085 | up: 1952/1953; dw: 1954/1955 |
| pIM-*sRNA71*::tag086 | *sRNA71* replacement by tag086 | up: 1910/1911; dw: 1912/1913 |
| pIM-*sRNA209*::tag093 | *sRNA209* replacement by tag093 | up:1987/1988; dw: 1989/1990 |
| pIM-*teg106*::tag095 | *teg106* replacement by tag095 | up2001/2002; dw: 2003/2004 |
| pIM-*sRNA260*::tag096 | *sRNA260* replacement by t | up: 2008/2009; dw: 2009/2010 |
| pIM-*sRNA345*::tag097 | *sRNA345* replacement by tag097 | up: 2015/2016; dw: 2017/2018 |
| pIM-*ncRNA2*::tag099 | *ncRNA2* replacement by tag099 | up: 2031/2032; dw: 2033/2034 |
| pIM-*ncRNA3*::tag100 | *ncRNA3* replacement by tag100 | up: 2038/2039; dw: 2040/2041 |
| pIM-*ssrS*::tag107 | *ssrS* replacement by tag107 | up: 2019/2020; dw: 2021/2022 |
| pIM-*sprF1*::tag110 | *sprF1* replacement by tag110 | up: 2161/2162; dw: 2163/2164 |
| pIM-*sprX2*::tag111 | *sprX2* replacement by tag111 | up: 2189/2190; dw: 2191/2192 |

| | | |
|---|---|---|
| pIM-*sprY2*::tag112 | *sprY2* replacement by tag112 | up: 2196/2197; dw: 2198/2199 |
| pIM-*sprY3*::tag113 | *sprY3* replacement by tag113 | up: 2168/2169; dw: 2170/2171 |
| pIM-*sau41*::Tag115 | *sau41* replacement by Tag115 | up:1966/1967; dw; 1968/1969 |
| pIM-*sau5949*::tag117 | *sau5949* replacement by tag117 | up: 2066/2067; dw: 2068/2069 |
| pIM-*sprF2*::tag118 | *sprF2* replacement by tag118 | up: 2026/2027; dw: 2028/2029 |
| pIM-*rsaC*::tag133 | *rsaC* replacement by tag133 | up: 2395/2396; dw: 2397/2398 |
| pIM-*S204*::tag134 | *S204* replacement by tag134 | up: 2387/2388; dw: 2389/2390 |
| pIM-*S596*::tag135 | *S596* replacement by tag135 | up: 2379/2380; dw: 2381/2382 |
| pIM-*S808*::tag137 | *S808* replacement by tag137 | up: 2363/2364; dw: 2365/2366 |
| pIM-*locus3*::tag139 | *locus3* replacement by tag139 | up: 2234/2235; dw: 2236/2237 |
| pIM-*locus2*::tag140 | *locus2* replacement by tag140 | up: 2222/2223; dw: 2224/2225 |
| pIM-locus1::tag141 | locus1 replacement by tag141 | up: 2228/2229; dw: 2230/2231 |
| pIM-*sau5971*::tag142 | *sau5971* replacement by tag142 | up: 2215/2216; dw: 2217/2218 |
| pIM-*sprA1*::tag144 | *sprA1* replacement by tag144 | up: 2140/2141; dw: 2142/2143 |
| pIM-*sprX2*::tag145 | *sprX2* replacement by tag145 | up: 2189/2190; dw: 2191/2192 |
| pIM-sprX1::tag149 | sprX1 replacement by tag149 | up: 2175/2176; dw: 2177/2178 |
| pIM-*sprX1*::tag146 | *sprX1* replacement by tag146 | up: 2175/2176; dw: 2177/2178 |
| pIM-*rsaH*::tag147 | *rsaH* replacement by tag147 | up: 2201/2202; dw: 2203/2204 |
| pIM-*sprY1*::tag148 | *sprY1* replacement by tag148 | up: 2182/2183; dw: 2184/2185 |

| | | |
|---|---|---|
| ***isrR* complementation and toxicity** | | |
| pCN38 | Shuttle vector, pT181 replicon, $Cm^R$ | (7) |
| pCN38-IsrR | *isrR* under the control of its endogenous promoter | 1489/1490 on pCN38 + 2343/2344 on HG003 |
| pIM-locus2 | SAOUHSC_03030-locus2-SAOUHSC_03031 region | 1536/1537 on pIMAY + 2228/2231 on HG003 |
| pIM-locus2::$isrR^+$ | For *isrR* insertion at HG003 locus2 | 2507/2508 on pIM-locus2 + 2511/2512 on HG003 |
| pIM-locus3 | SAOUHSC_01263-locus3-SAOUHSC_01264 region | 1536/1537 + pIM-locus3 + 2234/2237 on HG003 |
| pIM-locus3::$isrR^+$ | For *isrR* insertion at HG003 locus3 | 2509/2510 pIM-locus3 + 2511/2512 HG003 |
| **Fur-dependent isrR regulation** | | |
| $pP_{isrR}$::GFP | Translational fusion between isrR promoter region and s*gfp* | 2546/ 2547 on pCN38-isrR + 2548/2549 on pCM11 |
| $pP_{isrR1}$::GFP | $pP_{isrR}$::GFP with the *isrR* Fur motif 1 mutated. | 2573/2574 on $pP_{isrR}$::GFP |
| $pP_{isrR2}$::GFP | $pP_{isrR}$::GFP with the *isrR* Fur motif 2 mutated. | 2575/2576 on $pP_{isrR}$::GFP |
| $pP_{isrR1\&2}$::GFP | $pP_{isrR}$::GFP with the *isrR* Fur motif 1 & 2 mutated. | 2575/2576 on $pP_{isrR1}$::GFP |
| **Constitutive expression of isrR and mutated derivatives** | | |
| pRMC2 | Anhydrotetracycline (aTc) inducible promoter | (8) |
| pRMC2ΔR | Deletion of *tetR* from pRMC2 | 2538/2539 on pRMC2 |
| pRMC2-IsrR | Inducible expression of *IsrR* | 856/918 on pRMC2 with + PCR 2499/2500 on HG003 with |
| pRMC2ΔR-IsrR | Deletion of *tetR* from pRMC2-IsrR: Constitutive expression of *isrR* | 2538/2539 on pRMC2-IsrR |
| pRMC2ΔR-IsrRΔCRR1 | Constitutive expression of *isrR*ΔCRR1 | 2605/2606 on pRMC2ΔR-IsrR |
| pRMC2ΔR-IsrRΔCRR2 | Constitutive expression of *isr*RΔCRR2 | 2631/2632 on pRMC2ΔR-IsrR |
| pRMC2ΔR-IsrRΔCRR3 | Constitutive expression of *isrR*ΔCRR3 | 2667/2668 on pRMC2ΔR-IsrR |

| | | |
|---|---|---|
| **Targets 5'UTR in translational fusion with *gfp*** | | |
| pCN34 | Shuttle vector, pT181 replicon, Km$^{R}$ | (7) |
| pCM11 | Promoter-less s*gfp* transcriptional reporter | (9) |
| pECTO | pMAD2 derivative to integrate DNA sequences between SAOUHSC_00278 and SAOUHSC_00279 | Laboratory collection https://www.addgene.org/ |
| pCN34-*gfp* | *rrnB* terminator from pECTO and *sarA* promoter, *sod* RBS and s*gfp* from pCM11 | 2554/2555 on pECTO + 2552/2553 on pCM11 + 2534/2535 on pCN34 |
| p5'GltB2-GFP | Translational fusion between *gltB2* 5'UTR and *gfp*, under the control of $P_{sarA}$ ($P_{sarA}$5'UTR::*gltB2*::*gfp*) | 2600/2601 on pCN34-*gfp* + 2621/2622 on HG003 |
| p5'FdhA-GFP | Translational fusion between *fdhA* 5'UTR and *gfp*, under the control of $P_{sarA}$ ($P_{sarA}$5'UTR::*fdhA*::*gfp*) | 2600/2601 on pCN34-*gfp* + 2694/2695 on HG003 |
| p5'NarG-GFP | Translational fusion between *narG* 5'UTR and *gfp*, under the control of $P_{sarA}$ ($P_{sarA}$5'UTR::*narG*::*gfp*) | 2600/2601 on pCN34-*gfp* + 2617/2618 on HG003 |
| p5'NasD-GFP | Translational fusion between *nasD* 5'UTR and *gfp*, under the control of $P_{sarA}$ ($P_{sarA}$5'UTR::*nasD*::*gfp*) | 2600/2601 on pCN34-*gfp* + 2619/2620 on HG003 |
| ***hfq* inactivation** | | |
| pIM-hfq::tag143 | *hfq* replacement by tag143 | up: 2208/2209; dw: 2210/2211 |

* #/# indicates primer couples used for PCR amplifications. For primer sequences, see Table S3. Plasmids were constructed by isothermal assembly of PCR product(s) (10). pIM plasmids: pIMAY derivatives for *S. aureus* chromosomal modifications were constructed by the assembly of PCR products from i) pIMAY (amplified with primers 1536 and 1537), ii) upstream and iii) downstream regions of each HG003 modified regions. Primers used for the amplifications these regions (about 0.8 to 1kb) are indicated, "up" and "dw" in the table, respectively. iv) Deleted genes were replaced by specific tag sequences amplified with primers 1870 and 1871 from a partially random primer, as described (4). For these, the assembly mixt contained an additional PCR product corresponding to amplified DNA tag sequences.

**Table S3. Primers**

| Name | Description, PCR amplified region* | Sequence |
|---|---|---|
| 856 | pRMC2 F | GGTACCGTTAACAGATCTGAG |
| 918 | pRMC2 R | GCTTATTTTAATTATACTCTATCAATGATAGAG |
| 1489 | pCN38 amplification-F | CAGTTGCGCAGCCTGAATGG |
| 1490 | pCN38 amplification-R | CCTCTAGAGTCGACCTGCAG |
| 1536 | pIMAY F | GGTACCCAGCTTTTGTTCCCTTTAGTGAGG |
| 1537 | pIMAY R | GAGCTCCAATTCGCCCTATAGTGAGTCG |
| 1828 | pIMAY-Up-RsaA-F | CGACTCACTATAGGGCGAATTGGAGCTCAGTTGCCAAGTCACCTGTTG |
| 1829 | pIMAY-Up-RsaA-R | GCGTATGGACCTAGGTATATCTCTATACAATTTTTGTAATGGTTAACT |
| 1830 | pIMAY-Down-RsaA-F | ACCCCACAACCTAGGTATATCTCGGGTACACTTTGCTATGAG |
| 1831 | pIMAY-Down-RsaA-R | CCTCACTAAAGGGAACAAAAGCTGGGTACCGCGATGCACTTGTCACTGAA |
| 1836 | pIMAY-Up-sau76-F | CGACTCACTATAGGGCGAATTGGAGCTCGGTATCCTAGACTACCTGCTAA |
| 1837 | pIMAY-Up-sau76-R | GCGTATGGACCTAGGTATATTAATGTATTATCAATAACAAAGTACA |
| 1838 | pIMAY-Down-sau76-F | ACCCCACAACCTAGGTATATATAAACGAAAAATTCCAAGCTTAAACC |
| 1839 | pIMAY-Down-sau76-R | CCTCACTAAAGGGAACAAAAGCTGGGTACCAAACATGAGTCAAGCAGCCG |
| 1840 | pIMAY -Up-RsaOI-F | CGACTCACTATAGGGCGAATTGGAGCTCGCAACACAACCAGAAAGAGATAAC |
| 1841 | pIMAY -Up-RsaOI-R | GCGTATGGACCTAGGTATATCTTTATTACGGCTAATTACAGTTCTCAA |
| 1842 | pIMAY -Down-RsaOI-F | ACCCCACAACCTAGGTATATTACAGTATCAAATTTATCTAGGGC |
| 1843 | pIMAY -Down-RsaOI-R | CCTCACTAAAGGGAACAAAAGCTGGGTACCTGCAGCTTATCTCCACTGCT |
| 1870 | Tag amplification-R | GGTCTCTGAGATCCATACGCAGCTATGCAAT |
| 1871 | Tag amplification-F | GGTCTCATGTGTTGTGGGGTACAGCAATGAC |
| 1910 | pIMAY-Up-sRNA71-F | CGACTCACTATAGGGCGAATTGGAGCTCTGGTGGTAAGTCTGTTGAAAAGA |
| 1911 | pIMAY-Up-sRNA71-R | GCGTATGGACCTAGGTATATGTAACACATCACTTGATTAAAGACAATAC |
| 1912 | pIMAY-Down-sRNA71-F | ACCCCACAACCTAGGTATATGTCAAATACTCGCTTTTTTATTTCC |
| 1913 | pIMAY-Down-sRNA71-R | CCTCACTAAAGGGAACAAAAGCTGGGTACCCCCATTCCCAAATTTGATGTGC |

| | | |
|---|---|---|
| 1931 | pIMAY-Up-Teg16-F | CGACTCACTATAGGGCGAATTGGAGCTCTGAACCAGGACCTTCAGCAA |
| 1932 | pIMAY-Up-Teg16-R | GCGTATGGACCTAGGTATATAATTTACTATATCTGCTTTAGTATGTCAAC |
| 1933 | pIMAY-Down-Teg16-F | ACCCCACAACCTAGGTATATGTTTGAATGGACTTGTAAACGT |
| 1934 | pIMAY-Down-Teg16-R | CCTCACTAAAGGGAACAAAAGCTGGGTACCGCGAATCATTTCTCGTCGCT |
| 1952 | pIMAY-Up-sRNA287-F | CGACTCACTATAGGGCGAATTGGAGCTCAATTCATGGCGGTTGTGGTG |
| 1953 | pIMAY-Up-sRNA287-R | GCGTATGGACCTAGGTATATCGATTAGGTCATGCAGATGT |
| 1954 | pIMAY-Down-sRNA287-F | ACCCCACAACCTAGGTATATTATAGAACTGCGAACAGGTG |
| 1955 | pIMAY-Down-sRNA287-R | CCTCACTAAAGGGAACAAAAGCTGGGTACCAGACACAGGCAAAATGAGTT |
| 1959 | pIMAY-Up-sRNA334-F | CGACTCACTATAGGGCGAATTGGAGCTCAGCAAATATCTCTTCTCCAACCA |
| 1960 | pIMAY-Up-sRNA334-R | GCGTATGGACCTAGGTATATTCTACTTTAAATTATCATCTCCATACTATT |
| 1961 | pIMAY-Down-sRNA334-F | ACCCCACAACCTAGGTATATGTTAAACCTTAAAACAAGAAATATTATTCA |
| 1962 | pIMAY-Down-sRNA334-R | CCTCACTAAAGGGAACAAAAGCTGGGTACCAGCATGGTTTATCATTGGCTCA |
| 1966 | pIMAY-Up-Sau41-F | CGACTCACTATAGGGCGAATTGGAGCTCTGCTAAAGATGCAAACGACGT |
| 1967 | pIMAY-Up-Sau41-R | GCGTATGGACCTAGGTATATTTCATTGTACATAGTTATCTTGTGCGT |
| 1968 | pIMAY-Down-Sau41-F | ACCCCACAACCTAGGTATATACTTAAAATTCTCAGGCCACTATACC |
| 1969 | pIMAY-Down-Sau41-R | CCTCACTAAAGGGAACAAAAGCTGGGTACCTCTGAAGCGCAACAAACACA |
| 1987 | pIMAY-Up-sRNA209-F | CGACTCACTATAGGGCGAATTGGAGCTCAAACCCACACCGTTAGCAAC |
| 1988 | pIMAY-Up-sRNA209-R | GCGTATGGACCTAGGTATATTGACGCATCATACTATATTACTGAAATTC |
| 1989 | pIMAY-Down-sRNA209-F | ACCCCACAACCTAGGTATATAAATAACCACGTCCATCGAGA |
| 1990 | pIMAY-Down-sRNA209-R | CCTCACTAAAGGGAACAAAAGCTGGGTACCTGCGTCATAATTCCCACAAGG |
| 2001 | pIMAY-Up-Teg106-F | CGACTCACTATAGGGCGAATTGGAGCTCTCTGGTAGGACTATTGAATTTGCA |
| 2002 | pIMAY-Up-Teg106-R | GCGTATGGACCTAGGTATATCCATTCACCATATGATTTTTATTAATAGTT |
| 2003 | pIMAY-Down-Teg106-F | ACCCCACAACCTAGGTATATCGTCTTGAAATGCTCCCTTCA |
| 2004 | pIMAY-Down-Teg106-R | CCTCACTAAAGGGAACAAAAGCTGGGTACCTCGCCATCTTCACCAAGTTC |
| 2008 | pIMAY-Up-sRNA260-F | CGACTCACTATAGGGCGAATTGGAGCTCAACGCAACCAAGTGATGTTG |

| | | |
|---|---|---|
| 2009 | pIMAY-Up-sRNA260-R | GCGTATGGACCTAGGTATATCATAACAAAACTCCTAATGTACTAGTTTAG |
| 2010 | pIMAY-Down-sRNA260-F | ACCCCACAACCTAGGTATATGAACGTGCATCAGTCCTAAG |
| 2011 | pIMAY-Down-sRNA260-R | CCTCACTAAAGGGAACAAAAGCTGGGTACCCGTTCGAGGATTCACTGTTCG |
| 2015 | pIMAY-Up-sRNA345-F | CGACTCACTATAGGGCGAATTGGAGCTCGTATTCTCTGAAGACGTTTGGAACA |
| 2016 | pIMAY-Up-sRNA345-R | GCGTATGGACCTAGGTATATCTGTCGTACACCTTGATATTAAAGGATTTC |
| 2017 | pIMAY-Down-sRNA345-F | ACCCCACAACCTAGGTATATCGTTTGTGTGGGGAATATGGAATA |
| 2018 | pIMAY-Down-sRNA345-R | CCTCACTAAAGGGAACAAAAGCTGGGTACCTGCCTTCAGTACATTATATAACCTTTGT |
| 2031 | pIMAY-Up-ncRNA2-F | CGACTCACTATAGGGCGAATTGGAGCTCGGCGTTCAATGGACTCTGTT |
| 2032 | pIMAY-Up-ncRNA2-R | GCGTATGGACCTAGGTATATCTTTTCATCTGTCCGATTTTTTGA |
| 2033 | pIMAY-Down-ncRNA2-F | ACCCCACAACCTAGGTATATCTTGTGCTTCTCAATGATACAATG |
| 2034 | pIMAY-Down-ncRNA2-R | CCTCACTAAAGGGAACAAAAGCTGGGTACCTCACCACCCAGTCATCAACA |
| 2038 | pIMAY-Up-ncRNA3-F | CGACTCACTATAGGGCGAATTGGAGCTCTCGGTTGCACATACAGCTTT |
| 2039 | pIMAY-Up-ncRNA3-R | GCGTATGGACCTAGGTATATAAAGTTTGAAGGTGATAATGTACATG |
| 2040 | pIMAY-Down-ncRNA3-F | ACCCCACAACCTAGGTATATAAACACTTTGCCCAACTTGC |
| 2041 | pIMAY-Down-ncRNA3-R | CCTCACTAAAGGGAACAAAAGCTGGGTACCTTTTACGGGTCTGTTTTCTAATTTGA |
| 2066 | Up-Sau5949-F | CGACTCACTATAGGGCGAATTGGAGCTCTCGTAATAATCGTGTGGCCA |
| 2067 | Up-Sau5949-R | GCGTATGGACCTAGGTATATACAATGTTATACTAAATACCTTTGA |
| 2068 | Down-Sau5949-F | ACCCCACAACCTAGGTATATGTACGAAAAAACACTTATGATTGTATGT |
| 2069 | Down-Sau5949-R | CCTCACTAAAGGGAACAAAAGCTGGGTACCACGGTAATTCAATCTATAGGTCTTGT |
| 2119 | pIMAY-Up-ssrS-F | CGACTCACTATAGGGCGAATTGGAGCTCACTCGTAAAGATATGGATGCTT |
| 2120 | pIMAY-Up-ssrS-R | GCGTATGGACCTAGGTATATTATCTTATGATGTTATATTACCACATAATT |
| 2121 | pIMAY-Down-ssrS-F | ACCCCACAACCTAGGTATATTCTATCGATACGCAAGACTTTGTC |
| 2122 | pIMAY-Down-ssrS-R | CCTCACTAAAGGGAACAAAAGCTGGGTACCGGTGGCATTTGTCCTTTTCG |
| 2126 | pIMAY-Up-sprF2-F | CGACTCACTATAGGGCGAATTGGAGCTCTGGATGGATTAAGAGGTCGTGT |

| | | |
|---|---|---|
| 2127 | pIMAY-Up-sprF2-R | GCGTATGGACCTAGGTATATCCACTATAATGAAGCATGCCTC |
| 2128 | pIMAY-Down-sprF2-F | ACCCCACAACCTAGGTATATTGTCGTCTTTTTACATTTTTATA GTAAC |
| 2129 | pIMAY-Down-sprF2-R | CCTCACTAAAGGGAACAAAAGCTGGGTACCACGCTCTATTGAC CCACCAA |
| 2140 | pIMAY-Up-sprA1As1-F | CGACTCACTATAGGGCGAATTGGAGCTCACTAACAAATAATAC ACCAGCAGCT |
| 2141 | pIMAY-Up-sprA1As1-R | GCGTATGGACCTAGGTATATCACAGTCACTTGCTTCTGATAAG TTA |
| 2142 | pIMAY-Down-sprA1-F | ACCCCACAACCTAGGTATATGTGAGGGGATTGGTGTATAAGT |
| 2143 | pIMAY-Down-sprA1-R | CCTCACTAAAGGGAACAAAAGCTGGGTACCCGATTTATATGAA GTACAATGTGAAAGG |
| 2147 | pIMAY-Up-sprF3-F | CGACTCACTATAGGGCGAATTGGAGCTCATGCAGATAGTACAC ACCTGATTG |
| 2148 | pIMAY-Up-sprF3-R | GCGTATGGACCTAGGTATATCCAACTTTCCATACAGCAGAAAA TAC |
| 2149 | pIMAY-Down-sprF3-F | ACCCCACAACCTAGGTATATCGATAAACAGTTGAGTGACATAC CC |
| 2150 | pIMAY-Down-sprF3-R | CCTCACTAAAGGGAACAAAAGCTGGGTACCAGAGAACGGATAT ACAATTGATAAAGAAGA |
| 2161 | pIMAY-Up-SprF1-F | CGACTCACTATAGGGCGAATTGGAGCTCGGCGCTTTACTTCCA ACTGT |
| 2162 | pIMAY-Up-SprF1-R | GCGTATGGACCTAGGTATATCACACCATAATATAAATATCAAA TAGACGG |
| 2163 | pIMAY-Down-SprF1-F | ACCCCACAACCTAGGTATATTAAAAAGTCAGTACCGAAGCACT |
| 2164 | pIMAY-Down-SprF1-R | CCTCACTAAAGGGAACAAAAGCTGGGTACCCAACAATGTGCTG AGGAAGAGT |
| 2168 | pIMAY-Up-SprY3-F | CGACTCACTATAGGGCGAATTGGAGCTCAGATTAGAAGCGGGC ATTGC |
| 2169 | pIMAY-Up-SprY3-R | GCGTATGGACCTAGGTATATGTAAATAGAAAGCAGGTATGTAA CGC |
| 2170 | pIMAY-Down-SprY3-F | ACCCCACAACCTAGGTATATAACAGGCAGGTACTACGGTA |
| 2171 | pIMAY-Down-SprY3-R | CCTCACTAAAGGGAACAAAAGCTGGGTACCGAATCTCTTCGGC AACTTTG |
| 2175 | pIMAY-Up-SprX1-F | CGACTCACTATAGGGCGAATTGGAGCTCAACAGTTGGAAGTAA AGCGC |
| 2176 | pIMAY-Up-SprX1-R | GCGTATGGACCTAGGTATATCTTGAATACGTCTAGAAAGATTA TAACAT |
| 2177 | pIMAY-Down-SprX1-F | ACCCCACAACCTAGGTATATTATGACTTTAGCATTCCCGTATA ATAGT |
| 2178 | pIMAY-Down-SprX1-R | CCTCACTAAAGGGAACAAAAGCTGGGTACCACTCATTTTAGGA ATTTCGCAAA |
| 2182 | pIMAY-Up-SprY1-F | CGACTCACTATAGGGCGAATTGGAGCTCAACACACCATCGTTT GTTCC |
| 2183 | pIMAY-Up-SprY1-R | GCGTATGGACCTAGGTATATACATATTCAATCAAGACATTGCT T |

| | | |
|---|---|---|
| 2184 | pIMAY-Down-SprY1-F | ACCCCACAACCTAGGTATATATCAGTTAGGATGAAAAAGTGGAT |
| 2185 | pIMAY-Down-SprY1-R | CCTCACTAAAGGGAACAAAAGCTGGGTACCTACACACCATCATTCAGCGA |
| 2189 | pIMAY-Up-SprX2-F | CGACTCACTATAGGGCGAATTGGAGCTCAACGGAACAAATGAACGTGA |
| 2190 | pIMAY-Up-SprX2-R | GCGTATGGACCTAGGTATATATCATAACAAAAAACTAGCCCGAAG |
| 2191 | pIMAY-Down-SprX2-F | ACCCCACAACCTAGGTATATTTAGCATTCCCGTATAACAGTTTAC |
| 2192 | pIMAY-Down-SprX2-R | CCTCACTAAAGGGAACAAAAGCTGGGTACCCATGCCCTATTTTATTTGTTGATGA |
| 2196 | pIMAY-Up-SprY2-F | CGACTCACTATAGGGCGAATTGGAGCTCAGCGTTATTAAGCAAGCAACT |
| 2197 | pIMAY-Up-SprY2-R | GCGTATGGACCTAGGTATATGTAAGATTCCCTATAATTAATGTAGCAAAA |
| 2198 | pIMAY-Down-SprY2-F | ACCCCACAACCTAGGTATATTATGTTATAGCTAGCCTTCGGG |
| 2199 | pIMAY-Down-SprY2-R | CCTCACTAAAGGGAACAAAAGCTGGGTACCATGCAACGACTGATAAACCG |
| 2201 | pIMAY-Up-RsaH-F | CGACTCACTATAGGGCGAATTGGAGCTCTTAAACGGACCACTAGCTGA |
| 2202 | pIMAY-Up-RsaH-R | GCGTATGGACCTAGGTATATGGTACACCTTTATTATAACTTATATCATTT |
| 2203 | pIMAY-Down-RsaH-F | ACCCCACAACCTAGGTATATTAGTGGACCCGTACGTTAATC |
| 2204 | pIMAY-Down-RsaH-R | CCTCACTAAAGGGAACAAAAGCTGGGTACCTTGCTTTGTAGGTGCTTGTT |
| 2208 | pIMAY-Up-hfq-F | CGACTCACTATAGGGCGAATTGGAGCTCGGTGAAATCATAAGCGGTGAC |
| 2209 | pIMAY-Up-hfq-R | GCGTATGGACCTAGGTATATCTGTCGGACTCCTTTTACTTAATC |
| 2210 | pIMAY-Down-hfq-F | ACCCCACAACCTAGGTATATACGCTTCATATAAAGGTCGAGT |
| 2211 | pIMAY-Down-hfq-R | CCTCACTAAAGGGAACAAAAGCTGGGTACCCAACATAATATTTGCGATCTACACG |
| 2215 | pIMAY-Up-sau5971-F | CGACTCACTATAGGGCGAATTGGAGCTCAATTACTTCTTCAAACTAGCTTATTTCCG |
| 2216 | pIMAY-Up-sau5971-R | GCGTATGGACCTAGGTATATACTAGATAGTTTATACTTTTGGTCTGTTG |
| 2217 | pIMAY-Down-sau5971-F | ACCCCACAACCTAGGTATATGAAGAGCTATGCATTTTATTTAAAAT |
| 2218 | pIMAY-Down-sau5971-R | CCTCACTAAAGGGAACAAAAGCTGGGTACCCGAAGCAGATTTTATTAACTGTGT |
| 2222 | pIMAY-Up-00009-F | CGACTCACTATAGGGCGAATTGGAGCTCTGTGAACGCAGATACAATGT |
| 2223 | pIMAY-Up-00009-R | GCGTATGGACCTAGGTATATACCCCATTAAACCACAAACT |
| 2224 | pIMAY-Down-00010-F | ACCCCACAACCTAGGTATATGAACGCATTTCATTATAGCAACAA |

| | | |
|---|---|---|
| 2225 | pIMAY-Down-00010-R | CCTCACTAAAGGGAACAAAAGCTGGGTACCTGCATCCAACGATCATTGAT |
| 2228 | pIMAY-Up-03030-F | CGACTCACTATAGGGCGAATTGGAGCTCCTTGTTTCCTTAATTGTTGTACCT |
| 2229 | pIMAY-Up-03030-R | GCGTATGGACCTAGGTATATAGCACGCAATGATTTAAAGGAT |
| 2230 | pIMAY-Down-03031-F | ACCCCACAACCTAGGTATATTCACTGAAAATTTGTATAAAGATTTAAGTC |
| 2231 | pIMAY-Down-03031-R | CCTCACTAAAGGGAACAAAAGCTGGGTACCGTCACGTCCTACAAACAAGT |
| 2234 | pIMAY-Up-01263-F | CGACTCACTATAGGGCGAATTGGAGCTCGCTACATTTGAAGTGAACGC |
| 2235 | pIMAY-Up-01263-R | GCGTATGGACCTAGGTATATGGATAGAAAACCAATCATCTTTATAGG |
| 2236 | pIMAY-Down-01264-F | ACCCCACAACCTAGGTATATAATAAAAAAGAAGAGAAGATGTAACACA |
| 2237 | pIMAY-Down-01264-R | CCTCACTAAAGGGAACAAAAGCTGGGTACCGAGTTTGTTCTTGTGCTTCC |
| 2343 | *isrR* amplification-F | CTGCAGGTCGACTCTAGAGGTGTGCGATTTTGAACTTGGA |
| 2344 | *isrR* amplification-R | CCATTCAGGCTGCGCAACTGGCGGTCATGCTATGGGATCA |
| 2363 | pIMAY-Up-S808-F | CGACTCACTATAGGGCGAATTGGAGCTCTGCCCCACCTAATCAGATAT |
| 2364 | pIMAY-Up-S808-R | GCGTATGGACCTAGGTATATACCGAGTCATTTCAAGAATG |
| 2365 | pIMAY-Down-S808-F | ACCCCACAACCTAGGTATATGATAACCGCATCTTAACTGA |
| 2366 | pIMAY-Down-S808-R | CCTCACTAAAGGGAACAAAAGCTGGGTACCTCGTTTGCTAGAATAATTGCT |
| 2379 | pIMAY-Up-S596-F | CGACTCACTATAGGGCGAATTGGAGCTCTTGATGAAGAACAATTAACAGCA |
| 2380 | pIMAY-Up-S596-R | GCGTATGGACCTAGGTATATCGTTTTATAAAAGCAGTAAACCCT |
| 2381 | pIMAY-Down-S596-F | ACCCCACAACCTAGGTATATGATGTTCTATGTGGTATTGATAATCA |
| 2382 | pIMAY-Down-S596-R | CCTCACTAAAGGGAACAAAAGCTGGGTACCTGCGACAAATTTCTAAGCCA |
| 2387 | pIMAY-Up-S204-F | CGACTCACTATAGGGCGAATTGGAGCTCGCTTTAACTGCCATCGTTAC |
| 2388 | pIMAY-Up-S204-R | GCGTATGGACCTAGGTATATATAACCACATCACATAAATTGAGTTC |
| 2389 | pIMAY-Down-S204-F | ACCCCACAACCTAGGTATATGTCTTAGTAAATCATACGTTCTATGT |
| 2390 | pIMAY-Down-S204-R | CCTCACTAAAGGGAACAAAAGCTGGGTACCTATTGAATGCCGACAGACTC |
| 2395 | pIMAY-Up-RsaC-F | CGACTCACTATAGGGCGAATTGGAGCTCCCTGTTGGTCAAGATCCTCA |
| 2396 | pIMAY-Up-RsaC-R | GCGTATGGACCTAGGTATATTGTTGATGTGTGGCCTAAAA |

| | | |
|---|---|---|
| 2397 | pIMAY-Down-RsaC-F | ACCCCACAACCTAGGTATATCCTAAAATAAAAGGGATTGATGAAAAGC |
| 2398 | pIMAY-Down-RsaC-R | CCTCACTAAAGGGAACAAAAGCTGGGTACCCGAAACCTGCACCTAAAACA |
| 2499 | +1 *isrR* for pRMC2 F | GATAGAGTATAATTAAAATAAGCGTTGAAAATGATTATCAATACCAC |
| 2500 | +1 *isrR* for pRMC2 R | TCAGATCTGTTAACGGTACCAAATAGTAAAAAAACAAAAGCAGTAAAC |
| 2507 | pIMAY-locus2 R | CGCACATTGAAATGATGTGTGAGCACGCAATGATTTAAAGGAT |
| 2508 | pIMAY-locus2 F | TTGATCCCATAGCATGACCGTCACTGAAAATTTGTATAAAGATTTAAGTC |
| 2509 | pIMAY-locus3 R | CGCACATTGAAATGATGTGTGGGATAGAAAACCAATCATCTTTATAGG |
| 2510 | pIMAY-locus3 F | TTGATCCCATAGCATGACCGAATAAAAAAGAAGAGAAGATGTAACACA |
| 2511 | *isrR* for pIM-locus2 or 3 F | CACACATCATTTCAATGTGCG |
| 2512 | *isrR* for pIM-locus2 or 3 R | CGGTCATGCTATGGGATCAA |
| 2534 | Linearization pCN34 R | CTCGGTACCCGGGGATCCTC |
| 2535 | Linearization pCN34 F | CCGTCGTTTTACAACGTCGTG |
| 2538 | Δ*tetR* from pRMC2-isrR R | CCACAGACAAATCACAGATACTAGTTTTTTATTTGGATCCCC |
| 2539 | Δ*tetR* from pRMC2-isrR F | TATCTGTGATTTGTCTGTGGAAGCAGCATAACCTTTTTCCG |
| 2546 | pCN38-isrR (*isrR* promoter-terminator) F | ATGGTCATAGCTGTTTCCTGATAAAAGCAGTAAACCCTTACGA |
| 2547 | pCN38-isrR (*isrR* promoter-terminator) R | ATAATCATCCTCCTAAGGTACCCGGATGTTCTATGTGGTATTGATAATC |
| 2552 | PCR of GFP with an RBS (from pCM11) F | GGGTACCTTAGGAGGATGATTA |
| 2553 | PCR of GFP with an RBS (from pCM11) R | CAGGAAACAGCTATGACCATG |
| 2554 | PCR of *rrnB* terminator from pECTO F | CTGTTTCCTGAGTAGGGAACTGCCAGGC |
| 2555 | PCR of *rrnB* terminator from pECTO R | ACGACGTTGTAAAACGACGGAATTAATGACAATCCTACTCAGGAGAG |
| 2573 | Fur motif 1 mutagenesis F | TAGTTTCTAATTGACAATGAAAGACACTAATGTATAATAGTAGTTGAAAA |
| 2574 | Fur motif 1 mutagenesis R | TTTTCAACTACTATTATACATTAGTGTCTTTCATTGTCAATTAGAAACTA |
| 2575 | Fur motif 2 mutagenesis F | CTAATGTATAATAGTAGTTGTTTTTGATTATCAATACCACATAGAACATC |
| 2576 | Fur motif 2 mutagenesis R | GATGTTCTATGTGGTATTGATAATCAAAAACAACTACTATTATACATTAG |

| | | |
|---|---|---|
| 2601 | pCN34-sGFP F | AGCAAAGGAGAAGAACTTTTCAC |
| 2600 | pCN34-sGFP R | TTAGTTAATTATAACTAATTAAAAATGAGAAGTAAAC |
| 2605 | *isrR*ΔCRR1 F | CCACATAGAACATACAACGTTTCGTTCTTGTTGGAT |
| 2606 | *isrR*ΔCRR1 R | ACGAAACGTTGTATGTTCTATGTGGTATTGATAATC |
| 2617 | narG 5'UTR-F for pCN34-GFP F | TTACTTCTCATTTTTAATTAGTTATAATTAACTAAAAAGCAATAGTCTTGGGCATTTT |
| 2618 | narG 5'UTR-R for pCN34-GFP R | GGGACAACTCCAGTGAAAAGTTCTTCTCCTTTGCTTCTACTTTTACTTTCTAGGATCG |
| 2619 | nasD 5'UTR-F for pCN34-GFP F | TTACTTCTCATTTTTAATTAGTTATAATTAACTAAACCTTTTTTTGAAATAAATATTATG |
| 2620 | nasD 5'UTR-R for pCN34-GFP R | GGGACAACTCCAGTGAAAAGTTCTTCTCCTTTGCTGCGCTCTAATATTTCTTCGATT |
| 2621 | gltB2 5'UTR-F for pCN34-GFP F | TTACTTCTCATTTTTAATTAGTTATAATTAACTAATACATTAAAATTTAAAATGAAAAA |
| 2622 | gltB2 5'UTR-R for pCN34-GFP R | GGGACAACTCCAGTGAAAAGTTCTTCTCCTTTGCTTATAAATTGCATGACTGTAAGAA |
| 2631 | *isrR*ΔCRR2 F | CATTTTCAAATATTCTTTTATATGCCCGTAAAAGACAA |
| 2632 | *isrR*ΔCRR2 R | GGCATATAAAAGAATATTTGAAAATGACCAATCCAAC |
| 2667 | *isrR*ΔCRR3 F | CCCTTTTATATGGTAAAAGACAATATACGTTATAACAACG |
| 2668 | *isrR*ΔCRR3 R | ATTGTCTTTTACCATATAAAAGGGGAATATTTGAAAATGA |
| 2694 | fdhA 5'UTR-F for pCN34-GFP F | TTAATTAGTTATAATTAACTAAAATTCTATCTGAAAGATGTGTG |
| 2695 | fdhA 5'UTR-R for pCN34-GFP R | AAAAGTTCTTCTCCTTTGCTATCAAGTGTAACCACCAAATG |
| **Northern blot probes** | | |
| 2452 | IsrR | TCTTTTACGGGCATATAAAAGGGG |
| 2614 | *S. lugdunensis* IsrR | TCTTTTAAGGGCATATAAAAGGGG |
| 2526 | *fdhA* mRNA | TGGATACGCACCAGTACCAG |
| 2627 | *ssrA* (tmRNA) | CTTCAAACGGCAGTGTTTAGC |
| 2696 | *gltB2* mRNA F | ACGGTTATTGTTATCGGGCT |
| 2697 | *gltB2* mRNA R | AAGCGCCATAACTCATACCA |
| 2922 | IsrR EMSA | CAATCCAACAAGAACGAAACGTTG |
| **5'/3'RACE mapping** | | |
| 2728 | IsrR RT | CGTATATTGTCTTTTACGGGC |
| 2729 | IsrR 5'3' junction PCR F | CAACGTTTTATAAAAGCAGTAAACCC |
| 2730 | IsrR 5'3' junction PCR R | CAATCCAACAAGAACGAAACGTT |
| 2731 | IsrR 5'3' junction nested PCR F | CGACACTTTAGGTTTACTGCTTTTG |
| 2732 | IsrR 5'3' junction nested PCR R | GGGATGTTCTATGTGGTATTGATAATC |
| **SHAPE and EMSA. PCR for DNA template** | | |
| 2770 | IsrR F | TAATACGACTCACTATAGTTGAAAATGATTATCAATACCACATAG |

| | | |
|---|---|---|
| 2771 | IsrR R | ACAAAAGCAGTAAACCTAAAGTG |
| 2773 | FdhA F | TAATACGACTCACTATAGAATTCTATCTGAAAGATGTGTGG |
| 3018 | FdhA R | GGTACAAAAGTATCTTGTGATT |
| 3015 | FdhA 7mut_R | CAAGTGTAACCACCAAATGTTCTTG |
| 2775 | GltB2 F | TAATACGACTCACTATAGACTTTAGCACATATTACTTTGTATTG |
| 2776 | GltB2 R | CGATAACAATAACCGTAAGCATG |
| 2777 | NarG F | TAATACGACTCACTATAGAAAGCAATAGTCTTGGGCATTTTAA |
| 2778 | NarG R | TTGTTCTTACTTCTTTATCGTGGCT |
| 2779 | NasD F | TAATACGACTCACTATAGATTAGAAAGCTTAATGATTCCAATG |
| 2780 | NasD R | ATAGTTTGGATAAGGTTCTTTACCT |
| **SHAPE. Reverse transcription** | | |
| - | IsrR | CCTAAAGTGTCGTAAGGG |
| - | FdhA | TAATAAATTCAAGTAAATTCGTACC |
| - | GltB2 | AATCCTACAACGATAATGTTAAC |
| - | NarG | TTGTTCTTACTTCTTTATCGTGGCT |
| - | NasD | ATAGTTTGGATAAGGTTCTTTACCT |
| - | IsrR competitor | ACAAAAGCAGTAAA |

* F, forward primer; R, reverse primer.

**Table S4. Fitness library composition**

| Allele | Library 1 | Library 2 | Library 3 |
|---|---|---|---|
| ***ΔrnaIII*::tag004** | SAPhB618 | SAPhB618 | SAPhB618 |
| ***ΔrsaOG*::tag009** | SAPhB347 | SAPhB347 | SAPhB347 |
| ***ΔrsaG*::tag011** | SAPhB349 | SAPhB349 | SAPhB349 |
| ***Δteg147*::tag018** | SAPhB368 | SAPhB368 | SAPhB368 |
| ***ΔrsaB*::tag025** | SAPhB380 | SAPhB380 | SAPhB380 |
| ***ΔrsaD*::tag026** | SAPhB682 | SAPhB682 | SAPhB682 |
| ***Δteg116*::tag030** | SAPhB386 | SAPhB386 | SAPhB386 |
| ***Δsau85*::tag038** | SAPhB397 | SAPhB397 | SAPhB397 |
| ***Δsau6353*::tag042** | SAPhB402 | SAPhB402 | SAPhB402 |
| ***ΔrsaE*::tag045** | SAPhB404 | SAPhB404 | SAPhB404 |
| ***Δssr42*::tag050** | SAPhB412 | SAPhB412 | SAPhB412 |
| ***Δteg155*::tag053** | SAPhB415 | SAPhB415 | SAPhB415 |
| ***sprF3*::tag070** | SAPhB960 | SAPhB961 | SAPhB962 |
| ***sRNA334*::tag073** | SAPhB862 | SAPhB863 | SAPhB864 |
| ***rsaA*::tag075** | SAPhB943 | SAPhB944 | SAPhB945 |
| ***sau76*::tag076** | SAPhB890 | SAPhB891 | SAPhB962 |
| ***rsaOI*::tag077** | SAPhB883 | SAPhB884 | SAPhB885 |
| ***teg16*::tag080** | SAPhB865 | SAPhB866 | SAPhB867 |
| ***sRNA287*::tag085** | SAPhB871 | SAPhB872 | SAPhB873 |
| ***sRNA71*::tag086** | SAPhB874 | SAPhB875 | SAPhB876 |
| ***sRNA209*::tag093** | SAPhB907 | SAPhB908 | SAPhB909 |
| ***teg106*::tag095** | SAPhB899 | SAPhB900 | SAPhB901 |
| ***sRNA260*::tag096** | SAPhB921 | SAPhB922 | SAPhB947 |
| ***sRNA345*::tag097** | SAPhB910 | SAPhB911 | SAPhB912 |
| ***ncRNA2*::tag099** | SAPhB932 | SAPhB933 | SAPhB934 |
| ***ncRNA3*::tag100** | SAPhB940 | SAPhB941 | SAPhB942 |
| ***ssrS*::tag107** | SAPhB954 | SAPhB955 | SAPhB956 |
| ***sprF1*::tag110** | SAPhB1006 | SAPhB1007 | SAPhB1008 |
| ***sprX2*::tag111** | SAPhB974 | SAPhB975 | SAPhB976 |
| ***sprY2*::tag112** | SAPhB978 | SAPhB979 | SAPhB980 |
| ***sprY3*::tag113** | SAPhB957 | SAPhB958 | SAPhB959 |
| ***sau41*::Tag115** | SAPhB901 | SAPhB902 | SAPhB903 |
| ***sau5949*::tag117** | SAPhB948 | SAPhB949 | SAPhB950 |
| ***sprF2*::tag118** | SAPhB966 | SAPhB967 | SAPhB998 |
| ***sprB*::tag121** | SAPhB1031 | SAPhB1032 | SAPhB1033 |
| ***rsaC*::tag133** | SAPhB1242 | SAPhB1243 | SAPhB1244 |
| ***S204*::tag134** | SAPhB1234 | SAPhB1235 | SAPhB1236 |
| ***S596*::tag135** | SAPhB1231 | SAPhB1232 | SAPhB1233 |
| ***S808*::tag137** | SAPhB1239 | SAPhB1240 | SAPhB1241 |
| ***locus3*::tag139** | SAPhB1015 | SAPhB1016 | SAPhB1017 |
| ***locus2*::tag140** | SAPhB1012 | SAPhB1013 | SAPhB1014 |
| ***locus1*::tag141** | SAPhB1009 | SAPhB1010 | SAPhB1011 |
| ***sau5971*::tag142** | SAPhB1018 | SAPhB1019 | SAPhB1020 |
| ***sprA1*::tag144** | SAPhB976 | SAPhB977 | SAPhB996 |
| ***sprX2*::tag145/*sprX1*::tag149** | SAPhB1027 | SAPhB1028 | SAPhB1029 |
| **sprX1::tag146** | SAPhB1003 | SAPhB1004 | SAPhB1005 |
| ***rsaH*::tag147** | SAPhB971 | SAPhB972 | SAPhB973 |
| ***sprY1*::tag148** | SAPhB1021 | SAPhB1022 | SAPhB1023 |

### Table S5. IsrR sequences in *Staphylococcus* genus

---

>NC_007795 (*Staphylococcus aureus* NCTC 8325
AAAATAGCGTAATCATGCGTTTTATTTACTATTCTTAAAAAATATTCAAAAAAAGTTTTAGTTTC**TAATTGACAAT**
**G**ATTCTCACTAATGTATAATAGTA**GTTGAAAATGATTATCAAT**ACCACATAGAACATCCCCCCCACAACGTTTCGT
TCTTGTTGGATTGGTCATTTTCAAATATTCCCCTTTTATATGCCCGTAAAAGACAATATACGTTATAACAACGTTT
TATAAAAGCAGTAAACCCTTACGACACTTTAGGTTTACTGCTTT

---

>LS483491.1 (*Staphylococcus auricularis* NCTC12101)
ACTCTTCATTATAAATAATCTCAAATAAATGTATAGCAAAAAAAGTGAAAAGAAAGTTGCAGGATT**TAATTGACAA**
**TAATTCTC**AGTCATGTATAATGTAGTTGGAAATGATTATCAATACCAAGTAAGATCTTCCCCCCCACACTTATATG
TTCTTATTGGATTGATCATTTTCATCAATATCCCCTTTTATATTCCCGTAAAAGACTAACGTTGTAAATCAACGTG
TGAGATAAAGACCGTTTTTACGAATGTAAAACGGTCTTTATAATAAACA

---

>NZ_CP016760 (*Staphylococcus carnosus* LTH 3730)
TTAAAGATACTCATTTTTGTGTAAATAACTTACAGAAAAGAAAAATTCAAAAAAGTTACTTTTC**TAATTGACAATC**
**ATTCTCAAT**AATGTATAATAGTAC**TTGAAAATGATTATCAAT**ACCAAATGAGACATCCCCCCACAATTTCGTTCTT
ATAGTATTGGTCATTTTCATATCCCCTTTTATATATGCCCGTAAATACTGACGTTATGGTTATACATAACGTGACA
TACTTACCGATTTTGCTAGTAAATGCAAAATCGGTTTT

---

>NC_002976 (*Staphylococcus epidermidis* RP62A)
TTTAGAAAAAGTCACTCTAGCATTATCATTGTATTTAAGTTAAATGGAATAAAATATATATTTC**TAATTGACA**ATC
ATTATCAATCATGTATAATGATA**ATTGAAAATGATTATCAAT**ACCAATTGAAAAACATTCCCCCCACATACAAGTT
GTTCTTTTGGATTGGTCATTTTCAACTATCCCCTTTTATATGCCCGTAAAAGACTAACGTTAAGAAATGACGTTTC
AATAAAAGCAGTAGACCTTTGACACTTGAGGTCTGCTGTTTT

---

>NC_007168 (*Staphylococcus haemolyticus* strain Sh29/312/L2)
AAATAGAGAATGGATTAAAATTGTAATGATATAATCAAATTTAAATGAGAATTTTTCGCAATTA**TAATTGACA**ATG
ATTATCAATGATGTATAATAGTAT**TTGAAAATGATTATCAAT**ACCGAATAAGACAATTCCCCCCACATATATTTCG
TTCTTAAAGGATTGGTCATTTTCAAGTTATTCCCCTTTTATATGCCCGTAAAAGACTAACGTTAAAGTTTCAAACT
TACTTTAAACGTTTTAATAAAAAGCAGTGAATCTAATGCCGAGGT

---

>NZ_CP008747 (*Staphylococcus hyicus* strain ATCC 11249)
ACCTCACATCTATCATATTTCAAAACGAGCCCAGATAAAAGTTTTTTAAGAAAAATACGAGATTT**TAATTGACAAT**
**CATTCTC**AATAAGGTATAATGTAATTGAAAATGATTCTCAATCGTAACGACCCCCCACTACAATTCGTTCTTTTTG
ATTGAGGCATTTTCAAGTACTATCCCCTTTTATATGCCCGTATAAAAAAATAGTCGTTTTGATTAAACGTTAAATT
TAAGCTGCTACACTTATGTGTAGTAGCTCCTTTTTGT

---

>NZ_CP020768 (*Staphylococcus lugdunensis* strain C_33)
CACCAACCATAGTCTAAGATTTATGATTAACTTTCAAAAATTTATCATAAAAATTTCAGTTTTC**TAATTGACA**ATC
ATTATCAATGATGTATAATAATA**ATTGAAAATGATTATCAAT**ACCAAATAGAACTCCCCCCACATATTCGTTCTTA
TGGATTGATCATTTTCGAATTCCCCTTTTATATGCCCTTAAAAGACTAACGTAAAGCTTACAATAACGCTAAACGT
GTACATAAAAGCAGCTCCCTAATGGTAGCTGCTTT

---

>NC_007350 (*Staphylococcus saprophyticus* ATCC 15305)
TGATTTTAATCCCTTTTATTGTAATATATAGGTATTTCATAAAATTTGAAAAAATTATTCGAATT**TAATTGACAAT**
**CATTCTC**GGTGTTGTATAATGTAA**TTGAAAATGATTATCAAT**ACCAAAATAAGACATTCCCCCCACACATATTTCG
TTCTTATTTGGATTGATCATTTTCAAAAATATCCCCTTTTATATGCCCGTAAAAGACTAACGTTGCGAGACAACGT
GAATATAAAAACCGGTTTTACATTGTAAAATCGGTTTTTATAATAA

---

>NZ_CP022046 (*Staphylococcus sciuri* SNUDS-18)
AACATGTTCATTATATATGAGCATTACTATTTTATCATTTAAAATAGTAAAAATTTTAAATAAC**TAATTGACATTC**
**ATTCTC**AATTGGTTATAATAGTA**ATTGAAAATGATTATCAAT**CAAATAAAAGAGTTCCCCTCTAAGTATAGATTGA
ACATTTTCTATAACCCCCTTTTATGCCCATAAAAGTAAAATAGCCGCAGTGATTCTGTCCAAATCACTGTGGCTGT
TTTTTTTGTTTGGATTTATTGAGGA

---

>NZ_CP023497 (*Staphylococcus simulans* strain FDAARGOS_383)
ATCAAATGATTAATTGTGACAAAAATATGGAGTTGAATAAAAAAGTGAAATTATTTTTGATTTC**TAATTGACAACC**
**ATTCTC**ACTAATGTATAATAGTA**ATTGAAAATGATTATCAAT**CACAAATGAGACATCCCCCCACATTTCGTTCTTA
ATTGGATTGGTCATTTTCGATTTATCCCCTTTTATATATGCCCGTAAATACTGACGTTACACTAAAAGTAACGTGA
CATACTTAACCGATTCTCAATTAAGCAGAATCGGTTTTTTTCT

---

---

>NC_020164 (*Staphylococcus warneri* SG1)
TTCGTAATGTCAAAAATTGATGATGGTTAATAATTTATAAATAGTTCATCATTTTTCGTTATTTC**TAATTGACAAT**
**GATTCTC**ATTCATGTATAATAATAG**GTTGAAAATGATTATCAAT**ACCAAAAAGAACTTTCCCCCCCACATATATTTC
GTTCTTAAGGATTGGTCATTTTCATATAATCCCCTTTTATATGCCCGTAAAAGACTAACGTTGAAAAACGTTTTAA
TAAAGCAGTAGACCTTTAGACACTTGAGGTTTACTGCTTT

---

NCTC8325 *isrR* orthologs were searched within the Firmicutes phylum using GLASSgo (version 1.5.0 RNA tool) set with default parameters (11). As no sequence of *S. auricularis* was present in the GLASSgo dataset, the presence of an *isrR* sequence within this specie was search and found using the NCBI DNA sequence repository. An *isrR* ortholog (red characters) is present in all members of the staphylococcus genus. A sequence of one selected representative of each *Staphylococcus* genus group is shown. The 100 nt upstream sequences are shown (black characters). All sequences contains putative Fur boxes (characters in bold).

**Table S6. Proteins containing an Fe-S cluster in *S. aureus***

| Symbol | Fe-S cluster-containing protein |
|---|---|
| NirB | Nitrite reductase [NAD(P)H] large subunit |
| BioB | Biotin synthase |
| FdhL | Putative formate dehydrogenase |
| PflA | Pyruvate formate-lyase-activating enzyme |
| SdhB | Succinate dehydrogenase iron-sulfur subunit |
| MiaB-like | Uncharacterized protein |
| MiaB | tRNA-2-methylthio-N(6)-dimethylallyladenosine synthase |
| Nth | Endonuclease III |
| QueE | 7-carboxy-7-deazaguanine synthase |
| RlmN | Probable dual-specificity RNA methyltransferase |
| NirD | Nitrite reductase [NAD(P)H] small subunit |
| MoaA | GTP 3',8-cyclase |
| HemW | Heme chaperone |
| LipA | Lipoyl synthase |
| NarH | Respiratory nitrate reductase beta subunit |
| GltB | Glu_synthase domain-containing protein |
| QueG | Epoxyqueuosine reductase |
| GltB | Glutamate synthase large subunit |
| Ferredoxin | Ferredoxin |
| SufA | Fe-S_biosyn domain-containing protein |
| AddB | ATP-dependent helicase/deoxyribonuclease subunit B |
| SdaA | L-serine dehydratase |
| GltD | Glutamate synthase subunit beta |
| MutY | Adenine DNA glycosylase |
| AcnA | Aconitate hydratase |
| RumA | RNA methyltransferase |
| NrdG | Anaerobic ribonucleoside-triphosphate reductase-activating protein |
| YfkB | Uncharacterized protein |
| Grx | Glutaredoxin domain-containing protein |
| NarG | Nitrate reductase |
| Nfu | NifU domain-containing protein |
| LeuC | 3-isopropylmalate dehydratase large subunit |
| YhcC-like | Elp3 domain-containing protein |

List was manually curated for *S. aureus* strain USA300 combining data from MetalPredator (12) and UniProt (13) web servers, and whether the protein possesses the conserved cysteine ligands. (Béatrice Py, personal communication).

**Table S7. IsrR functional analogs and their targets**

| IsrR functional analog | Organism | RNA chaperone | mRNA targets and products | References |
|---|---|---|---|---|
| **FsrA** | *Bacillus subtilis* | FbpA, FbpB and FbpC | ***sdhCAB****, succinate dehydrogenase<br>***citB****, aconitase<br>***gltAB****, glutamate synthase<br>***leuCD***, isopropyl malate dehydratase<br>***ilvC***, ketol-acid reductoisomerase ***cydA***, cytochrome bd ubiquinol oxidase<br>***resA***, thiol-disulfide oxidoreductase<br>***ctaO***, heme O synthase<br>***cysH***, adenosine 5'-phosphosulfate reductase<br>***lutABC****, lactate oxidase<br>***dctP****, dicarboxylate permease | (14,15) |
| **MsrI** | *Mycobacterium tuberculosis* | - | ***bfrA****, bacterioferritin<br>***hypF***, hydrogenase maturation factor<br>***fprA***, NADPH-ferredoxin reductase<br>***acnA***, aconitase | (16) |
| **NrrF** | *Neisseria meningitidis* | Hfq | ***sdhCDAB****, succinate dehydrogenase<br>***petABC****, cytochrome $bc_1$<br>***gpxA***, glutathione peroxidase<br>***tadA***, tRNA-specific adenosine deaminase<br>***suhB***, extragenic suppressor protein<br>***mqo***, malate:quinone oxidoreductase<br>***hemO***, heme oxygenase | (17-19) |
| **PrrF1, PrrF2** | *Pseudomonas aeruginosa* | Hfq | ***sodB****, superoxide dismutase<br>***sdhCDAB****, succinate dehydrogenase<br>***acnA****, aconitase A<br>***acnB****, aconitase B<br>***katA***, catalase<br>***antABC****, anthranilate dioxygenase<br>***catBCA***, cathecol dissimilatory complex | (20,21) |
| **RyhB** | *Escherichia coli* | Hfq | Selected targets:<br>***acnA****, aconitase A<br>***acnB***, aconitase B<br>***bfr****, bacterioferritin<br>***fdhF***, formate dehydrogenase H<br>***fdoGHI***, formate dehydrogenase O<br>***gltB****, glutamate synthase large chain<br>***mqo***, malate:quinone oxidoreductase<br>***cydAB***, cytochrome d terminal oxidase<br>***hyaA***, hydrogenase-1 small chain<br>***katG***, catalase-peroxidase<br>***narG***, nitrate reductase alpha-chain<br>***narK***, nitrate/nitrite transporter<br>***nirB***, nitrite reductase large subunit<br>***sdhCDAB****, succinate dehydrogenase<br>***sodB****, superoxide dismutase<br>***mrsB****, methionine sulfoxide reductase<br>***oppB***, oligopeptide transport system permease protein | (22-30) |

*Validated targets. In red, shared functional targets with IsrR.

**Figure S1. Absence of IsrR is detrimental when iron is scarce**

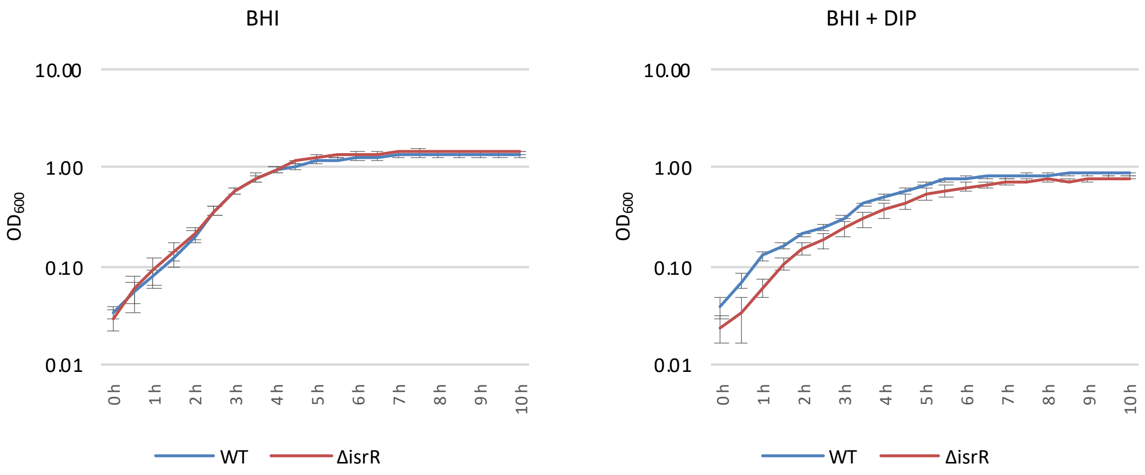

Growth of Δ*isrR* and its parental HG003 strain. Cultures were grown in rich medium (BHI) or BHI with 2,2'-dipyridyl (DIP) 1.5 mM at 37°C under vigorous agitation in microtiter plates in a volume of 200µL. Incubation and $OD_{600}$ were obtained using CLARIOstar microplate reader. Error bars indicate the standard deviation from three independent biological samples (n=3).

**Figure S2. *isrR* complementation restores optimal growth in low-iron conditions**

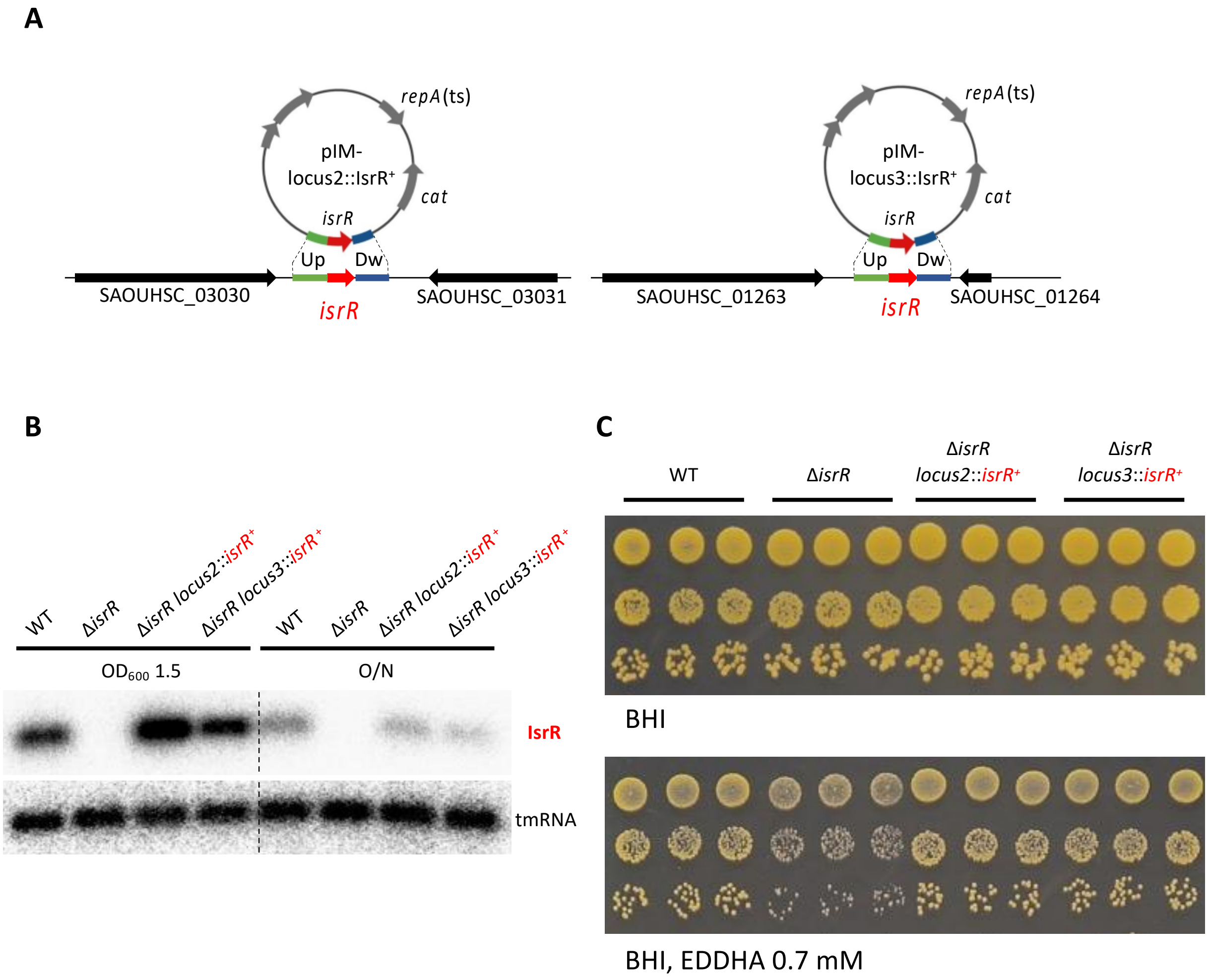

A) Schematic representation of constructions for chromosomal ectopic complementation of Δ*isrR*::*tag135* with insertion at locus2 (SAPhB1500) and locus3 (SAPhB1502). B) IsrR expression. Northern blot experiments with probes detecting IsrR and tmRNA (loading control). The indicated strains were sampled at $OD_{600}$ 1 and overnight cultures with DIP 1.25 mM (n=2). C) Plating efficiency (10-fold serial dilutions) of indicated strains on BHI medium without (upper panel) or with (lower panel) EDDHA 0.7 mM. Three independent biological clones are shown for each strain (n=3). For strains, see Table S1.

### Figure S3. IsrR 5'/3'RACE mapping and secondary structure prediction

**A**

GT$^{6}$TGA$^{6}$AAA$^{2}$TGA$^{2}$TTATCAATACCACATAGAACATCCCCCCCACAACGTTTCGTTCTTGTTGGATTGGT
CATTTTCAAATATTCCCCTTTTATATGCCCGTAAAAGACAATATACGTTATAACAACGTT TTATAAAAGCAGT
AAACCCTTACGACACTTTAGGTTTACTGCT T$^{1}$TT$^{1}$G$^{1}$T$^{6}$TT$^{2}$TT$^{1}$TTACTATTT

IsrR (174 nt):

TTGAAAATGATTATCAATACCACATAGAACATCCCCCCCACAACGTTTCGTTCTTGTTGGATTGGTCATTTTC
AAATATTCCCCTTTTATATGCCCGTAAAAGACAATATACGTTATAACAACGTTTTATAAAAGCAGTAAACCCT
TACGACACTTTAGGTTTACTGCTTTTGT

**B**

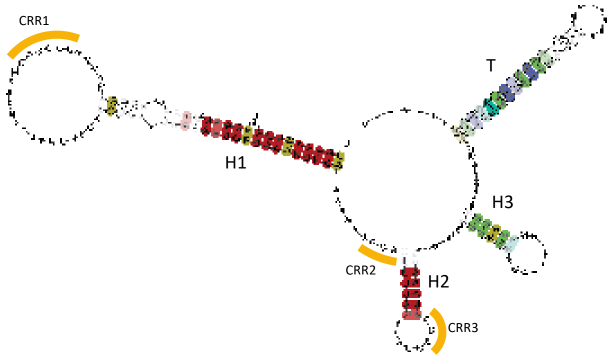

**C**

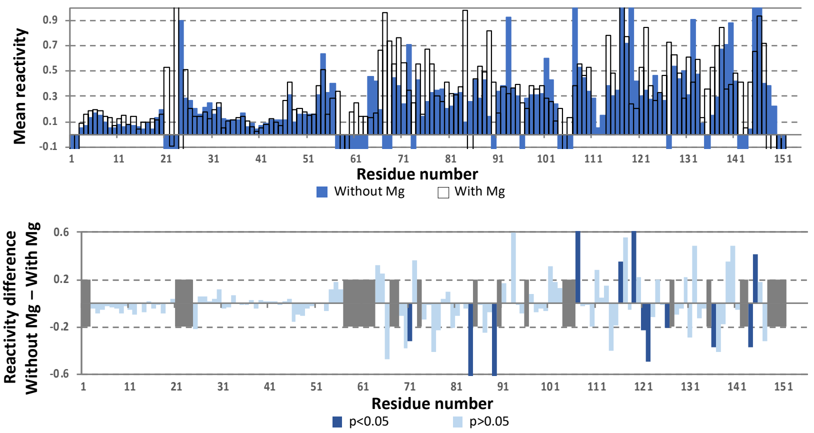

A) Mapping of IsrR extremities. Upper part, results from 5'/3'RACE mapping. Superscript numbers indicate the number of sequences ending with the corresponding nucleotide that were obtained. Lower part, IsrR sequence retained for the study. B) IsrR secondary structure predicted by LocARNA with 18 ortholog sequences for input (see Fig. 3A). LocARNA takes into account nucleotide covariations between different sequences restoring pairings to support the existence of stem-loop structures. Three stem loop structures (H1 to H3) and a rho-independent transcription terminator (T) are predicted. Three predicted C-rich regions (CRR1-3) are indicated. Colors indicate the number of base pairing types (red, 1; ochre, 2; green, 3, blue, 4; dark blue, 5; as per LocARNA parameters (31)) and hue shows sequence conservation (number of incompatible pairs: saturated, 0; medium, 1; light, 2). C) IsrR reactivity profiles to 1M7. Top panel: average reactivity to 1M7 for each IsrR nucleotide in the presence (white) or absence (blue) of MgCl2; tests were performed in triplicate. Bottom panel: difference of reactivity to 1M7 for each IsrR nucleotide when MgCl2 was added; nucleotides in intense blue presented a significant difference while those in light blue did not.

**Figure S4. IsrR putative targets**

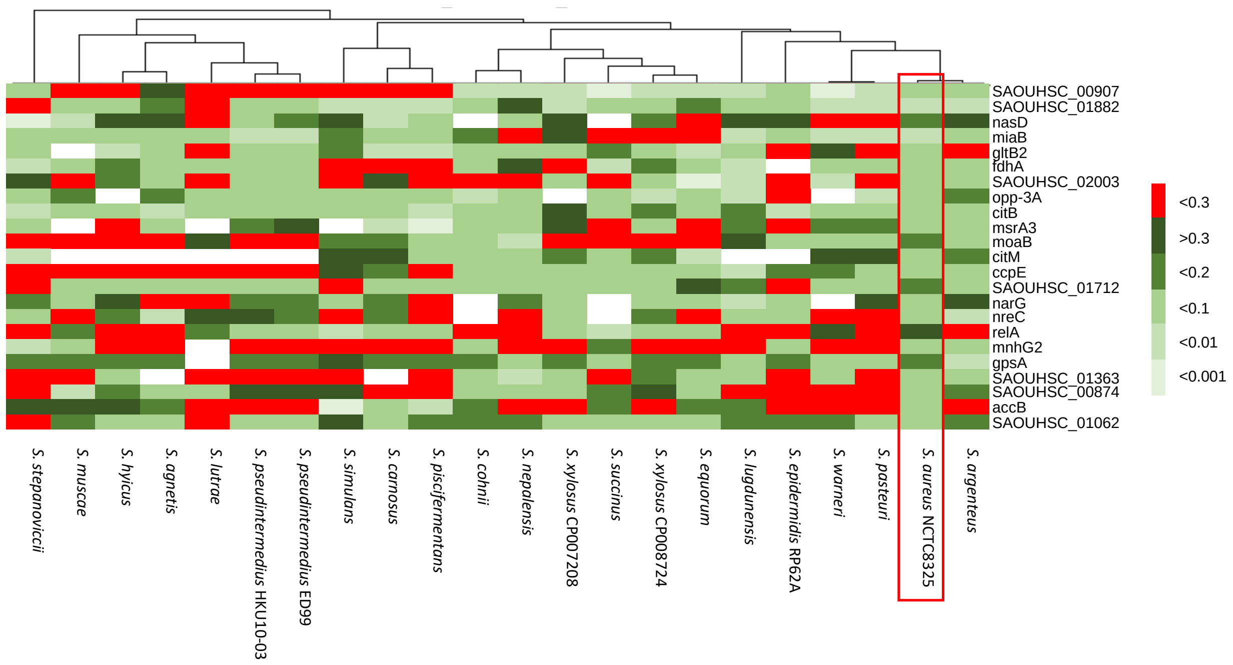

Adapted figure from a CopraRNA analysis with the 22 Staphylococci strains indicated using default parameters (32). Columns, investigated organisms; rows, targets; cell colors, IntaRNA p-value as indicated on the left panel; white cell, no homolog of a given target.

**Figure S5. IsrR putative targets involved in nitrate respiration pathway**

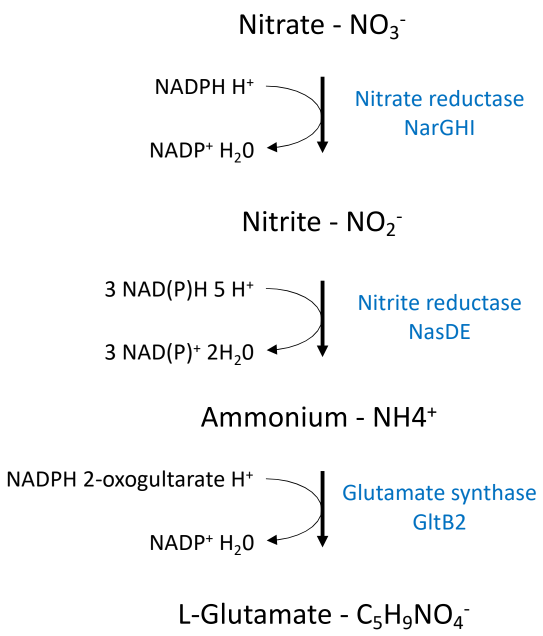

Dissimilatory nitrate reduction pathway. Adapted from BioCyc (33).

**Figure S6. Comparison of *fdhA* and *gltB2* mRNAs reactivity to 1M7 obtained in the presence/absence of IsrR**

**A**

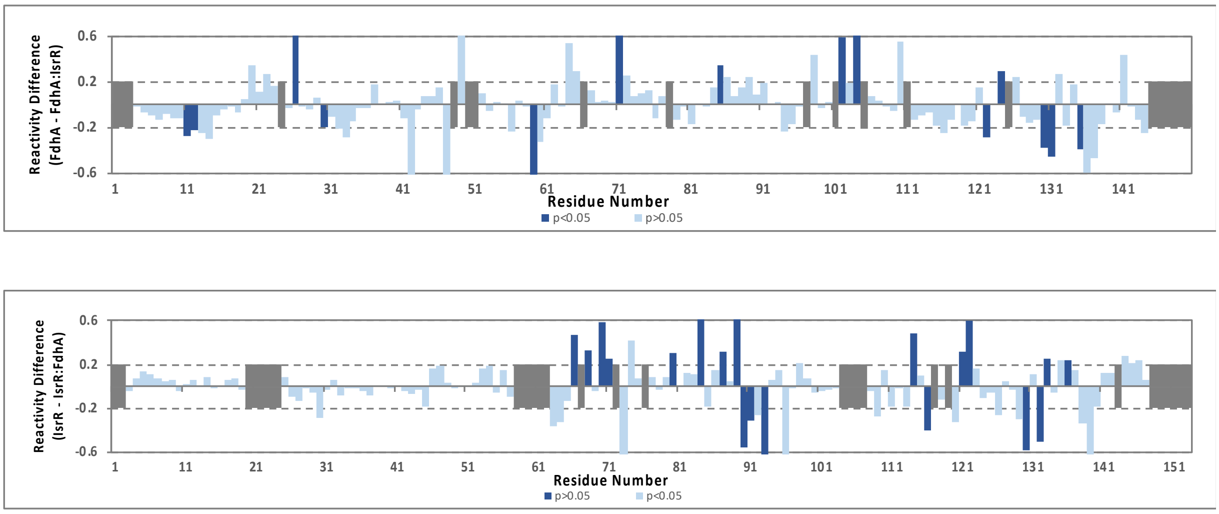

**B**

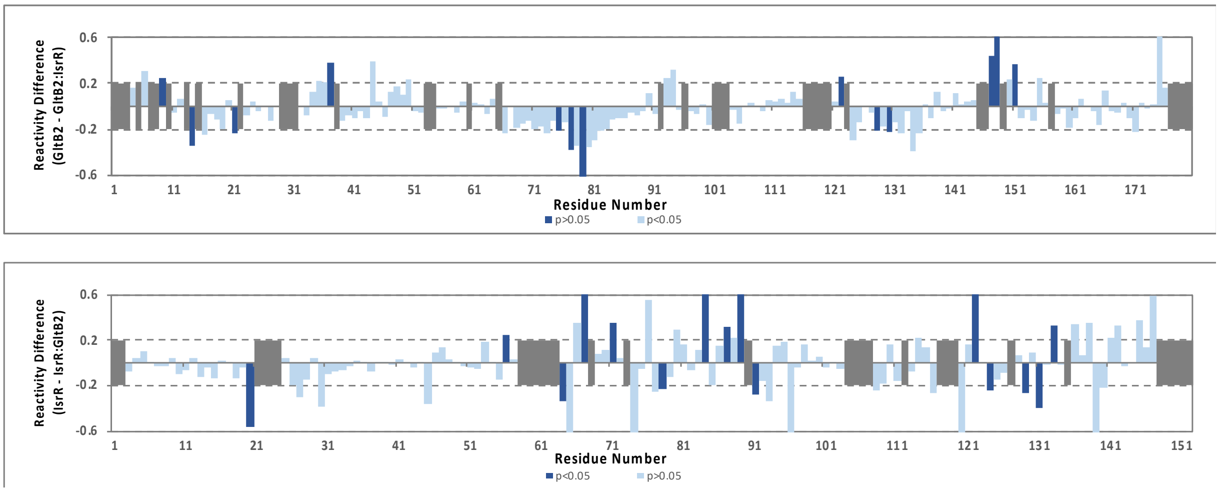

(A) Top panel: difference of reactivity to 1M7 for each nucleotide of *fdhA* mRNA when IsrR was added. Bottom panel: difference of reactivity to 1M7 for each nucleotide of IsrR when *fdhA* mRNA was added; nucleotides in intense blue presented a significant difference while those in light blue did not. (B) Top panel: difference of reactivity to 1M7 for each nucleotide of *gltB2* mRNA when IsrR was added. Bottom panel: difference of reactivity to 1M7 for each nucleotide of IsrR when *gltB2* mRNA was added; nucleotides in intense blue presented a significant difference while those in light blue did not.

### Figure S7. Comparison of *nasD* mRNA reactivity obtained to 1M7 in the presence/absence of IsrR with proposed interaction model

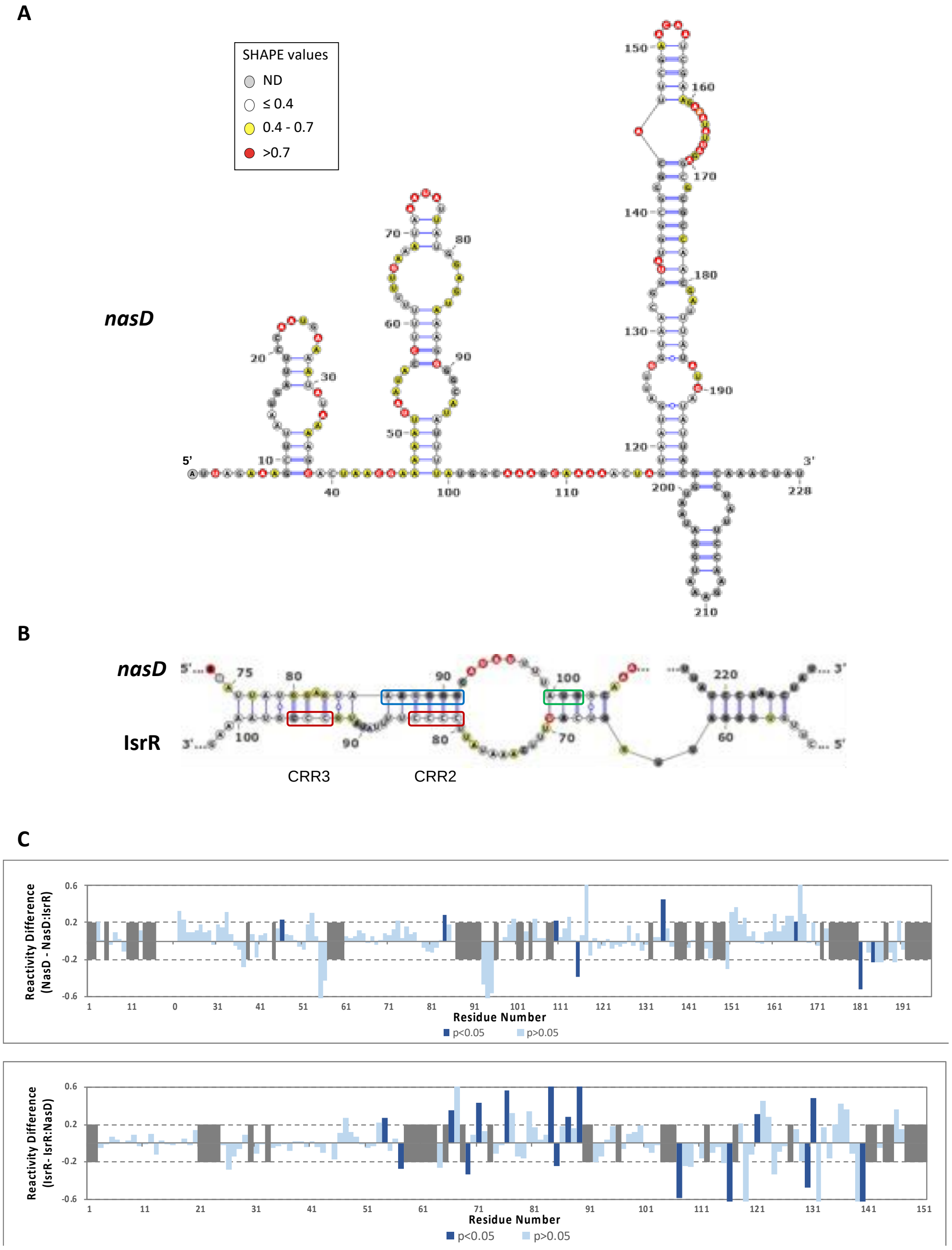

(A) Secondary structure model obtained with IPANEMAP for *nasD* 5'UTR using 1M7 reactivity as constraints. Nucleotides are coloured according to their reactivity in the absence of IsrR with the indicated colour code. ND, not determined. (B) Model for interaction between *nasD* mRNA and IsrR based on changes in reactivity in the presence of IsrR. mRNA Shine-Dalgarno sequence is shown in a blue rectangle and the start codon in a green rectangle; IsrR C-rich regions are shown in red rectangles. (C) Top panel: difference of reactivity to 1M7 for each nucleotide of *nasD* mRNA when IsrR was added. Bottom panel: difference of reactivity to 1M7 for each nucleotide of IsrR when *nasD* mRNA was added; nucleotides in intense blue presented a significant difference while those in light blue did not.

### Figure S8. Electrophoretic mobility shift assay of IsrR in the presence of *fdhA* mRNA

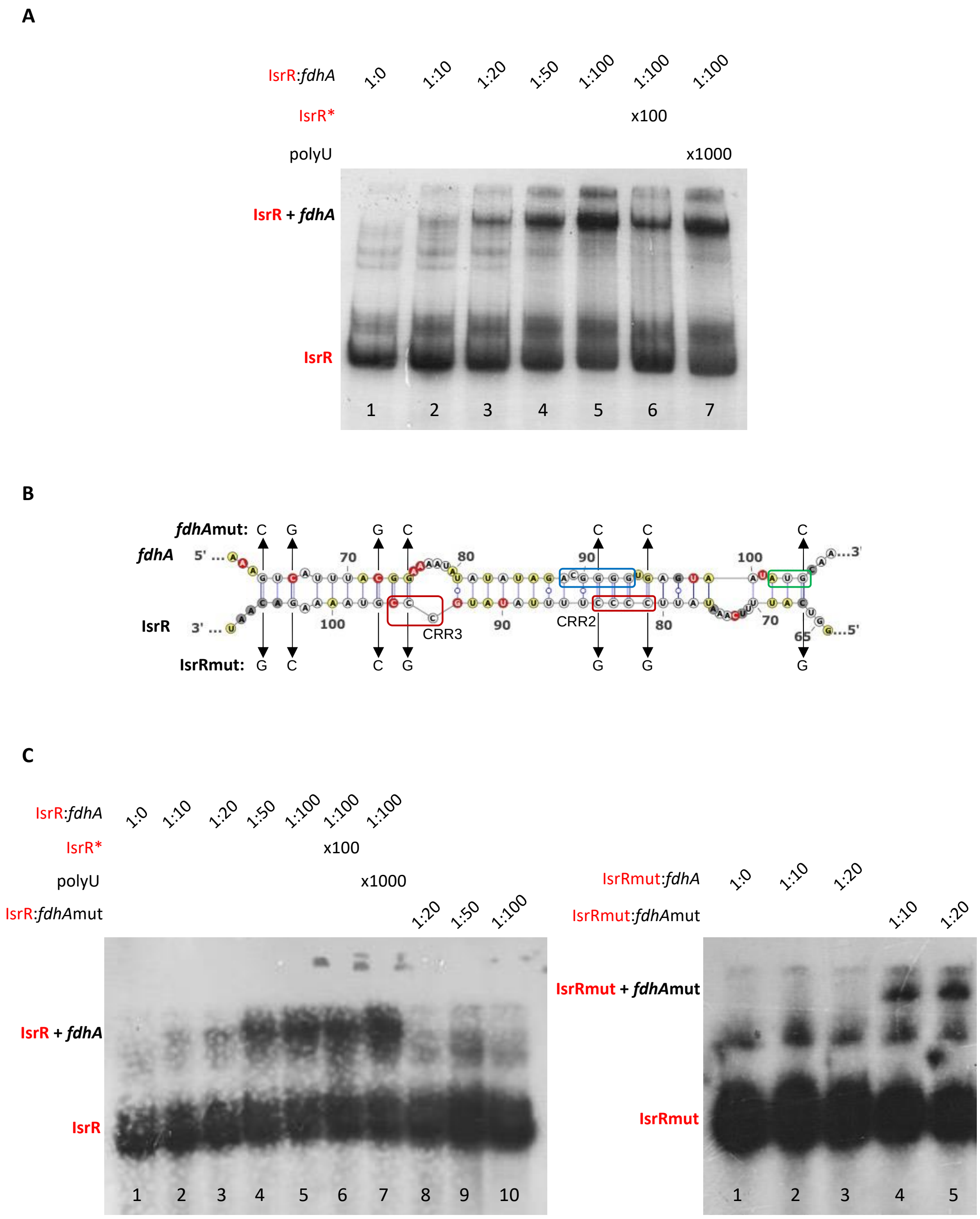

(A) Electrophoretic mobility shift assay (EMSA) with labelled IsrR. Constant amounts of IsrR (0.25 pmol) were mixed with increasing amounts of *fdhA* mRNA in the indicated proportions (columns 1-5). A specific competitor (unlabeled IsrR*, column 6) and a nonspecific competitor (polyU, column 7) were used as controls. (B) Predicted interaction between IsrR and *fdhA* mRNA as shown in Figure 6. Inserted point mutations (G↔C) are indicated with an arrow, resulting in IsrRmut and *fdhA*mut. (C) Left panel: EMSA with labelled IsrR as in (A) (columns 1-7) or mixed with increasing amounts of *fdhA* mRNAmut (columns 8-10). Right panel: EMSA with labelled IsrRmut. Constant amounts of IsrRmut were mixed with increasing amounts of *fdhA* mRNA (columns 1-3) or *fdhA* mRNAmut (columns 4-5) in the indicated proportions.

### Figure S9. Reporter fusions associated to nitrate respiration for IsrR activity tests

**A**

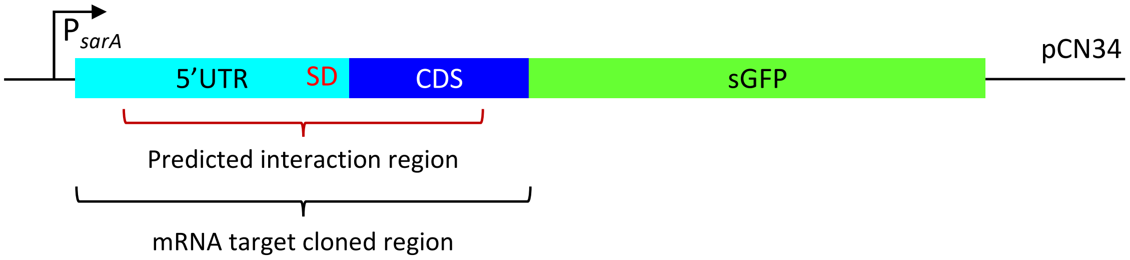

**p5'FdhA-GFP**

…aactaaAATTCTATCTGAAAGATGTGTGGGGCATCGTTATTTTAGGTGGATATGAGCAATTTATTAAAA **GTC**
**ATTTACGGAAAATATATATATAGACGGGGTGAGTAAT** **ATG CAA GAA CAT TTG GTG GTT** ACA CTT
GAT AGC AAA GGA GAA GAA CTT … CAC TAA GAATTC…

**p5'NarG-GFP**

…aactaaAAAGCAATAGTCTTGGGCATTTTAAAGTATTTAAAAACACGACATCTACACATCGTATTTTTATAC
GTA**GGAGGATATAAAT** **ATG GGA AAA TTT GGA TTG AAT TTC TTT AAG CCA ACA GAA AAA**
**TTT AAT GGG** AAT TGG TCG ATC CTA GAA AGT AAA AGT AGA AGC AAA GGA GAA GAA
CTT … CAC TAA GAATTC…

**p5'NasD-GFP**

…aactaaACCTTTTTTTGAAATAAATA**TTATGGAGTAAAGGGGCATATTTT** **ATG GCA AAG CAA AAA**
**CTA GTA ATG ATT GGT AAC GGT ATG GCG GG**C ATT CGA ACA ATC GAA GAA ATA TTA
GAG CGC AGC AAA GGA GAA GAA CTT … CAC TAA GAATTC…

**p5'GltB2-GFP**

…aactaaTACATTAAAATTTAAAATGAAAAACATATCATAAAATTGTTTTATAAAATGAAGCGCTTCCATTGT
GTTTTGT**TTTGTAAGGTGTATCATAAATATTGAATTGAAATTTTGGG** GGGAGGTATTGTA ATG ACG TTT
CTT ACA GTC ATG CAA TTT ATA AGC AAA GGA GAA GAA CTT … CAC TAA GAATTC…

**B**

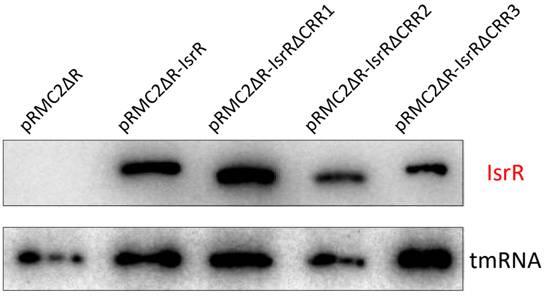

A) Upper part: Schematic representation of constructed reporter fusions. Below: *fdhA*, *narG*, *nasD* and *gltB2* cloned sequences. All sequences include the 5'UTR and first codons of IsrR target genes cloned in frame with the super-folder *gfp* CDS (minus its start codon). 5'UTR of IsrR target, light blue; first codons of IsrR target, dark blue; first and last sGFP codons, green; Shine-Dalgarno sequence, red; corresponding sequence to the predicted IsrR pairing region, bold italic font. B) *isrR* and its ΔCRR derivatives expression. Northern blot experiment. Total RNA extracts from HG003 Δ*isrR* derivatives containing the indicated plasmids. Membranes were probed for IsrR and tmRNA (loading control).

**Figure S10. Translational down-regulation of *narG* and *nasD* mRNAs by IsrR and CRR contribution**

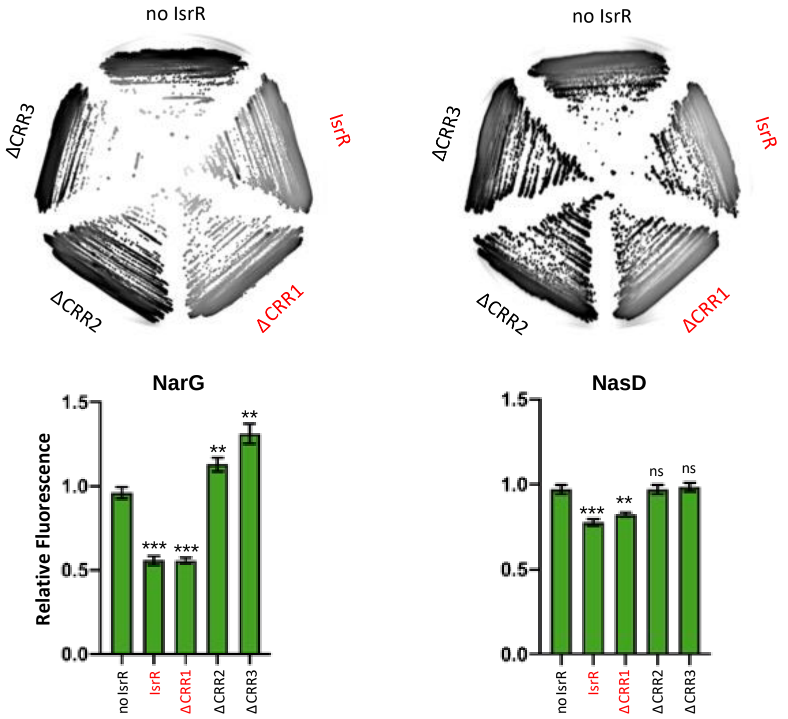

Leader fusions between the first codons of *narG* and *nasD* with a GFP were constructed (Figure S9 and Table S2). Δ*isrR* derivatives with either a control plasmid (no IsrR; pRMC2ΔR), or plasmids expressing IsrR (pRMC2ΔR-*isrR*), IsrRΔCRR1 (pRMC2ΔR-*isrR*ΔCRR1), IsrRΔCRR2 (pRMC2ΔR-*isrR*ΔCRR2), IsrRΔCRR3 (pRMC2ΔR-*isrR*ΔCRR3) were transformed with each engineered reporter gene fusions. Translational activity from the two reporters in the presence of the different *isrR* derivatives were evaluated by fluorescent scanning of streaked clones on plates (n=3). Fully active IsrR derivatives are shown in red. Translational activity of the reporter genes with the different *isrR* derivatives was also determined in liquid culture. The fluorescence of the 10 strains was measured in 6 h cultures using a microtiter plate reader. Results are normalized to 1 for each fusion with the control plasmid. Error bars indicate the standard deviation from three independent experiments (n=3). Statistical analyses were performed using t-test with Welch's correction: *** represents p-value between 0.0001-0.0004; ** represents p-value between 0.002-0.006; ns, non-significant.

**Figure S11. Hfq is not required for IsrR activity**

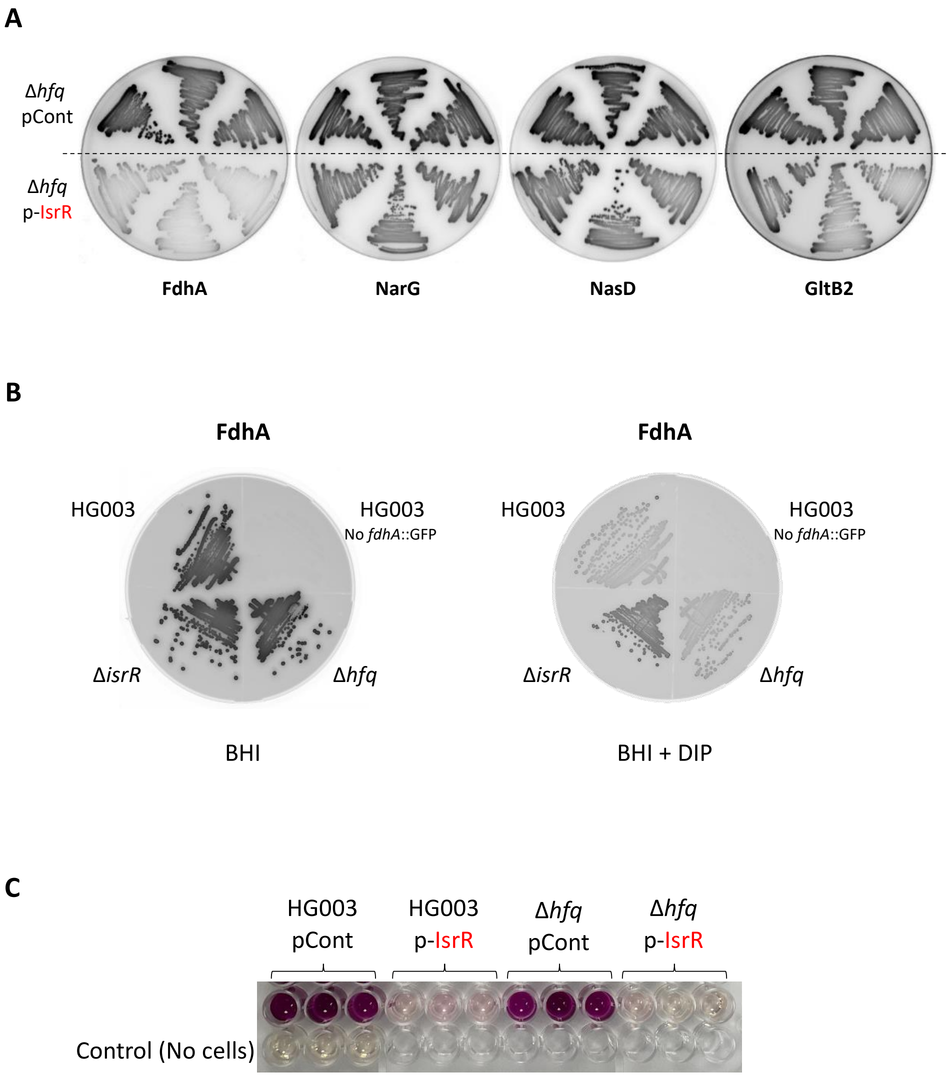

(A) Translational down-regulation of four targets by IsrR in Δ*hfq* background. Leader fusions between *fdhA*, *narG*, *nasD* and *gltB2* are the same as in Figure 7 and Supplementary Figure S10. HG003 Δ*hfq* derivatives with either a control plasmid (pCont; pRMC2ΔR), or a plasmid expressing IsrR (p-IsrR; pRMC2ΔR-*isrR*) were transformed with either p5'FdhA-GFP, p5'NarG-GFP, p5'NasD-GFP or p5'GltB2-GFP. Translational activity from the four reporters in the presence of *isrR* or not, was evaluated by fluorescent scanning of streaked clones on plates (n=3). (B) Translational down-regulation of *fdhA*::GFP leader fusion by IsrR. HG003 WT strain and its Δ*isrR* and Δ*hfq* derivatives were transformed with p5'FdhA-GFP and grown either in rich media (BHI), or BHI supplemented with DIP 0.5 mM. Expression of IsrR after iron chelation decreased translation of *fdhA* leader fusion in the Δ*hfq* strain as observed with the WT strain. HG003 strain without leader fusion was included as control. (C) Inhibition of nitrite production by IsrR. HG003 strain and its Δ*hfq* derivative harboring a control plasmid (pCont; pRMC2ΔR) or a plasmid expressing IsrR (p-IsrR; pRMC2ΔR-*isrR*), were grown in rich media under anaerobic conditions. Nitrate ($NaNO_3$) was added to the media and 150 min after, the nitrite produced was compared qualitatively using that Griess colorimetric method. Expression of IsrR inhibited nitrite production in the Δ*hfq* strain as observed in HG003.
